# cyto7: an open, vertex-level cortical type atlas of the human cortex

**DOI:** 10.64898/2026.10.01.755866

**Authors:** R. Salvador, B. Mercadal, F. Castaldo, M. Á. García-Cabezas, G. Ruffini

**Affiliations:** Research Department, Neuroelectrics Barcelona, Barcelona, Spain; Barcelona Computational Foundation, Barcelona, Spain; Department of Anatomy, Histology and Neuroscience, School of Medicine, Universidad Autónoma de Madrid, Madrid, Spain; Neural Systems Laboratory, Department of Health Sciences, Boston University, Boston, MA, USA

## Abstract

The cerebral cortex is organized along a gradient of laminar complexity, indexed principally by the emergence and granularity of layer IV and by the size of layer III neuron somata, and cortical types are the ordered categories along it. The Structural Model builds on that gradient to predict laminar connectivity, plasticity and disease vulnerability ^1,2^. Existing digital cortical-type maps are area-level: they assign one type per classical area and cannot represent a type boundary that runs through an area. We release cyto7, an open, vertex-level, seven-type cortical-type atlas delineated on the fsaverage surface following the García-Cabezas protocol and revised by the protocol’s authors.

We distribute the atlas with a reusable quality-control and support framework. A topological audit reports the map against four spatial rules that follow from cortical type analysis and the dual origin of the neocortex theory: neighbouring vertices differ by at most one type, the least complex cortex forms one continuous limbic belt, eulaminate II forms a single matrix enclosing a few holes, and eulaminate III and koniocortex form isolated islands. These were imposed during construction rather than tested, and the audit reports what imposing them cost. A vertex-wise benchmark against the existing area-level map, one type per von Economo–Koskinas area, shows exact agreement at 52.5% of vertices, with disagreement almost always a single level and concentrated inside the large, heterogeneous areas that a single areal label cannot resolve. Where the two maps disagree, cyto7’s label better matches the independent multimodal profile in 60.1% of pooled feature-by-vertex comparisons, against 46% under a rotation null and 54% under granularity-matched alternative partitions (*p* = 0.020), the harder of the two. A per-vertex anatomical support score, computed from anatomy alone, states how strongly each hand-drawn label is backed by the anatomical evidence, and is deliberately not a calibrated error probability.

We then validate cyto7 against independent brain measures not used to define it, under a spatial-autocorrelation (spin) null with FDR control. Intracortical myelin (T1w/T2w) increases, cortical thickness decreases, and the principal functional-connectivity gradient reverses along the type axis (all *q* ≲ 0.006). Source-reconstructed resting-state MEG dynamics also track the axis: the intrinsic timescale shortens toward koniocortex (*ρ ≈ −*0.46) and survives the null, as do the spectral centroid and the slow/fast band-power ratio, whereas the aperiodic 1/f exponent does not. The laminar marker genes are where cyto7 differs most from a quantisation of a continuous map: it shows a larger absolute correlation with the layer-IV marker RORB (*ρ* = +0.78) and the layer-V marker FEZF2 (*−*0.59) than seven-level septiles of myelin, of the functional gradient or of a histological gradient, on four of five markers, while tying or losing to those septiles on the macroscale proxies. These are larger observed associations against three tested quantisations, not a test that the differences between maps are reliable, and they do not establish the accuracy of any individual boundary. Defined from anatomy alone, cyto7 recovers the sensorimotor-to-association hierarchy and, on these markers, some of the laminar signal underlying it. It is released with its full pipeline and an uncalibrated anatomical support score as a reusable substrate for structure–function and modelling studies.

## 1. Introduction

The human cerebral cortex is not a uniform sheet: its microstructure varies systematically across its extension. Cortical areas can be ordered into a small number of cortical types by their degree of laminar complexity, indexed principally by the emergence and granularity of layer IV and by the size of layer III neuron somata ^3^, from the three-to-four-layered allocortex through agranular and dysgranular limbic cortex and the eulaminate isocortices to the hyper-granular koniocortex. The Structural Model builds on that ordering ^2^, recently articulated as a modern neurodevelopmental synthesis ^4^: type predicts the laminar pattern of cortico-cortical connections ^1,2,5^, it tracks a plasticity and stability continuum ^6^, and it orders the differential vulnerability of areas to disease ^2^. The ordering has a developmental and evolutionary basis in the dual origin of the neocortex theory ^7–9^.

The cortical-type axis can be inferred non-invasively. The T1w/T2w ratio, a proxy for intracortical myelin ^10^, indexes anatomical hierarchy and the dominant axis of transcriptional variation ^11^; it converges with the principal gradient of functional connectivity ^12^. More generally, this is one of a family of macroscale gradients that run along a common sensorimotor-to-association (unimodal-transmodal) axis that jointly organize cortical features: cortical microstructure and myeloarchitecture, gene transcription, and evolutionary/developmental cortical expansion ^9,13,14^. Microstructural gradients derived from histology track laminar complexity and dissociate from function only in transmodal cortex ^15^.

Cortical dynamics form a parallel hierarchy. Intrinsic neural timescales lengthen from primary sensory toward association cortex ^16,17^, the dominant frequency of resting oscillations decreases along the same hierarchy ^18^, and this dynamical gradient is mechanistically tied to cortical microarchitecture: cell density, myelination and synaptic/ion-channel gene expression ^19^. Therefore, it should track cytoarchitectural type as well. Concretely, along the gradient of laminar complexity from limbic cortex to koniocortex we predict that the intrinsic timescale decreases (faster dynamics in more granular, more complex cortex) while a power-weighted spectral centroid, a coarse proxy for dominant frequency, increases; bandlimited power, being more redundant and spatially smooth, is a weaker test.

These convergences motivate a direct test: does a cytoarchitectural type map, defined independently of imaging, organize these modalities?

Answering this requires a type map on a standard surface. The digital cortical-types map in common use is area-level: the von Economo–Koskinas areal atlas ^20^ (digitized by Scholtens et al. ^21^) is relabelled with the cortical types of García-Cabezas et al. ^3^, yielding one (sometimes averaged) type per area. Such a map cannot represent a type transition that runs within an area, and the most prominent reconstruction of the classical atlases into MRI space ^22^ does not address this either. The cost of that gap is visible in current practice. To validate a signal-flow hierarchy against cortical type, Oh et al. ^23^ had to build a type map for the occasion, applying the annotations of García-Cabezas et al. ^3^ to the Brodmann atlas and projecting the result onto the 360-area multimodal parcellation. The transfer places 261 of 360 parcels in two types and leaves the entire mesocortex in 39, which is the resolution any area-level relabelling can reach. The relabelling also borrows the atlas only as a set of spatial parcels: the types are assigned to the areas post hoc from the protocol of García-Cabezas et al. ^3^, not read off von Economo and Koskinas’s own laminar classification, which follows Brodmann’s two-type (homogenetic/heterogenetic) scheme, has no separate mesocortex, and groups areas of disparate architecture and developmental origin together (for example classing some parahippocampal and retrosplenial areas with primary visual cortex as heterotypical granular; ^4^). Cortical type analysis instead treats the mesocortex, the agranular and dysgranular limbic belt, as a graded sector between allocortex and isocortex ^1,2,5^.

Here we release cyto7, a seven-type cortical atlas delineated vertex by vertex on the 164k fsaverage surface, and make three contributions. First, the atlas itself: allocortex-aware, openly licensed, and released in the surface formats the field already uses. It ships as FreeSurfer annotation files on fsaverage and as label files on the 32k fs_LR surface adopted by the Human Connectome Project, so it transfers to an individual reconstruction or into an existing surface pipeline without conversion. Second, a reusable quality-control and support framework for hand-drawn type maps: a topological consistency audit, a vertex-wise benchmark against the area-level map, and a per-vertex support estimate. Third, a multimodal validation of the anatomy-defined type axis against independent data: intracortical myelin (T1w/T2w), cortical thickness, the principal functional-connectivity gradient, MEG dynamics, a transcriptomic expression gradient (AHBA), and a population-averaged tractogram, with a direct histological check against single-specimen BigBrain reported as an open and inconclusive comparison. We also clarify what the fsaverage surface can and cannot represent (the archicortex is off-surface, and the allocortical dual-origin “double loop” closes only in the three-dimensional mantle, even though the agranular and dysgranular type rings themselves are recovered on the surface).

### Box 1: The cortical types

Cortical areas are ordered along a single *gradient of laminar complexity*, indexed by two cytoarchitectonic criteria: principally the emergence and granularity of layer IV, and alongside it the size of the largest pyramidal somata in layer III (after García-Cabezas et al. ^3^, Table 2; ordinal codes *τ* = 1… 7 used throughout). The two do not run in step at the granular end. Layer III pyramids enlarge steadily from allocortex to eulaminate III, a progression Sanides named externopyramidization, but koniocortex departs from it: its superficial layers are hypercellular with small, unprominent neurons. Layer IV granularity continues to rise where layer III soma size does not. These seven types group into three tiers: **allocortex** (type 1), **mesocortex** (agranular–dysgranular, types 2–3) and **isocortex** (eulaminate, types 4–7).

**1 - Allocortex.** Three to four layers; no granular layer IV. Comprises olfactory allocortex (paleocortex: piriform cortex and anterior olfactory nucleus), hippocampal allocortex sensu stricto (fascia dentata, cornu ammonis, subiculum), and modified hippocampal allocortex (presubiculum, parasubiculum, entorhinal cortex). On the fsaverage surface the olfactory paleocortex (piriform) and the modified hippocampal allocortex (entorhinal, pre-/parasubiculum) are present; the hippocampal allocortex sensu stricto is off the surface, in the volume.

**2 - Agranular.** Five recognizable strata; layer IV absent; the largest pyramids lie in layer V (e.g. the agranular sectors of the anterior cingulate, the rostral insula and the temporopolar cortex).

**3 - Dysgranular.** An incipient, discontinuous layer IV; the largest pyramids still lie in layer V (e.g. the dysgranular sectors of the cingulate, the mid-insula, the temporopolar cortex and the rhinal/parahippocampal transition). Agranular and dysgranular sectors coexist within every mesocortical region, so no such region is of a single type as a whole.

**4 - Eulaminate I.** Six well-defined layers with a thin-to-moderate layer IV; large pyramids in both layer III and layer V.

**5 - Eulaminate II.** Six layers with a clearly developed layer IV, larger neuron bodies in layer III than in layer V, sublayers in V–VI and sharper laminar borders.

**6 - Eulaminate III.** Dense, sharply defined lamination with a prominent layer IV and quite large neuron bodies in layer III, larger and more prominent than in eulaminate II. Includes primary motor cortex (FA/M1): despite a thin layer IV, its high laminar complexity places it here, following the principle that “in motor cortex, less is more” ^3^.

**7 - Koniocortex.** Maximal granularity: a dense, wide layer IV and hypercellular layers II–III, whose neurons are small and unprominent rather than enlarged (primary sensory cortices V1, S1/area 3b, A1).

*Note. The gradient orders degrees of laminar complexity. It varies smoothly between neighbouring vertices, which is what rule R1 enforces, but it is not a single ramp across the sheet: the same type recurs in disconnected locations, and types 1 to 5 form nested concentric rings while only eulaminate III and koniocortex form islands (§3.2). A single classical area may span more than one type, which is precisely what a vertex-level map can represent*.

## 2. Methods

### Ethics

This study analysed previously collected, de-identified data obtained under the data-use terms of their original providers: group-average template maps, and individual-level Human Connectome Project young-adult data (resting-state MEG, *N* = 89; T1w/T2w myelin maps, *N* = 200) accessed under the HCP Open Access Data Use Terms. No new human or animal data were collected. The original studies obtained ethics approval and participant consent, and no additional approval was required for this secondary analysis.

### 2.1 Atlas construction and expert reconciliation

Seven cortical types (allocortex, agranular, dysgranular, eulaminate I–III, koniocortex) were painted vertex-wise on the standard 164k-vertex fsaverage surface ^24^ using a purpose-built interactive tool, following the cortical-type criteria of García-Cabezas et al. ^3^. The initial hand-painting was then reconciled with the topological constraints of the gradient of laminar complexity and revised by the protocol’s authors; the released map is therefore hand-painted and systematically corrected, not a raw drawing. Type definitions are reproduced in Box 1 (after García-Cabezas et al. ^2^). For the limbic belt, labels were adjudicated against a reliability-weighted anatomical reference combining FreeSurfer ex-vivo histological labels ^25,26^, the Glasser HCP-MMP parcellation ^27^ and the gyral (Desikan/Destrieux) atlases, ordered by how directly each reflects cytoarchitecture (ex-vivo histology the most reliable, gyral atlases the least). Where sources disagree, the more reliable prevails: an allocortex-versus-isocortex call requires histological or Glasser support, the gyral atlases able to corroborate but not override (formalized in Annex A–B). Two allocortical subfields have no direct surface criterion: the presubiculum and parasubiculum are delineated by a geometric proxy on the template mesh (the parahippocampal band within three mesh hops of the medial wall, minus entorhinal cortex), which sets the extent of the surface allocortex and is a template rule rather than a cytoarchitectural or developmental one. The construction and reconciliation workflow, including the change log relative to the initial painting, is described in Supplementary Annex A; the change log ships with the data release as provenance.

Three choices in how the map was drawn are worth mentioning explicitly, because they affect how it should be read.

#### Criteria and anatomy

Each vertex was typed by the cytoarchitectonic criteria of García-Cabezas et al. ^3^ (Box 1): principally how well developed the granular layer IV is, from absent (allocortex and agranular cortex) to hyper-developed (koniocortex), together with the size of the largest layer III pyramids, which enlarge from allocortex to eulaminate III and then give way to the small, hypercellular superficial layers of koniocortex (Box 1). The usual anatomical landmarks (gyri, sulci and known areal locations) were used as reference. The topological rules (the sequential type gradient and the ring/island structure) were treated as constraints that the finished map must satisfy, not as the drawing rule: boundaries were placed by cytoarchitecture and anatomy first, and only then checked and minimally repaired for topology.

#### Left–right symmetry was not enforced

The same criteria and reference hierarchy were applied independently to each hemisphere by the same rater, so genuine interhemispheric differences are preserved rather than averaged away; we instead report left–right agreement as a quality-control measure (Supplementary Annex A).

#### No boundary smoothing was applied

The released labels are kept at the native vertex resolution, so each boundary follows the hand-drawn cytoarchitectural judgement exactly rather than being idealised into a smooth curve. This keeps the map faithful to the evidence at the cost of some jaggedness where types meet. A boundary-smoothed variant (a topology-preserving majority-vote relaxation) is provided for display only; we do not release it as the map, because smoothing that is not carefully guarded erodes the smallest features, notably the koniocortex islands of V1, A1 and S1.

### 2.2 Topological quality control

Cortical type analysis and the dual origin of the neocortex theory predict a specific spatial arrangement of cortical type: type varies gradually across the sheet, the least complex cortex forms a peripheral allocortical ring, successive rings of increasingly complex tissue are nested inside it, and the most complex tissue appears as isolated koniocortical islands at the centre ^2,4,7–9,28^. We translate these qualitative predictions into four objective, machine-checkable topological rules on the surface graph (vertices connected by mesh edges), and validate the map against them (Supplementary Annex A):

#### R1 - sequential gradients

adjacent vertices differ by at most one cortical type, | *τ* (*i*) *− τ* (*j*) | *≤* 1 for every edge (*i, j*), as the gradual transitions along the gradient of laminar complexity require^3,4^;

#### R2 - core continuity

the least complex cortex forms one continuous limbic (allo- and mesocortex) belt rather than disconnected patches (a belt-level criterion specified a priori, not one closed ring per grade). Operationally, the union of allocortex, agranular and dysgranular vertices is a single connected component encircling the medial wall (an annulus separating it from isocortex), reflecting the concentric, continuous gradation of types from the allocortical ring through the mesocortex (Filimonoff’s cortex intermedius) to the isocortex (Sanides’ concentric rings; ^2,4,7–9)^. In the released map, each grade is moreover individually continuous (a single connected component): the agranular and dysgranular belts close into complete rings on the surface (first Betti number *β*_1_ = 1), whereas the allocortex forms a single open arc (*β*_1_ = 0). This is expected, as the allocortical ring does not close on the cortical surface but only in the three-dimensional brain, through the hippocampal formation (the hippocampus sensu stricto and indusium griseum), which lies off the fsaverage surface (§3.2, §4);

#### R3 - the eulaminate-II matrix

must be a single continuous matrix enclosing a small number of specialized-area holes (three in the released map), so that eulaminate II is a ring like the types outside it rather than an island;

#### R4 - higher-order islands

Eulaminate III and Koniocortex must exist as isolated clusters (islands) rather than continuous rings.

Only objective, topology-restoring repairs (isolated speckle removal, buffer insertion) were applied. Larger reassignments were only performed if flagged by the expert reviewer.

### 2.3 Benchmark against the area-level map

To quantify the benefits of a vertex-level map, we compared cyto7 to the area-level type map: the digitized von Economo–Koskinas areal atlas ^21^ with each area assigned a type from García-Cabezas et al. ^3^. Both maps were placed on a common mesh and compared vertex-by-vertex with a confusion matrix, Cohen’s *κ*, a quadratically-weighted *κ_w_* (appropriate for the ordinal type scale), the adjusted Rand index (ARI), and a per-vertex difference map (Supplementary Annex B). Where the two maps disagree, we further assessed which label better fits the independent multimodal profile: for each of the three axis features (T1w/T2w myelin, cortical thickness and the principal functional gradient) we computed the median feature value of every isocortical type under each map’s own labels, and at each disagreement vertex assigned the win to whichever map placed the vertex closer to the median of the type it had assigned, pooling these per-feature outcomes into a win-fraction over the disagreement set. Because neighbouring vertices are strongly autocorrelated and the chance win-rate is not 50%, we assessed this fraction against a spin null (rotating the cyto7 labels, rebuilding the per-type medians and recomputing the win-fraction), which sets the chance baseline (*≈*46%) and the spin *p* (§3.3).

### 2.4 Per-vertex anatomical support score

We estimate, for each labelled vertex *v*, an **anatomy-only** support score supp(*v*) *∈* [0, 1] as a weighted geometric mean of four components (Supplementary Annex C, summarized in Box 2),

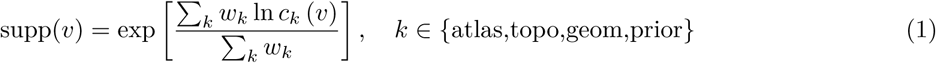

combining three terms read from the map and its geometry, namely atlas concordance (ordinal agreement with the von Economo–derived areal type map), topological consistency (down-weighting vertices near any topological-rule violation) and geometric/medial-wall reliability (lower near the medial wall and near type borders), together with a per-type provenance prior, an a priori estimate of how securely each type can be adjudicated (higher for the eulaminate isocortex and koniocortex, lower for the limbic belt where typing is least certain). Functional and microstructural data are deliberately excluded, so the score stays independent of the modalities used to validate the atlas. We further asked whether the support score could be improved by additional components, under a rule fixed in advance: a candidate is adopted only if it raises the map’s ability to predict *where* cyto7 and the area-level reference disagree (the *benchmark-independent disagreement AUC*, i.e. how well the support score, using only components that do not depend on that benchmark, localizes the cyto7-vs-areal disagreement), and only if it is not redundant with an existing component. Three candidates were tested (local label heterogeneity, cortical curvature, and a borrowed cross-subject variability prior from the Jülich probabilistic cytoarchitectonic atlas; ^29^); the outcome is reported in Annex C.

#### Box 2: The per-vertex anatomical support score

For every labelled vertex *v* we estimate a support score supp(*v*) ∈ [0, 1] as a weighted geometric mean of four **anatomy-only** components (Equation 1).

##### Why a geometric mean

It penalises a low component more than an arithmetic mean would, so a weak term pulls the score down rather than being averaged away. It is a multiplicative penalty, not a veto: with equal weights, components of 1, 0.5, 1 and 1 still give 0.841, above the 0.66 high-support threshold, so a topological penalty can coexist with high overall support. A component of exactly zero would send the logarithm to negative infinity, so components are floored at a small positive value; the floor is given in Annex C.

##### Why anatomy-only

T1w/T2w and functional data are deliberately excluded, so the score stays independent of the modalities used to validate the atlas (no circularity); a separate descriptive myelin-agreement overlay never enters the score.

##### Outputs

A continuous map plus a categorical high / medium / low map (≥ 0.66 / 0.33–0.66 / < 0.33) for visualization, and per-vertex weights that downstream analyses can use to down-weight or mask uncertain tissue.

##### Behaviour

Eulaminate and koniocortex vertices score highest (expert-vetted, well inside area interiors); the small, near-medial-wall piriform allocortex sliver scores lowest. Full component definitions are in Supplementary Annex C.

### 2.5 Multimodal data and analyses

We validated cyto7 against independent brain measures, none of which was used to paint it. All maps were brought to the 32k fs_LR surface, except the per-subject MEG measures, which are computed and tested on the native 4k fs_LR source mesh (§2.6) ^30,31^ and tested with the same spin+FDR pipeline, with FDR controlled separately within each measurement family (families defined in §2.6); each measure is summarised in Box 3. The cyto7 map itself was resampled from 164k fsaverage to 32k fs_LR by label-appropriate resampling using the standard registration spheres (Supplementary Annex D).

#### Structure–function

Group-average maps of T1w/T2w myelin (bias-corrected) and cortical thickness from the HCP Q1–Q6 RelatedValidation210 group average (N = 210, MSMAll-registered, 32k fs_LR; ^27^), the group against whose mesh the cyto7 labels were resampled; the individual T1w/T2w maps of 200 HCP young-adult subjects (S1200 release, MSMAll-registered, 32k fs_LR; ^31^), used to test how the map generalises across individuals (§4.8); the principal functional-connectivity gradient^12^, accessed through neuromaps ^32^; resting-state MEG dynamics (intrinsic timescale, band powers, spectral centroid, the slow/fast band-power ratio and the aperiodic 1/f exponent) computed from the HCP young-adult MEG (N = 89), source-reconstructed with a FieldTrip realistic single-shell (FieldTrip 20200607, MATLAB R2018a) and fitted per subject with specparam, and validated against the corresponding neuromaps group maps (^33^; Annex F); and a population-averaged tractogram (HCP-1065; ^34^) analysed with DIPY ^35^. From these we fit the myelin progression across types (Annex E), summarised the nine structure–function features per type as a fingerprint, derived single-metric frequency summaries and a power-weighted spectral centroid (Annex F), and tested type-distance connectivity in the tractogram (Annex G).

#### Molecular and receptor

The first principal component of Allen Human Brain Atlas cortical gene expression (via abagen; ^36^); and the 19 PET receptor and transporter maps of Hansen et al. ^37^, analysed both as a single principal component and map by map, grouped by pharmacological class into ionotropic and metabotropic composites and their contrast, and summarised as a per-vertex receptor-diversity index (the Shannon entropy of the 19 normalised receptor densities at each vertex; Annex H).

#### Evolutionary, developmental and histological checks

Evolutionary and developmental cortical expansion ( ^38,39^, via neuromaps). These were tested against type and their values are reported in Table S8, but we do not build on them, for the reason given in §3.7. As a direct histological check, single-specimen intracortical staining-intensity profiles from BigBrain ( ^40^; via BigBrainWarp, ^41^). Two separate BigBrain analyses follow from these profiles. The first validates the profile itself: to guard against a spurious result from a single specimen, a pre-specified index, profile skewness, had to track type, and it did not (|*ρ*| 0.209 against a 0.4 threshold), so that gate failed. The second is the added-value test, which used the microstructural-profile-covariance gradient (Hist-G1) and was run (§3.3, Annex H).

#### Disease vulnerability

Eight ENIGMA case–control cortical-thickness maps (Cohen’s *d*, the standardized case-minus-control difference in cortical thickness; ^42^), the most widely available cross-disorder structural phenotype, correlated at Desikan–Killiany parcel resolution with a continuous per-parcel cyto7 type score, with a partial correlation controlling for the principal functional gradient (§3.7, Annex H).

Structure–function results appear in §3.5 (microstructure) and §3.6 (dynamics), and the molecular and receptor alignment with the gradient of laminar complexity (gene PC1, receptor composites, cortical expansion and disease vulnerability) in §3.7. Figure 5 is the scorecard for all of them: each headline measure with its direction, its effect size, and whether it survives the spin null with FDR control. A connectivity analysis of the population-averaged tractogram is reported in Annex G rather than here: short-range streamline counts are highest between cytoarchitecturally similar types, but the falloff does not survive a spin null once expressed per available endpoint pair, is absent at the level of the template bundles, and cannot separate cytoarchitectural from physical distance, so no inferential claim is drawn from it (§4.9). The histological (BigBrain) check is inconclusive in a single specimen and is taken up as a limitation in §4.9 (Annex H).

##### Box 3: The multimodal validation measures

The atlas is validated against independent brain measures, none of which was used to paint it. Each indexes the same sensorimotor-to-association hierarchy that cytoarchitectural type is expected to track, so we note for each the direction predicted along the gradient of laminar complexity (allocortex *τ* = 1 → koniocortex *τ* = 7).

###### T1w/T2w ratio (intracortical myelin)

The ratio of a T1- to a T2-weighted MR image, a widely used non-invasive proxy for how heavily myelinated the cortex is ^10^. High in primary sensory cortex, low in limbic/association cortex, predicted to *increase* with type.

###### Cortical thickness

The width of the cortical ribbon (mm). Thin in highly complex primary sensory cortex, thick in agranular/association cortex, predicted to *decrease* with type.

###### MEG band power (*δ, θ, α, β, γ*)

The strength of resting-state magnetoencephalographic oscillations in each canonical frequency band: which rhythms dominate where. Of the five bands, aperiodic-corrected *β* power tracks type once the 1/f background is removed; *δ* and *γ* are not estimable from a 2 to 40 Hz spectral fit and are reported as such rather than as nulls (§3.6).

###### Intrinsic (neural) timescale

How long the local signal stays correlated with its own recent past, i.e. how fast or slow a region’s dynamics are ^33^. Short (fast) in primary sensory cortex, long (slow) in association/limbic cortex, predicted to *decrease* with type.

###### Spectral centroid

The power-weighted mean resting frequency at each vertex: a single-number summary of “how fast” the local rhythm is. Higher (faster) toward more complex cortex, predicted to *increase* with type.

###### Aperiodic 1/f exponent

The slope of the non-oscillatory background of the power spectrum, an indirect correlate of local excitation–inhibition physiology ^43^: steeper means more inhibition-dominated. Tested but, unlike the timescale, no association with type was detected under its primary specification (§3.6).

###### Principal functional-connectivity gradient

*Not* a spatial gradient across the surface, but a coordinate derived from the resting-state functional-connectivity matrix (how each region’s time-course correlates with every other’s). A dimensionality-reduction step (diffusion-map embedding; ^12^) compresses each vertex’s whole connectivity profile to a single value, so vertices with similar connectivity get similar values. Its dominant axis separates unimodal (primary sensory/motor) from transmodal (default-mode/association) cortex, recovering the cortical hierarchy from function alone. Because koniocortex sits at the unimodal end and the limbic types at the transmodal end, it is predicted to run opposite to myelin (*decreasing* with type).

###### Template tractogram (HCP-1065)

A population-averaged whole-brain set of white-matter streamlines reconstructed from diffusion MRI ^34,35^, used to count short-range cortico-cortical connections between type pairs and test whether connectivity declines with cytoarchitectural type-distance.

###### Gene expression PC1 (AHBA)

First principal component of cortical transcription (5 of the 6 source donors; Annex H), the dominant molecular gradient. Predicted to *increase* with type; convergent with, not independent of, the imaging gradient.

###### Layer-marker genes (AHBA)

Cortical expression of canonical laminar markers from the same Allen atlas, read per layer at Desikan-parcel resolution: RORB (layer IV), CUX2 (upper layers), and FEZF2/FOXP2/TLE4 (deep layers). RORB directly tests the layer-IV criterion and is predicted to *increase* with type; the layer-V marker FEZF2 *decreases*.

###### Receptor PC1 (PET, 19 tracers)

First PC of neurotransmitter-receptor densities. A molecular cross-check; reported as context.

###### BigBrain intensity profiles

Single-specimen 20-µm histological depth profiles; their shape encodes lamination. Used for a direct histological check (Annex H).

*Note. Because these measures are independent of the anatomy used to draw cyto7, agreement is convergence with independent data rather than a re-description. It is not cross-validation, since nothing is held out and refitted. All but the per-subject MEG dynamics are group-average template maps; how they vary across individuals is discussed in §4*.

### 2.6 Statistics

For each feature we computed the Spearman correlation *ρ* between the per-vertex value and the ordinal cortical type. Because neighbouring vertices are strongly autocorrelated, so naive *p*-values are grossly anticonservative, significance was assessed with a **spin test**^44^. As part of its standard inter-subject registration FreeSurfer inflates each cortical hemisphere onto a sphere; the spin test re-uses that same sphere, applying a random rotation that slides the whole type map to a new position while preserving its spatial structure: the relative size of each type and the way neighbouring vertices cluster. Rotating the map *N* = 1000 times builds a spatially-constrained null distribution of correlations, and the spin *p*-value is the fraction of rotations whose (absolute) null correlation matches or exceeds the observed one,

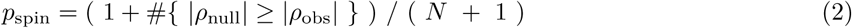

and p-values across the feature panel were adjusted with a Benjamini–Hochberg FDR^45^ (spin-null and FDR details in Supplementary Annex D, which lists all 21 families, their sizes and whether correction was applied within each, in Table S4). Table S8 lists every test in those families; robustness and sensitivity analyses reported in the text sit outside them and are listed, without correction, in Table S9. The MEG measures are computed on the native 4k fs_LR source mesh (8,004 vertices), to which the type map and every spin rotation are downsampled; the 32k fs_LR surface is used for the other modalities and for display. Unless stated otherwise, reported correlations are over all seven types; where a fit excludes allocortex, or allocortex and agranular, that is stated with the value, and both arms are given in the robustness listing (Table S9). The exclusions were motivated for the myelin regression in particular (it is under-sampled by surface maps, and its myelin does not follow the sensory progression that the gradient of laminar complexity captures). For the disease-vulnerability maps, which are defined on the Desikan–Killiany atlas rather than at vertex resolution, the null was a parcel-based spin (a rotation of the parcellated type score) rather than the vertex-level rotation. Single-metric additions, the slow/fast band-power ratio (§3.6) and the receptor-diversity index (§3.7), were each committed in advance as one test rather than selected from several candidates.

## 3. Results

### 3.1 The atlas

cyto7 provides a per-vertex, seven-type map of the human cortex on fsaverage (Fig. 1). Each vertex is assigned a cortical type by the granularity of layer IV together with the size of layer III neuron somata, the gradient of laminar complexity on which cortical type analysis rests ^3^ (Box 1): koniocortex (*τ* = 7) marks the primary sensory cortices with a hyper-developed granular layer IV, in the calcarine visual cortex (V1), Heschl’s auditory cortex (A1) and the fundus and posterior wall of the central sulcus (primary somatosensory cortex, area 3b); the eulaminate isocortices (*τ* = 4–6) carry a full six-layer pattern whose layer IV weakens progressively from the eulaminate-III collar around each koniocortical core, in occipital, perisylvian and ventrolateral prefrontal cortex, through parietal, temporal and prefrontal association cortex toward the limbic border; and the agranular and dysgranular mesocortex (*τ* = 2–3) forms the cingulate, insular, posterior orbitofrontal, parahippocampal, perirhinal and temporopolar belt, each of those regions carrying both an agranular and a dysgranular sector. Across the isocortex these assignments follow the typing of von Economo–Koskinas areas by García-Cabezas et al. ^3^, refined to vertex resolution and constrained so that neighbouring vertices never skip a type (rule R1; Methods §2.1). The surface allocortex (*τ* = 1) comprises the olfactory (piriform) paleocortex at the ventral fronto-temporal junction together with the modified hippocampal allocortex (entorhinal cortex and the pre-/parasubiculum) on the medial temporal lobe (*≈* 7,340 vertices; LH 4,147 / RH 3,193; delineated on developmental grounds, §3.4); the hippocampus sensu stricto (fascia dentata, cornu ammonis, subiculum) remains off the cortical surface, in the volume (§3.2, §4). Surface allocortex is somewhat more extensive on the left (2.8% vs 2.2% of the hemispheric surface); this follows the larger left entorhinal/medial-temporal parcel on the fsaverage template and the hop-distance proxy applied to it (§2.1), and is not offered as a biological finding (both hemispheres’ entorhinal cortex is essentially fully allocortical), and, as we do not enforce left–right symmetry, the per-vertex support score flags these developmentally-defined vertices (§3.4). Bringing the initial hand-painting into agreement with the topological constraints of the gradient of laminar complexity (the reconciliation step, Methods §2.1) changed labels only within this medial/temporal limbic belt, leaving the eulaminate and koniocortex isocortex as first painted (Supplementary Annex A).

**Figure 1.**
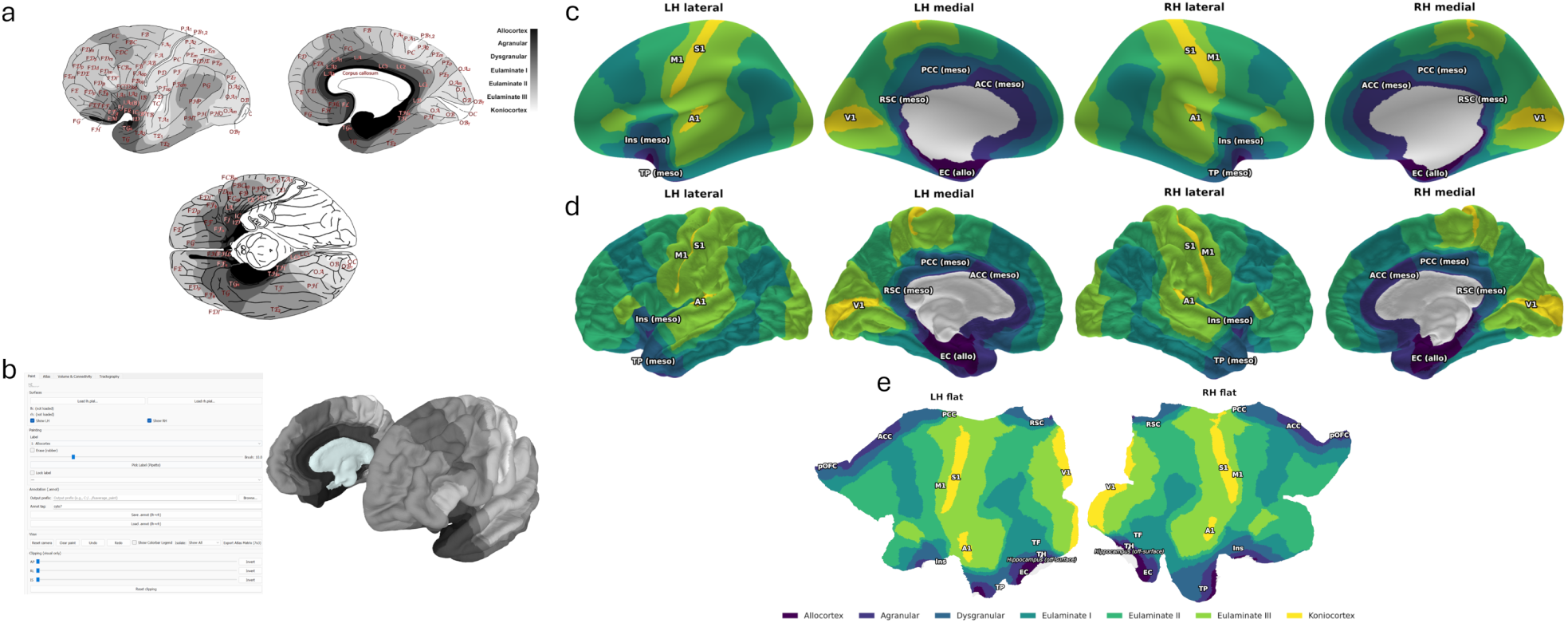
The cyto7 seven-type cytoarchitectural atlas on the cortical surface. *a*. The areal cortical-type reference scheme the labels follow (lateral, medial and ventral views), adapted from the grey-scale cortical-type map of García-Cabezas et al. ^3^ (licensed under CC BY 4.0); areas are shaded by cortical type from allocortex (dark) to koniocortex (light). *b*. The surface-native painting interface used to draw and revise the atlas directly on the fsaverage mesh. *c*. The cyto7 map on the inflated fsaverage surface (164k vertices), both hemispheres in lateral and medial views. Each vertex is coloured on a scale ordered by degree of laminar complexity, from allocortex (dark purple) through agranular, dysgranular and eulaminate I–III to koniocortex (yellow); the medial wall is masked (grey). For orientation, key areas are labelled: the koniocortical primary sensory areas V1 (visual), A1 (auditory) and S1 (somatosensory), and primary motor cortex M1, together with representative mesocortical areas (ACC/PCC, anterior/posterior cingulate; Ins, insula; TP, temporal pole), allocortex (EC, entorhinal) and retrosplenial cortex (RSC), which the map types as mesocortex and which is discussed as a borderline case in §4.9. *d*. The same on the pial surface (164k fsaverage). *e*. The same on a flat 32k fs_LR surface (both hemispheres). At that coarser resolution a thin (*∼*1-vertex) eulaminate-II buffer can be dropped, so a few type boundaries (e.g. at the insula–inferior-frontal junction) may appear to abut across a step; the released 164k map is topologically clean (rule R1, zero skip edges), as seen in the inflated views.

### 3.2 Topology and the concentric cortical-type rings

The map satisfies the sequential-gradient rule R1 with no violations after objective repair, and rules R2–R4 hold. In the released map the two low-complexity mesocortical limbic types (agranular and dysgranular) each form a single connected component that encircles the medial wall. Read as a flat map, with the medial wall at the perimeter, the arrangement is a set of nested rings: the allocortex outermost, the agranular ring inside it, the dysgranular ring inside that, eulaminate I forming a third concentric ring inside those, and the eulaminate-II matrix innermost, with the eulaminate III and koniocortex islands enclosed within it (R2–R4). This is the concentric topology of Puelles et al. ^28^ and of Fig. 2D of Zaldívar-Díez and García-Cabezas ^4^, a peripheral allocortical ring with a central isocortical island, recovered here directly on the cut fsaverage surface rather than only in the three-dimensional mantle; the residual dual-origin “double loop” of the allocortex still closes off-surface through the hippocampal formation, a point we return to in §4. Anatomically, each ring is continuous rather than a set of regional patches, and each mesocortical region contributes both an agranular and a dysgranular sector: the agranular ring runs without interruption through the agranular sectors of the anterior cingulate, the temporopolar cortex and the ventro-anterior (frontoinsular) insula, the dysgranular ring inside it runs through the dysgranular sectors of the posterior and anterior cingulate, the mid-insula and the temporopolar cortex and on into the retrosplenial transition, and the eulaminate-I ring closes inside both; the insula thus follows the expected ventro-anterior-agranular to dorso-posterior-granular gradient. R1 has zero skip edges in both hemispheres; eulaminate II is a single component enclosing exactly three holes; and eulaminate III and koniocortex each form three non-annular islands, with every koniocortical core collared entirely by eulaminate III (full audit values, Table S2). The eulaminate-III islands can be named, and they are the same in both hemispheres: an occipital island enclosing V1, a central and perisylvian island enclosing both S1 and A1, and a ventrolateral prefrontal island in the pars triangularis of the inferior frontal gyrus that encloses no koniocortex at all. That third island corresponds to von Economo’s area FDT (*frontalis granularis triangularis*), which García-Cabezas et al. ^3^ likewise type as eulaminate III, so the vertex map and the source areal classification agree on a highly complex island that no primary sensory area anchors. Because the rules were enforced rather than merely tested, we audited what enforcing them cost. Vertices relabelled to satisfy the sequential-gradient rule number 2,368 (0.78% of labelled cortex, 960 mm^2^), concentrated in the peri-insular and superior-temporal belt (Supplementary Fig. S13; Annex A). The rule itself required far fewer: applying the anatomical relabels alone and then restoring R1 needs 42 vertices, and only 298 of the graded band carry a label the rule strictly demands, so the band is a drawn gradient rather than a minimal repair. Removing every such vertex together with a two-hop mesh margin leaves all four primary correlations intact (largest change Δ*ρ* = 0.011; Annex A). The as-painted map and the per-vertex change log ship with the release, so every relabel can be inspected.

**Figure 2.**
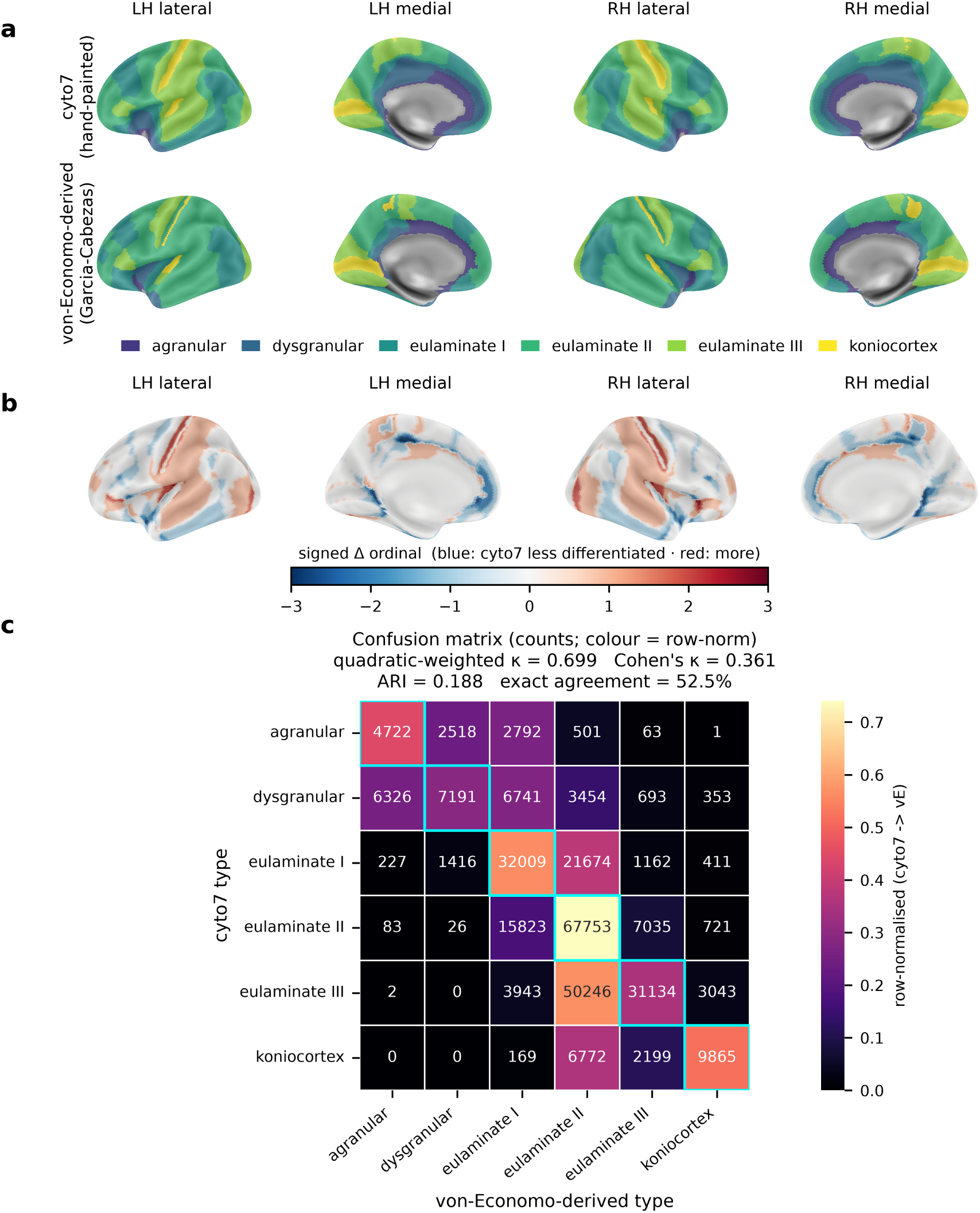
cyto7 refines the area-level cortical-type map at the vertex level. *a*. The two maps on the inflated fsaverage surface: the hand-painted cyto7 map (top) and the area-level reference (bottom), the digitised von Economo–Koskinas atlas ^21^ typed following García-Cabezas et al. ^3^. Colours denote the six non-allocortical types; allocortex and the medial wall (grey) lie outside this benchmark. *b*. Per-vertex signed ordinal difference (cyto7 *−* von-Economo): blue where cyto7 assigns a less complex type, red where more, white where they agree. Disagreement is overwhelmingly a single step, concentrated at areal borders and inside the largest, most heterogeneous areas (§3.3). *c*. Confusion matrix (rows, cyto7; columns, area-level type); cell numbers are vertex counts, colour is the row-normalised fraction, diagonal outlined. Agreement statistics are printed in the panel.

### 3.3 What a vertex-level map adds

Against the area-level map, cyto7 agrees at the type level for the majority of cortex, with disagreement overwhelmingly of a single ordinal step (Cohen’s *κ* = 0.36, quadratic-weighted *κ_w_* = 0.70, adjusted Rand index 0.19; 52.5% exact agreement). The near-doubling from the unweighted to the quadratic-weighted *κ* is itself informative: because the quadratic weighting discounts small ordinal disagreements, this rise confirms that most disagreements are a single type apart rather than scattered across the scale. The per-vertex difference map shows that this disagreement is not scattered uniformly, as random labelling noise would be, but is spatially structured, concentrating in two places: along the borders between von Economo areas, where a single sharp areal boundary meets the gradual vertex-level transition; and, more tellingly, inside the largest and most heterogeneous areas, where one areal label is stretched across tissue that the vertex map resolves into two neighbouring types (Fig. 2). Consistent with this, most classical parcels are cytoarchitecturally mixed: across both the Desikan–Killiany (34 parcels) and von Economo (43 parcels) atlases, the median parcel is only 67% a single cyto7 type and spans 2.4 (Desikan) and 2.0 (von Economo) types on average at *≥* 10% occupancy. Only the primary sensory and limbic anchors are near-pure (4/34 Desikan and 7/43 von Economo parcels are a single type at *≥* 90%, e.g. V1/pericalcarine koniocortex, entorhinal allocortex), whereas association and transitional cortex blend two or more types (Supplementary Table S5, Fig. S8; Annex I). The vertex map therefore recovers the within-area structure that any area-level label must discard.

Whether that refinement is an improvement can be tested, because the features that arbitrate it never entered the painting. On the 27,298 isocortex vertices where cyto7 and the area-level map disagree, cyto7’s label better matches the independent multimodal profile (myelin, thickness, functional gradient) in 60.1% of pooled feature-by-vertex comparisons (spin-test p = 0.001; chance *≈* 46%), increasing to 74.3% where the maps differ by *≥* 2 ordinal steps. The unit here is the comparison rather than the vertex: three arbiters across 27,298 vertices give 81,894 comparisons, which are not independent of one another. Scored instead as one decision per vertex, by majority over the three arbiters, the fraction is 62.8%. We report the pooled figure throughout, for comparability with the per-arbiter and comparator panels, and neither quantity is histological labelling accuracy. These three arbiters were fixed in advance (§2.3) as the axis features, and they are the three, of the nine tested, that track type at the vertex level under the spin null with FDR control (§3.5). Table S8 reports the same win fraction for the other six, which run from 0.48 to 0.64. The highest of all nine is among them: the intrinsic timescale.

Three checks help determine whether that advantage is a resolution artefact rather than an anatomical one. First, the chance baseline already prices resolution in: the spin null rotates cyto7 at its own vertex-level granularity and rebuilds the per-type medians from the rotated labels (§2.3), so a partition of cyto7’s shape earns 46% from geometry alone. Second, the type profiles are not fitted and tested on the same tissue: refitting the per-type medians on one hemisphere and scoring only the disagreement vertices of the other gives 59.7% and 60.5% in the two directions, against 60.1% in sample. Third, the advantage holds inside areas and not only at their edges: on the 4,844 disagreement vertices more than five mesh hops inside a von Economo area the win fraction is 58.7% (chance 44.8%, spin *p* = 0.007) against 60.4% in the boundary band, and it stays flat with distance from the border (59.8% at ten hops; Supplementary Fig. S12). What these establish is stronger alignment with the independent macroscale profile, and specifically that the alignment is not confined to areal borders; they do not establish that a given boundary is histologically correct, since the arbiters index the gradient of laminar complexity rather than lamination directly.

The one arbiter that does index cytoarchitecture directly favours neither map, and the two BigBrain analyses are separate. The added-value test was run first and used the microstructural-profile-covariance gradient (Hist-G1) as arbiter: on the disagreement set the win fraction is 0.4993 against a chance level of 0.519 (*p* = 0.740), a measured null a little below chance. Because Hist-G1 is a derived embedding that itself tracks type only weakly, we then specified in advance a check on the arbiter rather than on the map: a single differentiation index, profile skewness, was designated before that comparison was made and had to reach |*ρ*| *≥* 0.4 against type for BigBrain to serve as arbiter at all. It did not (|*ρ*| = 0.209, spin *p* = 0.321), so the gate failed and the profile comparison was not interpreted. One feature of the panel did survive the test (profile SD, *ρ* = +0.436, *q* = 0.015), but it indexes overall staining contrast rather than laminar differentiation, and we did not substitute it for the designated index; the index and the threshold were fixed in a written analysis plan, released with the code. The whole-cortex comparison favours cyto7 nominally without reaching significance (|*ρ*| 0.209 against 0.158, *p* = 0.343). Neither result establishes why it came out that way, and we do not attribute either to low power (§4.9). So the case that cyto7 improves on the area-level map rests on macroscale MRI proxies alone (§4.9), which is why the histological result sits here beside the 60.1% rather than in a limitations paragraph. The wins are also not ordered by anatomical support: across support tertiles they run 57.1%, 69.5% and 56.2% from low to high, the middle tertile some twelve points above both flanks. With three bins and no ordered trend we do not read a mechanism into that shape, but its consequence for downstream use is worth stating plainly: the support score does not rank vertices by how much cyto7 improves on an areal label, and using it as a weight for that purpose would down-weight the middle band, where the improvement is in fact largest.

Beyond the disagreement set, cyto7 also correlates more strongly than the area-level map with each continuous feature across the whole cortex (myelin |*ρ*| 0.61 vs 0.49; thickness 0.51 vs 0.41; functional gradient 0.58 vs 0.42; all q *≤* 0.006). Two further checks ask whether that comparison is fair. Both make it harder.

The first replaces the chance baseline. Rotating cyto7 rebuilds its per-type medians over anatomically arbitrary tissue, which is a weaker alternative than a plausible seven-class map. We therefore built 200 spatially contiguous partitions carrying cyto7’s own type-size distribution. These raise chance from 46% to 54.2% (s.d. 3.0%) and place the rotation baseline at the 0.5th percentile of their distribution, so the rotation null was too generous. cyto7’s 60.1% still clears the matched baseline, at its 98.5th percentile (*p* = 0.020), with the margin narrowed from fourteen points to six.

The second asks whether a quantisation of a single continuous map would do as well. Seven-level equal-count septiles of T1w/T2w myelin and of the principal functional gradient reach 0.703 and 0.714, above cyto7’s 0.601. But each is scored partly against the arbiter it was built from, at 0.909 and 0.927, and only that self-comparison is guaranteed to be high; how a septile performs on the other two arbiters is an empirical result. Because the pooled win fraction is exactly the mean over the three arbiters, each partition’s defining arbiter can be dropped exactly rather than approximately. Dropping it from both sides, so that the two maps are scored on the same features, gives cyto7 0.594 against the myelin septile’s 0.599, and 0.592 against the gradient septile’s 0.608. The gaps are 0.005 and 0.016 in the septiles’ favour. We have not estimated their uncertainty, so we read this as the benchmark failing to separate the maps rather than as either one winning. A septile of the BigBrain microstructural gradient, built from none of the three arbiters, reaches 0.502.

What this benchmark measures is therefore alignment with the macroscale hierarchy, which any map encoding that axis attains, and it cannot distinguish cyto7 from a quantisation of one continuous feature. The measures that do separate them are the laminar marker genes of §3.7, most sharply the layer-IV marker RORB and the layer-V marker FEZF2, where cyto7 shows a larger absolute correlation than any of the three septiles on four of the five markers, and than the histological septile on all five (Annex J). Those are larger observed associations on the same parcels, not a test that the differences between maps are reliable, and §4.10 sets out what would settle it.

### 3.4 Anatomical support

The support score states, from anatomy alone, how strongly the hand-drawn label at each vertex is supported by the atlas, geometric and provenance evidence (0 to 1, see Equation 1 and Box 2; Fig. 3). Its pattern mirrors how the map was built. The eulaminate isocortices and koniocortex (large, well-separated, expert-vetted types) are the best supported (median *≈* 0.9), and the agranular/dysgranular limbic belt is intermediate.

**Figure 3.**
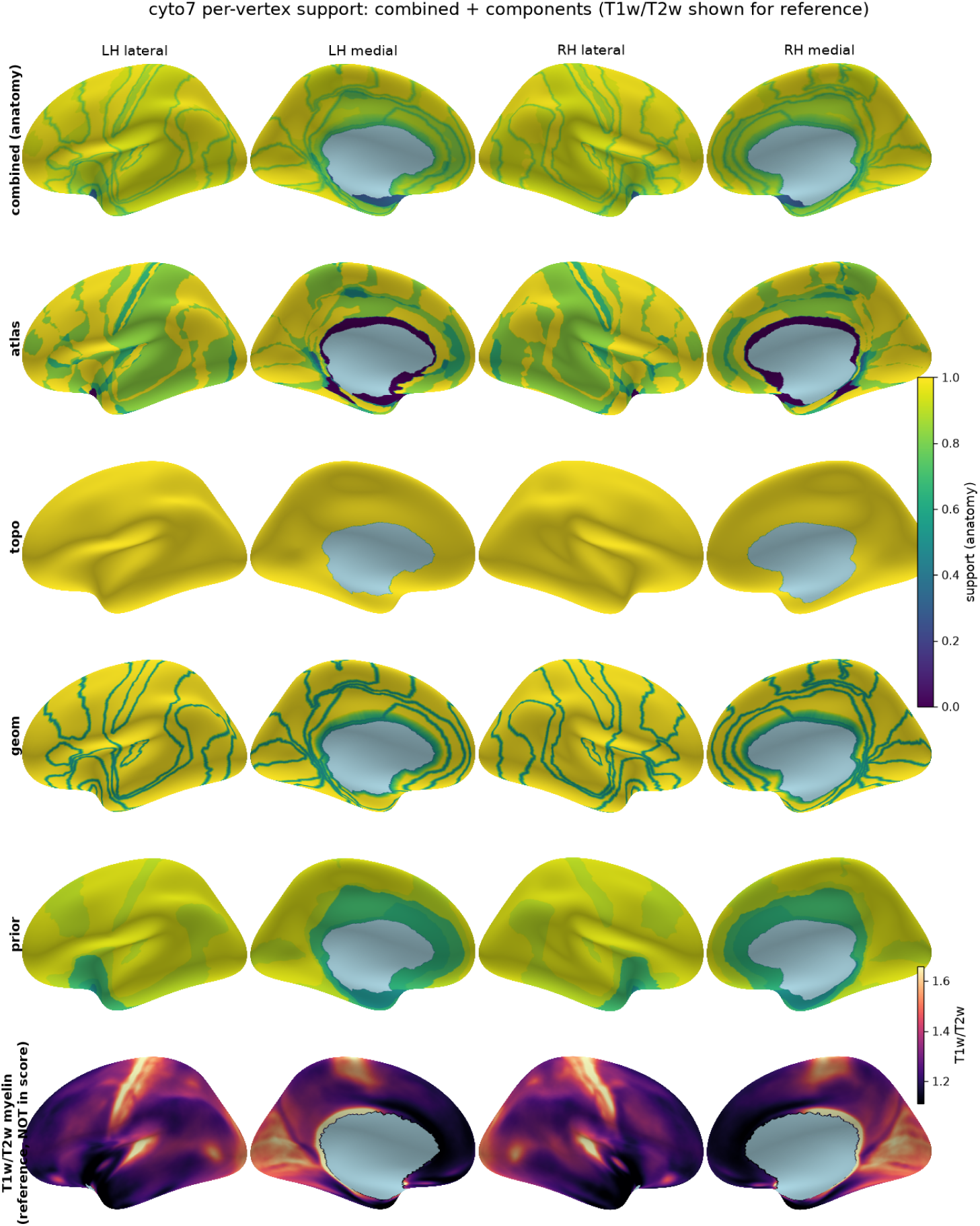
Per-vertex anatomical support score and its components. Inflated fsaverage surfaces (both hemispheres, lateral and medial views; medial wall masked in light blue). The combined map and its four components (top five rows) share a 0–1 support scale (viridis, upper colorbar); the bottom row uses a separate T1w/T2w scale (magma, lower colorbar). **Combined (anatomy):** the released support score, a weighted geometric mean of the four components below. It is highest over eulaminate and koniocortex, lowest at type borders, near the medial wall and over the developmentally-defined allocortex (piriform sliver least certain). Components: **atlas**, concordance with independent cortical-type atlases; **topo**, agreement with the sequential-gradient rule (uniformly high, as the map is topologically clean); **geom**, geometric reliability, decreasing towards type borders and the medial wall; **prior**, a per-type provenance prior. **Bottom row (T1w/T2w, reference, not in the score):** the observed myelin map on its own scale, shown for comparison and deliberately excluded from the anatomy-only support score (avoiding circularity with the structure–function validation); high in sensorimotor cortex, low in association/limbic cortex.

The allocortex, by contrast, is low-support (median *≈* 0.32): the surface allocortex (entorhinal cortex and the pre-/parasubiculum) is delineated on *developmental* grounds, since these areas arise from the same medial-pallial sector as the hippocampus ^28^, a kind of evidence that the atlas-concordance term of the support score cannot register. The low score is therefore the map flagging the tissue whose typing rests on expert developmental judgement rather than on the surface atlases. The index thus makes the evidence behind each label inspectable. Using it as an analysis weight is a heuristic sensitivity choice rather than inverse-error weighting: the scale is uncalibrated, and §3.3 shows it does not rank vertices by how much cyto7 improves on an areal label.

This score can be interpreted as a transparent, largely geometric reliability indicator rather than a finely calibrated error model, and its four components are not statistically independent of one another, the topology term being near constant by construction because the map was repaired until the rules held. It predicts only modestly where cyto7 and the area-level reference disagree, an ex-vivo histological patch of 509 belt vertices shows labelling error decreasing monotonically with the score, though the belt was partly built from those labels, so this is an internal consistency check rather than an external test, and the vertices are spatially clustered rather than 509 independent observations (0.15, 0.07 and 0.02 across tertiles, on 39 mislabelled vertices in total), and three candidate additional components were tested against the pre-specified rule of §2.4 and none qualified; these supporting analyses are in Annex C. One apparent contradiction in its construction is worth stating plainly. Atlas concordance is one of the four components and they carry equal weight, so agreement with the area-level reference contributes a quarter of the score, while §3.3 argues that many disagreements with that reference are legitimate within-area refinements. A vertex can therefore be scientifically informative precisely because it departs from the coarse map, and be scored slightly lower for doing so. The penalty is small in practice, for two reasons: the component is graded rather than binary, and the disagreement vertices at which cyto7 wins are not concentrated in the least-supported tissue (§3.3). A version of the score that excludes the atlas-concordance component can be rebuilt from the released per-component maps, which ship separately.

### 3.5 Structure–function validation

Independent imaging measures track the cyto7 axis (Fig. 4). For each of nine group-average structural and functional features (myelin, cortical thickness, the functional gradient, and the group-average neuromaps intrinsic-timescale and five band-power maps) we correlated its per-vertex value with the ordinal cortical type and tested the correlation against a spatial-autocorrelation (spin) null with FDR control. Three survive: intracortical T1w/T2w myelin increases with laminar complexity (*ρ ≈* +0.59, *q ≈* 0.004), whereas cortical thickness (*ρ ≈ −*0.50, *q ≈* 0.006) and the principal functional-connectivity gradient (*ρ ≈ −*0.58, *q ≈* 0.004) decrease (Fig. 4). The per-type box-plots make the progression explicit: myelin lowest in the agranular/dysgranular belt and highest in koniocortex, thickness and the functional-connectivity gradient running the other way. This is consistent with the canonical view in which primary sensory cortex is thin and heavily myelinated while limbic/association cortex is thick and lightly myelinated. These directions agree with independent reports that T1w/T2w indexes anatomical hierarchy ^11^ and that microstructural gradients track laminar complexity ^15^.

**Figure 4.**
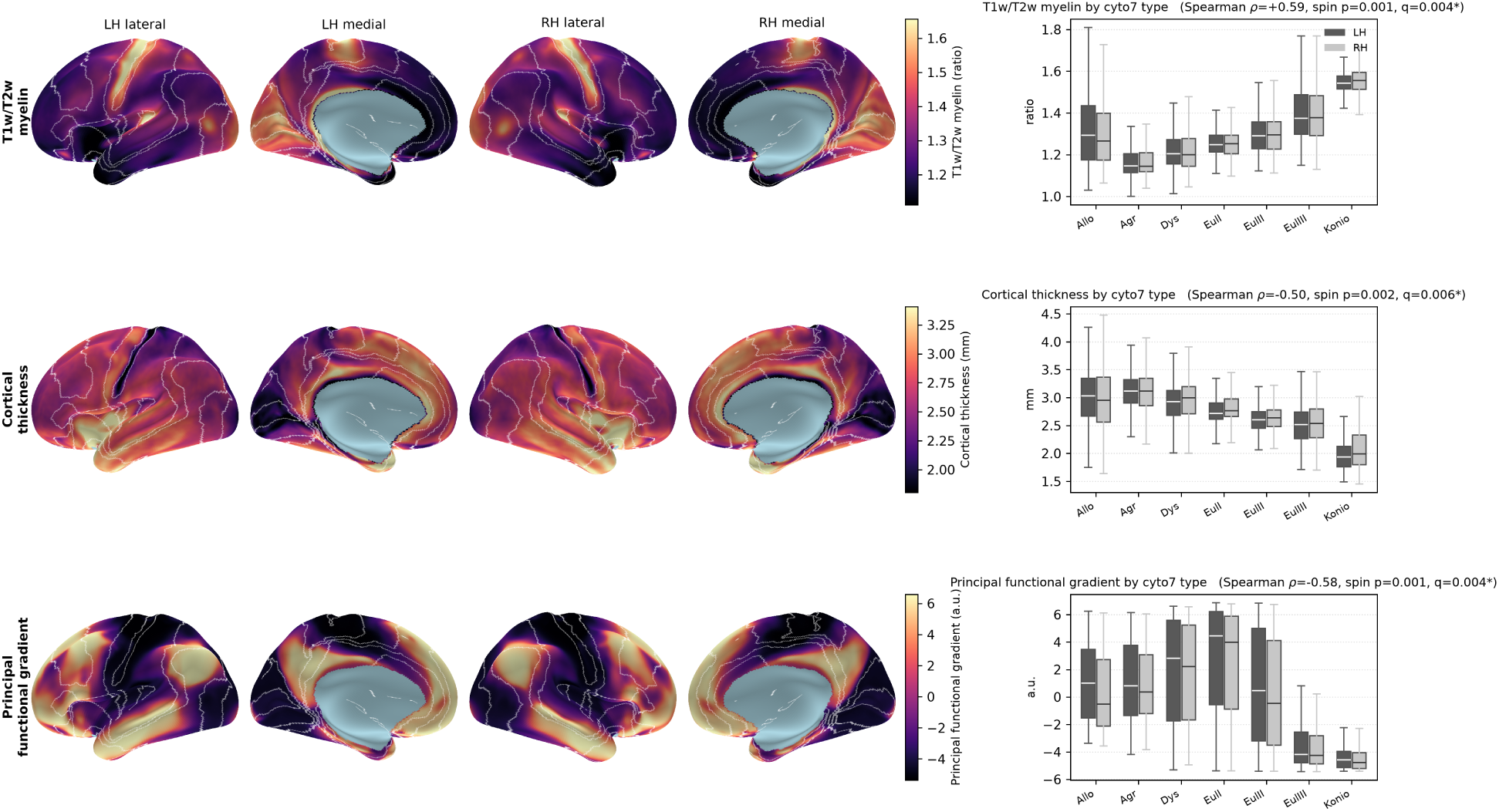
Structure–function validation: cortical features track the cyto7 gradient of laminar complexity. Each row shows one feature that survives the spatial-autocorrelation (spin) null with FDR control: T1w/T2w myelin, cortical thickness, and the principal functional-connectivity gradient. *Left:* the feature on the inflated 32k fs_LR surface (both hemispheres, lateral and medial views; magma scale, right colorbar; medial wall in light blue), with cyto7 type borders overlaid in white. *Right:* the feature’s distribution by cyto7 type (box-and-whisker; left hemisphere dark, right hemisphere light; boxes = IQR, whiskers = 1.5*×*IQR, outliers omitted). Each panel is annotated with the vertex-level Spearman correlation between the per-vertex feature and the ordinal cortical type, its spin-test *p* (1000 rotations) and FDR *q* (*∗*: q < 0.05). Myelin increases with laminar complexity (*ρ* = +0.59) while thickness (*ρ* = *−*0.50) and the functional gradient (*ρ* = *−*0.58) decrease, recovering the canonical thin/heavily-myelinated primary-sensory versus thick/lightly-myelinated limbic organization. Surfaces are shown on the 32k fs_LR mesh, where these features are natively defined; the box plots display all seven types, while the correlation is computed as in the Methods.

Two pre-specified negative controls behave as they should: sulcal depth and cortical curvature, which carry no cytoarchitectural prediction, return *ρ* = *−*0.04 and *−*0.05 and do not approach the spin null (*p* = 0.66 and 0.53), so the pipeline is not manufacturing correlations from surface geometry alone. The three robust features are themselves moderately inter-correlated (|r| 0.37–0.57), indexing one sensorimotor-to-association axis rather than three independent tests, and the trends are not artefacts of the type exclusions: the myelin correlation is +0.589 over all seven types, +0.604 with allocortex excluded and +0.585 with agranular excluded as well, each surviving its spin null at *p* = 0.001, so the exclusions move it by less than 0.02 in either direction (robustness listing, Table S9).

One further exclusion is owed for the same reason. Primary motor cortex is typed 6 despite its thin layer IV, because the protocol grades overall laminar complexity rather than layer-IV thickness alone, and motor cortex is otherwise highly complex ^3^. It is also large, heavily myelinated and unusually thick, so it could inflate the myelin correlation and flatten the thickness one at once. Excluding the precentral territory (4,035 vertices, 6.9% of labelled cortex) moves all four primary correlations by less than 0.04, in the directions the concern predicts: myelin +0.59 to +0.57, thickness *−*0.50 to *−*0.54, the functional gradient *−*0.58 to *−*0.54 and gene expression PC1 +0.67 to +0.66, each still surviving its spin null.

### 3.6 A frequency/timescale gradient

Resting-state MEG dynamics track the type axis. The group-average neuromaps versions of these measures, the intrinsic-timescale and band-power maps in the nine-feature panel of §3.5 (Table S3), trend as predicted but do not survive the spin null, so we recomputed the dynamics per subject: we reconstructed source-level power spectra from the HCP young-adult MEG (N = 89; §2.5) and, per subject, separated each spectrum into its aperiodic (1/f) component and oscillatory peaks, then tested each derived measure against ordinal type under the spin+FDR null.

The intrinsic timescale shortens toward koniocortex and survives the null: the broadband autocorrelation timescale decreases with type (*ρ* = *−*0.46, spin *p* = 0.004, *q* = 0.014), and the exponential-decay, 1/e and knee-derived timescale definitions agree in direction (all spin *p* < 0.05). The per-type medians decrease monotonically from the agranular belt to koniocortex (46.8 to 21.9 ms) but the allocortical median is faster than the agranular one (41.5 ms), so the slowest cortex on this measure is agranular rather than allocortex; we read this as the allocortex being poorly resolved by MEG source estimates rather than as a reversal of the gradient, consistent with its exclusion from the trend fits (§2.6, Table S3). The frequency measures follow: the power-weighted spectral centroid increases (*ρ* = +0.41, *q* = 0.014), the slow-to-fast band-power ratio (*δ* + *θ*)*/*(*β* + *γ*) decreases (*ρ* = *−*0.36, *q* = 0.021), and aperiodic-corrected *β* power increases (*ρ* = +0.22, *q* = 0.021). Dynamics are faster and shorter-timescale toward the more complex end, resolving the resting-state frequency gradient of Mahjoory et al. ^18^ on the cytoarchitectural axis with every subject contributing in the predicted direction for the primary timescale (89 of 89; Annex F) (Supplementary Fig. S1; Annex F; Table S3).

No association with type was detected for the aperiodic exponent (*ρ* = *−*0.12, spin *p* = 0.50), an indirect spectral correlate proposed to be sensitive to excitation–inhibition physiology ^43^, under its pre-specified fixed-mode specification. The measurement itself is sound: the source spectra fit well (99.9% with *R*^2^ *≥* 0.9), the exponent is reproducible across the three resting-state recordings (across-run *r* = 0.87), and the reconstruction is validated (source *α* matches the reference *α* map at *ρ* = 0.98 and the broadband timescale matches the reference intrinsic-timescale map at *ρ* = 0.85). Good fit and reproducibility do not establish that the effect is negligible, which would need an equivalence analysis we have not run, and the sign reverses under the knee-mode fit (Annex F). Peak frequency and the spectral offset are also null: the discrete spectral peak is well defined only where a clear oscillation exists, so the dominant-frequency gradient is better read from the power-weighted spectral centroid (above), which survives the spatially constrained spin null that the distance regression of Mahjoory et al. ^18^ did not apply. Cortical type is therefore reflected in the speed of cortical dynamics, whereas no association was detected for the aperiodic exponent; we take up this dissociation in §4.5.

### 3.7 Convergence with predicted molecular and cellular gradients

Two molecular references, not used as inputs to the painting, were tested against the type axis (Fig. 5; Table S3; Supplementary Fig. S6; Annex H). The dominant transcriptomic gradient (AHBA gene-expression PC1) increases monotonically across cyto7 types (*ρ ≈* +0.67, spin *p* = 0.001, *q* = 0.002); because it correlates *r ≈* 0.60 with the functional gradient, it is best read as a fourth (transcriptomic) window on the one sensorimotor–association axis rather than an independent test. The usual AHBA caveats apply and are sharper here than the donor count suggests: five of the six donors contribute, the samples are predominantly left-hemisphere, and the parcel values are produced with bidirectional mirroring and interpolation of unsampled regions, so the 68 parcels are not 68 independent tissue observations (Annex H).

**Figure 5.**
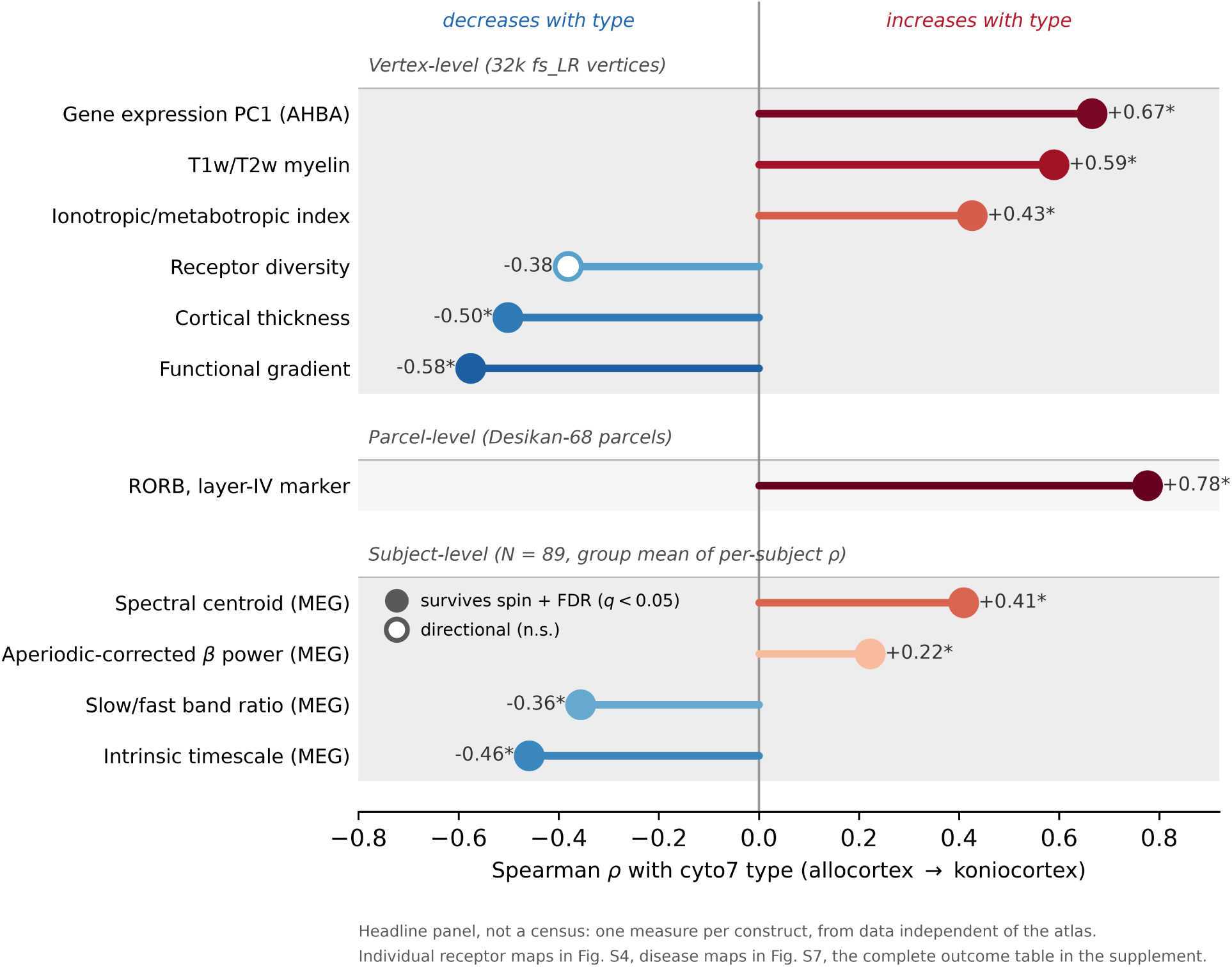
Independent cortical measures order along the cyto7 gradient of laminar complexity. For each measure, the vertex-level Spearman correlation with ordinal cyto7 type (allocortex *→* koniocortex) is shown as a signed value: measures with positive *ρ* increase toward koniocortex (red) and those with negative *ρ* decrease (blue), with bar length the strength of the association. Filled markers denote measures that survive the spatial-autocorrelation (spin) null with FDR control (*q <* 0.05) within their test family; open markers denote directional trends that do not. Measures are grouped into three bands by statistical unit and are comparable only within a band: vertex-level correlations on the 32k fs_LR mesh, a parcel-level correlation on the Desikan-68 parcellation (RORB, for which AHBA expression is available only at parcel resolution), and subject-level statistics that are the group mean over 89 subjects of each subject’s per-vertex correlation (§3.6, Annex F). Ten of the eleven measures shown survive: gene-expression PC1, T1w/T2w myelin, the ionotropic-to-metabotropic receptor index, the layer-IV marker RORB, the spectral centroid and aperiodic-corrected *β* power increase with type, while cortical thickness, the principal functional gradient, the intrinsic timescale and the slow/fast band-power ratio decrease; receptor diversity decreases in the predicted direction but does not survive the null. This panel is a declared headline selection rather than a census: it shows one measure per construct, drawn from data not used to paint the atlas. Individual receptor maps are in Fig. S4, individual disease maps in Fig. S7, the 147 entries of the declared families in Table S8, and the robustness and sensitivity analyses in Table S9. The count below is the survival rate within this selected panel, not an overall success rate for the study. Recovered from data independent of the atlas, this reproduces the direction of several molecular and cellular gradients associated with cortical type ( ^4^, Fig. 9). The signed correlation is the appropriate summary, because the per-type median trajectory of some measures is non-monotone (the functional gradient, for instance, peaks in transmodal association cortex).

One further test bears directly on the defining criterion. The layer-IV excitatory-neuron marker *RORB*, predicted in advance to increase with type, does so steeply along the axis (*ρ* = +0.78, parcel spin *p* = 0.003, parcel-level; robust to allocortex exclusion, +0.76). RORB is a canonical layer-IV marker, so this is molecular convergence with the layer-IV granularity gradient that is the principal criterion behind cyto7, bearing on a defining feature where most other measures index only the generic hierarchy. The axis is not defined by layer IV alone, since layer III soma size is a second criterion (Box 1), and no transcriptomic test here speaks to that one; what RORB establishes is convergence on the granularity criterion, not on the whole of the typing. Because the test is at parcel resolution, it supports that criterion rather than the placement of any individual boundary. The wider laminar-marker program agrees: the upper-layer marker CUX2 and the deep markers FOXP2 and TLE4 increase, while the layer-V marker FEZF2 decreases (*ρ* = *−*0.59), the laminar specificity expected if the axis reads genuine layer identity (Supplementary Fig. S9). One qualification applies to the transcriptomic tests generally. The expectation that expression increases with type is not uniform across genes but depends on what the gene does: markers of glutamatergic transmission should track the axis, whereas markers of GABAergic transmission need not, since inhibitory populations do not scale with lamination in the same way. PC1 and the laminar markers do not separate genes by transmitter class, so neither result speaks to that distinction, and a class-resolved test is left to future work (§4.10). Cortical type has been proposed as the organising variable for exactly this kind of high-throughput expression analysis ^46^.

The neurotransmitter-receptor architecture also reorganises along the type axis, but this is apparent only when the receptors are examined individually. The first principal component of the 19 PET receptor and transporter densities shows only a weak decreasing trend (*ρ ≈ −*0.33) that does not survive the spin null. Individual receptor systems do track the axis, in both directions, and we have not established why PC1 does not reflect that: a principal component carries signed loadings and can represent opposing patterns, so opposing directions do not by themselves force a null component. Examined map by map (with FDR control across the 19-receptor family), four maps survive, and the directions are specific: the *µ*-opioid receptor (*ρ* = *−*0.55), the serotonin 5-HT_1A_ receptor (*ρ* = *−*0.54) and the cannabinoid CB_1_ receptor (*ρ* = *−*0.41) increase toward the limbic belt, whereas the noradrenaline transporter (NET) increases toward koniocortex (*ρ* = +0.44; all *q* = 0.043). Which receptors run in which direction, for all 19 maps, is given in Supplementary Fig. S4 and Annex H. Grouping the maps by pharmacological class recovers the axis that the PC1 obscures: a metabotropic composite decreases with type (*ρ* = *−*0.41) against an approximately flat ionotropic composite, so that an ionotropic-to-metabotropic index increases from the limbic belt to koniocortex (*ρ* = +0.43, *q* = 0.016; Supplementary Fig. S5). This is the receptor-level counterpart of the molecular and cellular gradients associated with cortical type ^2,4^, recovered here from independent PET data; a clean excitatory-versus-inhibitory (glutamate-versus-GABA) contrast, by contrast, is under-powered in this predominantly neuromodulatory group of receptors and does not survive the null (Annex H). A direct histological comparison (BigBrain) is inconclusive in a single specimen and is discussed as a limitation (§4.9, Annex H).

Taken together, these readouts let us ask directly how far cyto7 recovers the molecular and cellular gradients associated with cortical type ( ^4^, their Fig. 9). Of the axes we could test from independent data, the ionotropic-to-metabotropic receptor balance increases with type and survives the spin null (§3.7), while receptor diversity, estimated as the Shannon entropy of the 19 PET maps, decreases in the predicted direction (*ρ* = *−*0.38) though it does not survive FDR. Evolutionary cortical expansion decreases with type in the predicted direction (*ρ* = *−*0.27) but does not survive FDR, and developmental expansion is flat (*ρ* = *−*0.11, *p* = 0.42; Table S8). We do not build on either. Expansion is not a property of one cortex but of a correspondence between two, and that correspondence has to be fixed before expansion can be measured: the available estimates fix it from functional-connectivity similarity ^38^ or surface registration ^39^, whereas cortical type analysis fixes it from laminar structure itself ^9^. A test of our map against those estimates therefore weighs two homology criteria against each other rather than testing type against expansion. The informative version is a direct human-to-macaque comparison in areas typed by the same criteria (§4.10).

Disease vulnerability, which Zaldívar-Díez and García-Cabezas ^4^ place at the low-complexity end of the axis, shows the same partial convergence at parcel resolution: of eight ENIGMA case–control cortical-thickness maps, major depression, ADHD, obsessive–compulsive disorder and 22q11.2 deletion track lower cyto7 type (depression and ADHD surviving control for the functional gradient), whereas epilepsy runs opposite and three disorders are null, so we report it as convergent context rather than a predictive claim (Annex H; Supplementary Fig. S7).

Across families, the alignment is partial but consistent, and Fig. 5 summarises it as each measure’s signed correlation with ordinal type. The surviving measures span microstructure (myelin increasing, thickness decreasing), the functional-connectivity gradient (decreasing), molecular architecture (gene-expression PC1 and the ionotropic-to-metabotropic index increasing) and resting-state dynamics (the intrinsic timescale decreasing, with the concurring frequency measures of §3.6); receptor diversity trends as predicted but does not pass the null, and no association was detected for the aperiodic exponent under its primary specification (§3.6; it is not shown in Fig. 5). Ten of the eleven measures in Fig. 5 survive their family null. The type signature is therefore recovered across microstructural, molecular, macroscale-gradient and dynamical measures, from data not used to paint the atlas. Full per-type detail is in the fingerprint heatmap (Fig. S2), the timescale and band-power surfaces (Fig. S1), and Table S3.

## 4. Discussion

### 4.1 Main contribution

To our knowledge, cyto7 is the first vertex-level, allocortex-aware, openly released map of the discrete cortical types of García-Cabezas et al. ^3^ that ships with a per-vertex support score. Three claims of different strength should be kept apart. That the cortex is ordered along a gradient of laminar complexity, and that cyto7 recovers it, is strongly supported here by independent imaging, molecular and dynamical data. That seven discrete types are the right quantisation of that axis is inherited from the protocol and is moderately supported. That any individual vertex-level boundary sits exactly where cyto7 places it is a hypothesis this release makes testable, not a result we establish; it awaits independent raters and multi-specimen histology. What it offers is not a new type scheme, as it follows the seven types of García-Cabezas et al. ^3^, but a change of resolution: we place type boundaries vertex by vertex, so a transition that runs *through* a classical area is placed at vertex resolution from the expert anatomical synthesis and the topological constraints of the gradient of laminar complexity rather than collapsed to a single areal label, and we state, per vertex, how strongly that placement is supported.

This makes cyto7 complementary to, not competing with, the continuous microstructural axes now available (BigBrain-derived microstructural profile covariance, and the data-driven cytoarchitectural axis of ^47^) and the area-level atlases that preceded it ^21,22^. A continuous axis asks how graded the microstructure is; cyto7 asks which discrete laminar type a vertex belongs to, under an explicit topological constraint. The two answer different questions, and a discrete map with a per-vertex support score is what many downstream uses (stratifying an analysis, setting per-node model priors, parameterizing laminar structure) actually need.

The painting, the reconciliation with the topological constraints of the gradient of laminar complexity, the type scheme and the benchmark’s typing rule are expert-internal: the protocol’s authors are co-authors. The multimodal imaging validation is the one fully external, independent leg.

### 4.2 What the fsaverage surface represents

Furthermore, a recurring source of confusion is clarified. On the fsaverage surface, allocortex is represented by the olfactory paleocortex (piriform) and the modified hippocampal allocortex (entorhinal cortex, pre-/parasubiculum); only the hippocampal allocortex sensu stricto (fascia dentata, cornu ammonis, subiculum) is off the surface, in the volume, because FreeSurfer represents the hippocampal formation volumetrically. This carries a topological consequence for the dual origin of the neocortex theory. Its concentric growth rings emanate from two allocortical origins, olfactory (paleocortex) and hippocampal (archicortex), yet on the cut surface both origins lie on a single rim around one medial-wall hole, so the allocortex–mesocortex belt forms a single continuous annulus rather than two closed loops: the two origins appear as adjacent sectors of that belt, with the strict sequential gradient (allocortex through mesocortex to isocortex; R1) running from each. The genuine double loop closes through the hippocampal formation, which lies off the surface; that closure is a property of the three-dimensional mantle. A faithful archicortical surface representation would require dedicated hippocampal unfolding ^48^. That the two origins are separable is supported independently of cytoarchitecture: Sebenius et al. ^49^ recover two principal gradients of a genetically defined cortical similarity network whose weightings track geodesic distance from the piriform and the parahippocampal cortex respectively.

### 4.3 Structure–function convergence

The central empirical result is that three correlated in-vivo maps, intracortical myelin (T1w/T2w), cortical thickness, and the principal functional-connectivity gradient, correlate with ordinal type beyond a spatially constrained autocorrelation null (spin + FDR). The first two also vary near-monotonically across the seven types, reversing at only one of six steps; the functional gradient does not, rising from allocortex to eulaminate I and falling steeply thereafter, so its correlation with type reflects that descent rather than a monotone climb. Each map is a proxy of the same sensorimotor-to-association axis. Two features matter. First, the axis was defined from laminar cytoarchitecture, without using any of the maps it is validated against, so the convergence is with independent data rather than a re-description: an anatomy-defined ordering predicts the organisation of microstructural and functional MRI, the concrete corroboration that cortical type analysis anticipates ^3^, and it aligns cyto7 with the sensorimotor-to-association gradient literature ^12–15^. Convergent evidence has since come from a different direction: Oh et al. ^23^ report that a hierarchy derived from effective connectivity increases as granularity decreases across cortical types. That corroborates the gradient of laminar complexity as an organiser of macroscale function, using a coarser area-level type map than the one released here, and it is not a validation of cyto7. One dependency is worth naming precisely. No validation value entered the painting, but the limbic belt was adjudicated in part against the Glasser HCP-MMP parcellation (§2.1), which was itself built from multimodal HCP data including myelin maps, and the imaging validations use HCP group averages on the same registration; the overlap is at the level of shared anatomical references, cohort and registration rather than shared numbers, and it is confined to the belt. Excluding the vertices where the Glasser term was decisive (4,558 vertices, 1.5% of labelled cortex) moves no primary correlation by more than 0.017. Excluding the entire belt, a far more conservative test that also removes three of the seven types and most of the axis range, leaves myelin (*ρ* = +0.59), thickness (*−*0.44), the functional gradient (*−*0.62) and gene PC1 (+0.58) all surviving their nulls; the MEG intrinsic timescale keeps its sign and 88 of 89 subjects but is no longer significant over the isocortical range alone (*−*0.35, spin *p* = 0.066). Second, the progression is graded, not a set of disconnected category means: myelin increases and thickness decreases toward the granular koniocortical end while the functional gradient runs the other way, recovering the canonical view in which primary sensory cortex is thin and heavily myelinated and limbic/association cortex is thick and lightly myelinated. Because it is anatomy-defined, cyto7 can stratify or parameterise imaging analyses without the circularity of deriving the axis from the data it then explains. It constitutes a discrete, biologically interpretable scaffold for the same hierarchy that data-driven gradients capture.

What that convergence does not show is that the anatomical labour was necessary to obtain it. Tested directly against seven-level quantisations of T1w/T2w myelin, of the principal functional gradient and of the BigBrain microstructural gradient, cyto7 attains the higher correlation on none of the twelve vertex-level features (Annex J). Part of that is arithmetic: a septile reproduces its own source map at |*ρ*| = 0.99. How a septile performs on the other features is not, and is an empirical result. The difference appears where the measures speak to the defining criterion rather than to the generic hierarchy. On the laminar marker genes cyto7 shows the largest absolute correlation of the four partitions for RORB, CUX2, FOXP2 and FEZF2, and a smaller one only for TLE4, with the widest margin on FEZF2 (0.60 against 0.17 for the myelin septile). Direction does not explain that margin: because the comparison is of absolute correlations, a myelin septile can represent a feature running against myelin, simply with the opposite sign. What the pattern shows is that cyto7 ties or loses on macroscale proxies and has larger observed associations on the layer markers, which is what would follow if the anatomy carries laminar rather than hierarchical information. We have not tested whether those between-map differences are statistically reliable, which would need a paired null preserving the dependence among the competing maps (§4.10). A reader who wants only an ordering of cortex along the unimodal-transmodal axis does not need this atlas; a reader who wants discrete, interpretable laminar categories may.

### 4.4 T1w/T2w as a myelin proxy, not a measurement

The T1w/T2w ratio is here interpreted as a relative index of the myeloarchitectural gradient, not as quantitative myelin. It was introduced as a practical contrast that suppresses receive-field bias and tracks heavily myelinated cortex ^10^, but it is not myelin-specific: it is additionally sensitive to iron, water/free-water content, axon and dendrite density, and residual transmit-field (B1+) inhomogeneity, and it lacks biophysical calibration ^50–52^; group-averaging and surface projection blur it further. The myelin claim is thus directional and relative; refining it further would require quantitative measures (MT saturation, quantitative T1/R1, g-ratio mapping; ^53^) or histological calibration (BigBrain; ^40^). Reassuringly, thickness and the functional gradient, which do not share these confounds, track the same axis, so the convergence does not rest on the myelin proxy alone. The proxy is least reliable in the allocortex, where the ratio is elevated by superficial and juxtacortical myelin (the alveus, the lateral olfactory tract, the presubicular myelin band) and by partial volume with adjacent white matter in thin cortex, not by intracortical laminar myelin; this is why allocortex is excluded from the myelin trend fits (§2.6).

### 4.5 A frequency/timescale gradient

We had predicted that dynamics would track type: faster, shorter-timescale activity toward the granular, koniocortical end of the axis. Source-reconstructing the HCP young-adult MEG per subject, instead of reanalysing the group-average maps, resolves it on the cytoarchitectural axis in this cohort (§3.6). The prediction holds: the intrinsic timescale shortens and the spectral centroid, the slow/fast band-power ratio and aperiodic-corrected beta power move as predicted and survive the spin+FDR null, reproducing the resting-state frequency gradient of Mahjoory et al. ^18^ on the cytoarchitectural axis.

The aperiodic 1/f exponent is the exception: no association with type was detected under the pre-specified fixed-mode fit, on well-fitted and reproducible spectra (§3.6), though the sign reverses under the knee-mode alternative, so this is specification-sensitive rather than a settled negative. The dissociation does not rest on one measure surviving where another fails, which would be weak given that both summarise the same spectrum. We also tested it conditionally: the timescale still tracks type after partialling out the aperiodic exponent and all five coded band powers (*ρ* = *−*0.28, nominal spin *p* = 0.009, 88 of 89 subjects), an attenuation of 38% from *−*0.46, whereas the exponent acquires no association with type when the timescale is partialled out of it (*−*0.05, *p* = 0.82). Two limits on how far that goes. The *δ* and *γ* covariates are poorly estimated under the 2–40 Hz fit (Annex F), and a control restricted to the three reliable bands has not been run, so we do not know how the result behaves under that specification. And the *p*-values are nominal rather than FDR-controlled within a declared family. We therefore report this as an exploratory conditional association: the timescale retains an association with type after adjustment for these particular covariates, which is a narrower statement than showing that shared spectral structure has been adequately controlled. Because the exponent is at best an indirect correlate of excitation–inhibition physiology ^43^, the absence of a detected association concerns that spectral feature and not excitation–inhibition organisation as such. That resting timescales lengthen along the sensorimotor-to-association axis and track cortical microstructure has been shown from intracranial recordings ^19^; what the present result adds is that the ordering follows the discrete cortical types of García-Cabezas et al. ^3^, defined from lamination rather than from a continuous macroscale gradient, and that it holds per subject. Whether type structures excitation–inhibition through channels the resting exponent does not capture, such as laminar receptor densities or interneuron composition, is a question for laminar modelling; a second cohort would test whether the absence of a detected association reproduces.

### 4.6 Multimodal fingerprints as type phenotypes

Combining all the information of the different modalities per type yields a multimodal *fingerprint*: a compact vector of structural and functional properties (myelin, thickness, band-limited MEG power, intrinsic timescale, functional-gradient position) attached to each type. Making these explicit is useful in three ways. Descriptively, they convert an ordinal label into an interpretable phenotype and show which properties separate types cleanly (myelin, thickness, the functional gradient) versus weakly (band-limited power). As a tool, type fingerprints are candidate priors for the regional heterogeneity that whole-brain models need (more on this, below), and a template against which an individual’s or a novel area’s profile can be matched and typed. And *normatively*, a region whose fingerprint departs from its type’s expectation is a data-driven flag for atypical microstructure, the natural bridge to the soft, probabilistic per-vertex typing we propose below. We keep these data-derived signatures separate from the anatomical support score to avoid circularity, but they are the empirical counterpart to the type definitions.

### 4.7 cyto7 as a substrate for large-scale models

A possible application of this map is to serve as a substrate for large-scale dynamical models of cortex, which improve sharply when regional parameters are made *heterogeneous* along a biological gradient instead of held uniform: tuning local recurrent excitation by the T1w/T2w myelin map improves fits to resting-state dynamics and functional connectivity ^54,55^, and a hierarchy of intrinsic timescales is a core ingredient of such models ^16^. But these gradients are read off imaging, often the very functional data the model is then asked to reproduce (the functional-connectivity gradient, intrinsic timescales), or structural maps closely correlated with it. Tuning a model by one and scoring it on the other therefore explains the data partly with a summary of itself. What is missing is a heterogeneity axis whose relationship to function is discovered rather than built in.

cyto7 offers a discrete, anatomy-defined alternative. The seven types can set per-node laminar structure and the parameters that follow from it (excitation–inhibition balance, recurrent gain, intrinsic time constants) from cytoarchitecture rather than from the imaging the model is meant to predict. This is most natural for laminar neural-mass formulations ^56^, where the type *is* the laminar architecture the model represents: the ordering supplies a directional prior on regional time constants (faster toward more complex cortex), and the same scaffold carries over to modelling brain-stimulation responses and type-dependent vulnerability. We stop short of reading calibrated per-type electrophysiological values off the MEG. The dynamics gradient is significant (§3.6), but even a per-subject correlation with type is not a set of calibrated per-type time constants, and for the aperiodic exponent, the measure closest to an excitation–inhibition parameter, no association with type was detected. cyto7’s role here is the scaffold, not the numbers that fill it.

For region-based whole-brain models which place one node per parcel, each Desikan or von Economo parcel can take its majority cyto7 type as a per-node prior (Table S5). This is convenient but lossy: the average classical parcel spans about two types (§3.3), so one type per parcel misassigns the type of about 40% of labelled cortex (34% for von Economo), which is exactly what the vertex map keeps and can hand to vertex- or laminar-resolved models. There is a straightforward middle course, which we adopt: intersecting a classical parcellation with cyto7, so that a heterogeneous parcel becomes several type-homogeneous nodes. This is the product that the type-by-parcel transfer of Oh et al. ^23^ was improvising, and it is released in the fs_LR space their parcellation already occupies. Crossing the released map with the Desikan–Killiany atlas yields 101 sub-parcels above 200 vertices across both hemispheres from 34 parcels (a mean of 3.0 per parcel; 115 from the 43 von Economo parcels), and discarding every cell at or below that size costs 0.6% of labelled vertices (1.2% for the von Economo crossing), so the result is a node-based parcellation of about a hundred regions rather than a fragmentation into slivers. Supplementary Figure S14 makes the cost of each step explicit: relative to the vertex map, the crossed parcellation misassigns the cyto7 type of 0.6% of labelled vertices (1.2% for von Economo), the residue absorbed by the minimum node size, whereas one majority type per parcel misassigns 40.1% (33.5% for von Economo). It drops into existing region-based pipelines while no longer averaging across a type boundary the atlas exists to resolve. We release it as FreeSurfer annotation files on fsaverage, so it transfers to any subject with a completed cortical reconstruction through the standard spherical registration, alongside 32k fs_LR label files, a colour table and a per-node table giving the base parcel, the cyto7 type and the node geometry. We report three build choices: a 200-vertex minimum node size evaluated across both hemispheres, so that a node may fall below it in one hemisphere and per-hemisphere counts are given in the node table; absorption of the residue into the largest neighbouring node within the same anatomical parcel, so that no anatomical boundary is crossed; and retention of one node per parcel-by-type cell even where it spans more than one patch, with per-node component counts reported so that a modeller who needs contiguous nodes can split them. Users who prefer different choices can rebuild from the released per-parcel type compositions (Table S5; Annex I). Each node inherits the local boundary uncertainty discussed in §4.1.

### 4.8 Individual variability and scope

cyto7 is a group-average, template-space map, and cortical organisation is known to vary across individuals, with functional and areal variability greatest in association and limbic cortex and least in unimodal cortex ^57–59^. We cannot yet quantify this variability for cyto7 itself: we have a single group delineation, and the per-vertex support score reports how well the *template* boundary is backed by the anatomical evidence, not how that boundary moves across people, and it is calibrated to neither. Doing the latter would require per-subject cortical-type delineations that we do not have: either many individual paintings/raters, or a validated way to infer type from individual microstructure. Two types of studies point the way for a future, individualised cyto7: observer-independent probabilistic cytoarchitectonic mapping, which quantifies inter-subject variability of areal borders from serially-sectioned post-mortem brains (^60^; Julich-Brain, ^29^), and individual-specific parcellation from in-vivo MRI ^58,59^. Such a map would carry, per vertex, both a type distribution and a between-subject variability estimate, which is complementary to, but distinct from, the template-delineation support indexed here. As a first empirical step, projecting the group map onto 200 individual HCP subjects shows the type-to-myelin ordering holds in every one: each subject’s T1w/T2w correlates with ordinal type at a median *ρ* of 0.53, with a strictly monotonic agranular-to-koniocortex increase in all 200, and the group-average value of 0.59 is an upper bound on the individual fit, not a typical one (Supplementary Fig. S10).

### 4.9 Limitations

Several limitations bound what cyto7 can claim. We list them by our perceived order of importance.

#### It is hand-painted by a single rater

Expert revision by the protocol’s authors and the support score mitigate this, but there is no independent second rater, so we cannot report inter-rater reliability, and the support score reflects how well the *template* boundary is backed by anatomical evidence rather than disagreement between raters. The one internal check we can offer is cross-hemisphere consistency: the two hemispheres were painted independently under the same criteria and agree at *κ_w_* = 0.96 (Annex A), which bounds gross within-rater inconsistency but is not a substitute for independent raters. A second-rater study is the most direct way to harden the map.

#### Direct histological validation is still outstanding

The typing ultimately rests on classical cytoarchitectonics and a finite set of directly examined material (the von Economo–Koskinas atlas and the Nissl-stained post-mortem sections analysed under the García-Cabezas protocol), plus expert judgement, rather than a quantitative survey of many hemispheres. Our one direct check, against BigBrain ^40^, did not support the typing, and its two forms failed differently: neither the microstructural-profile-covariance gradient nor a pre-specified profile index (skewness) resolved the seven types under the spatial null on the fs_LR projection, and on the disagreement set of §3.3 the Hist-G1 arbiter returns a measured null slightly below chance (0.4993 against 0.519). Taken together, these analyses did not establish histological validation of the seven-type scale, nor improved performance over the area-level reference. Descriptive profile differences are present, with the mean intensity profiles differing in shape by type and koniocortex showing the expected mid-cortical peak, and profile SD tracks type in an exploratory analysis (*ρ ≈* 0.44, *q* = 0.015). What we cannot say is why the pre-specified comparisons failed: the contributions of specimen variability, the projection to a group surface, statistical sensitivity and the choice of scalar summary remain unresolved. One methodological consideration is worth naming among those candidates: BigBrain’s Merker staining is a direct cytoarchitectural measurement, more direct than the MRI proxies used elsewhere here, but the scalar index we extract from it (profile skewness) is a one-dimensional summary of the laminar depth profile, whereas a cortical type integrates the full laminar pattern (layer-IV emergence and granularity, layer III soma size, sublamination, border sharpness); a single profile moment need not recover the type axis even when the histology carries it. A definitive test needs a multi-specimen, 3-D-reconstructed reference (of order ten sex-balanced brains), which would also turn the discrete labels into a properly probabilistic type map.

#### The surface allocortex is delineated on developmental grounds

Entorhinal cortex and the pre-/parasubiculum share the hippocampus’s medial-pallial origin ^28^, so we type them from embryology rather than from the surface atlases, following the protocol of García-Cabezas et al. ^3^ ; the support score marks this tissue transparently as low-support (§3.4). The hippocampal allocortex *sensu stricto* is off the fsaverage surface and deferred to volumetric/unfolded representations.

#### The architectonic-type principle is not tested here

The principle is a claim about the laminar pattern of cortico-cortical connections: projections between areas of different type originate and terminate in layers that the direction and magnitude of the type difference predicts^1,2^. A streamline carries no laminar information, so diffusion tractography can ask whether connection density is patterned by cortical type, which is what the Annex G analysis does, but cannot address the principle itself. This limit is independent of the statistical objections to that analysis, and it is why the result is offered as consistent with the principle rather than as evidence for it. The relevant test needs laminar-resolved connectivity, which in the human brain is not currently available in vivo.

#### Other remaining caveats

The tractography uses a single population-averaged template with a deliberately high 80 mm short/long split; the type-distance falloff attenuates by about a third once connection counts are expressed per available endpoint pair, and then fails both a spin null and, more tellingly, a bundle-level model, so it is reported as exploratory only (Annex G); the spin null is spatially constrained rather than conservative in general, and loses power for smooth normative maps (band-limited MEG most of all); and the ordinal type scale is treated as equally spaced, which is a simplification. Some regions, finally, are genuinely unsettled, and retrosplenial cortex is the clearest case. We type it as mesocortex, following the histological protocol we implement, in which the retrosplenial areas of von Economo and Koskinas (LD, LE1, LE2) are agranular and dysgranular ^3^; the crossed von Economo parcellation reproduces this, and the eulaminate I sector that falls inside the Desikan–Killiany *isthmus cingulate* parcel belongs to the adjacent dorsal posterior cingulate area LC1, which that protocol likewise types as eulaminate I. Genoarchitectonic evidence in the rodent has since moved the boundary: Puelles et al. ^61^ show that a substantial part of the adult mouse retrosplenial cortex is modified allocortex rather than mesocortex, and Zaldívar-Díez and García-Cabezas ^4^, following that work and Puelles et al. ^28^, place some retrosplenial areas in the medial-pallial ring together with the entorhinal cortex, in the strip between the indusium griseum and tenia tecta and the agranular cingulate mesocortex. Since our lowest type already carries the entorhinal cortex on exactly that basis, a modified-allocortical retrosplenial sector would be consistent with the scheme. We do not draw one here: the equivalent analysis has not been carried out in the human cortex, and the reclassification does not yet specify which retrosplenial areas would be affected or where the boundary falls on the cortical surface. Future studies of postmortem human brains may reclassify part of this cortex, and this is precisely the kind of borderline area a versioned atlas exists to revise.

### 4.10 Future work

Several of the previously discussed limitations pave the way to future extensions of this work. (1) An archicortex companion built with hippocampal unfolding ^48^ would extend the allocortex from the current surface periarchicortex to the hippocampal formation sensu stricto and its subfields, in a representation matched to the sheet. (2) Confirming the resting-dynamics gradient in a second MEG/EEG cohort and probing excitation–inhibition balance directly (laminar receptor densities, interneuron fractions), to test whether cortical type organises excitation–inhibition beyond the resting-MEG aperiodic exponent, which does not track it (§3.6). (3) Soft, probabilistic per-vertex type vectors (a distribution over types at each vertex, informed by the multimodal fingerprints) would replace hard labels with graded typing and explicit uncertainty, complementing the support framework. (4) Individualisation: projecting or learning the map on single subjects, with a between-subject variability layer anchored to probabilistic-cytoarchitectonic and individual-parcellation work ^29,59^. (5) Using cyto7 as the heterogeneity substrate for whole-brain and laminar neural-mass models, and for stimulation and disease-vulnerability modelling. (6) Replacing the T1w/T2w proxy with quantitative myelin measures and broadening the benchmark to volumetric and histological references (BigBrain/Julich-Brain). (7) A direct human-to-macaque comparison in matched cortical types, which is the form an evolutionary test of this axis should take: the available whole-cortex expansion estimates rest on cross-species alignments whose phylogenetic assumptions differ from the cortical-type framework, so comparing types across species that have been typed by the same criteria is the informative version of that question ^9^. (8) Splitting gene expression by transmitter class, to test the class-dependent prediction noted in §3.7 that glutamatergic markers should track the axis where GABAergic markers need not. Each builds on, rather than revises, the released map.

## Declaration of the use of AI

Generative AI tools (a large language model) were used to assist with drafting, editing and condensing the manuscript text. All study design, analyses, results and interpretations are the authors’ own; the authors reviewed all AI-assisted text and take full responsibility for the content of this article.

## Data and Code Availability

The atlas annotations, together with the initial as-painted map and the change log as provenance, the per-vertex support maps, and the quality-control / comparison / analysis pipelines are released under an open license (data under CC BY 4.0, code under the MIT license). All files, including the large binaries, are distributed directly in the repository (https://github.com/bcom-foundation/bcom-collab-cyto7), with a SHA-256 manifest so that any download can be verified against the released state, and with a versioned archive on Zenodo (https://doi.org/10.5281/zenodo.23063166). The crossed parcellation is released at atlas/fsaverage/crossed/ as FreeSurfer annotation files on fsaverage (?h.aparc_cyto7.annot, ?h.voneconomo_cyto7.annot), so it transfers to any subject with a completed cortical reconstruction through the standard spherical registration with a single mri_surf2surf –sval-annot call; 32k fs_LR label files (?h.aparc_cyto7.32k_fs_LR.label.gii and the von Economo equivalents) at atlas/fs_LR_32k/crossed/ for surface-based pipelines in that space; FreeSurfer colour tables (aparc_cyto7.ctab, voneconomo_cyto7.ctab); and per-node tables (aparc_cyto7_nodes.tsv, voneconomo_cyto7_nodes.tsv) giving for each node its identifier, name, hemisphere, base parcel, cyto7 type, vertex count, surface area, number of connected components and the fraction of vertices in the largest, together with a secondary contiguity-split variant (?h.*_cc.annot), a usage guide (README_crossed.md) and a provenance record with input checksums (PROVENANCE_crossed.txt). Node names split on the final underscore into the base parcel, spelled as in the source atlas, and the cyto7 type. The transfer was verified by mapping the annotation to a subject with a completed recon-all and reading it back, with every node recovered. All third-party datasets used for validation (HCP, ENIGMA, Allen Human Brain Atlas, the Hansen PET collection, BigBrain, and the neuromaps distributions) are publicly available from their original providers under their respective data-use terms.

## Author contributions

R.S.: Conceptualization, Methodology, Software, Formal analysis, Investigation, Data curation, Visualization, Writing – original draft, Writing – review & editing. B.M.: Methodology, Software, Formal analysis, Writing – review & editing. F.C.: Methodology, Formal analysis, Writing – review & editing. M.Á.G.-C.: Conceptualization, Validation (cortical-type criteria, reconciliation and expert revision of the map), Writing – review & editing. G.R.: Conceptualization, Supervision, Funding acquisition, Writing – review & editing.

