## Supplementary Material: Figures S1-S14, Tables S1-S9 for "cyto7: an open, vertex-level cortical type atlas of the human cortex"

This document contains Annexes A–J: extended methods, benchmark metrics, external validation against molecular and histological references (genes, receptors, BigBrain), the cortical-type composition of classical parcellations, the comparison against quantised continuous maps, and reproducibility detail supporting the main manuscript. Citations refer to the reference list at the end of this document.

### Annex A – Cortical-type ordinal scale, topological rules, and the reconciliation workflow

**Ordinal scale.** $\tau \in \{1, \ldots, 7\}$: allocortex (1), agranular (2), dysgranular (3), eulaminate I–III (4–6), koniocortex (7). Analyses that require a "complexity" coordinate use $\tau$ directly (an equal-spacing assumption, stated as such).

**R1 (sequential gradients).** For every mesh edge $(i, j) : |\tau(i) - \tau(j)| \leq 1$. Violations are enumerated as edges/clusters.

**R2 (concentric cortical-type rings).** Types 1 to 5 form concentric rings around the medial wall, nested from the peripheral allocortical ring inwards to the eulaminate-II matrix; only eulaminate III and koniocortex form islands (R4). Allocortex forms a single open arc, closing off-surface through the hippocampal formation, whereas agranular, dysgranular and eulaminate I each form a single connected component encircling the medial wall (a closed annular ring, first Betti number $\beta_1 = 1$). Assessed per grade by connected components, the Euler characteristic, and whether the component separates the medial-wall region from the rest (annularity) (see §3.2, §4).

**R3 (Eu-II matrix).** Eulaminate II forms a single connected component containing a fixed number of enclosed holes; assessed via connected components and the Euler characteristic / first Betti number of the patch.

**R4 (nested islands).** Eulaminate III and koniocortex each form isolated, non-annular clusters (three each in the released map). Because koniocortex (type 7) can under R1 border only eulaminate III (type 6), each koniocortex island is enclosed within an eulaminate-III island; in the released map no koniocortex vertex abuts the medial wall, and the three koniocortex islands distribute among the eulaminate-III islands, so the enclosure is not one-to-one. Naming them (RR22): the occipital eulaminate-III island encloses V1, the central and perisylvian island encloses both S1 and A1, and the ventrolateral prefrontal island (pars triangularis of the inferior frontal gyrus, von Economo's FDT) encloses none. The pattern is identical in both hemispheres.

**Correction workflow.** Rule clauses applied (García-Cabezas/Zaldívar-Díez tiers): entorhinal cortex and pre-/parasubiculum (modified hippocampal allocortex) → allocortex; piriform/periamygdaloid (olfactory paleocortex) → allocortex; perirhinal → mesocortex, assigned agranular or dysgranular per vertex to preserve the gradient; anterior parahippocampal → dysgranular; cingulate → agranular (anterior) / dysgranular (posterior); posterior parahippocampal kept eulaminate I; the hippocampal allocortex sensu stricto is declared off-surface. Presubiculum and parasubiculum use a parahippocampal-band proxy ($\leq 3$ hops from the medial wall, minus entorhinal), documented in the provenance file accompanying the data release. A topology-preserving, graded mesocortex buffer restores the strict gradient (R1 = 0) wherever a relabel would otherwise create a skip. The released map, together with its full per-vertex provenance and change log, is archived in the versioned data release (Zenodo DOI).

**Supplementary Table S1.** Reliability weights for the reference atlases combined into the weighted-consensus reference used to check and adjudicate the limbic belt (allocortex and mesocortex). Higher weight = more direct cytoarchitectonic evidence (ex-vivo histology > Glasser > gyral); values are defaults, not fitted.

| Source | Weight $r_s$ | Basis |
|---|---|---|
| Freesurfer *ex-vivo* histology (entorhinal+perirhinal/BA35-36) | 3 | cytoarchitectonic, *ex-vivo* ground-truth |
| Glasser HCP-MMP1.0 ({EC, PreS, PeEc, Pir, PHA1–3}) | 2 | Multimodal MRI |
| Destrieux (gyral) | 1 | macroanatomical / folding only |
| Desikan (gyral) | 1 | macroanatomical / folding only |

**Cost of enforcing the rules (audit).** The audit behind §3.2 classified every vertex changed between the topologically-clean starting map and the released v9 as topology-motivated or anatomy-motivated, by replaying each step. Of the 2,368 vertices changed solely to satisfy the sequential-gradient rule (967 LH, 1,401 RH), 84% are dysgranular and they concentrate in the peri-insular and superior-temporal belt: insula 8.2% of the parcel, superior temporal 6.3%, isthmus cingulate 8.2%, parahippocampal 5.1%, with 28 Desikan parcels touched at all (Supplementary Fig. S13). Counting anatomical relabels that the rule also constrained raises the total to 2,717 (0.89%), while the strictest reading of "changed solely for topology" gives 320 (0.10%), so the honest bracket is 320 to 2,717. Removing the most inclusive set with a two-hop mesh margin retains 57,181 of 58,731 vertices and leaves every primary correlation intact: T1w/T2w myelin +0.589 to +0.588, cortical thickness −0.502 to −0.495, the principal functional gradient −0.576 to −0.587 and AHBA gene expression PC1 +0.666 to +0.664, no shift larger than 0.011. All four still survive their family null. Two minor bookkeeping discrepancies in the intermediate version records, neither affecting the released map, are documented in `atlas/provenance/PROVENANCE_NOTES.md` of the data release.

**Resampling to 32k fs_LR.** The released 164k fsaverage annotation satisfies R1 with zero skip edges. The nearest-neighbour resampling to the 32k fs_LR mesh used for the structure–function analyses introduces three skip edges at triple junctions (LH vertex 1193, types 3 and 5; LH 11263 and RH 1470, types 4 and 6). These are a property of the coarser mesh, not of the released map, and they affect three vertices out of 59,412.

**Weighted belt reference (reconciliation aid).** This reliability-weighted consensus was used during map construction to adjudicate limbic-belt labels where the areal atlas is unreliable; it is a construction reference, not a benchmark that yields a reported agreement score. Sources $s$ with reliability $r_s$ (ex-vivo histology > Glasser > gyral). The consensus label at a vertex $v$ is the reliability-weighted plurality of the sources,

$$\updownarrow^*(v) \;=\; argmax_t \; \Sigma_s r_s \cdot 1[\; \updownarrow_s(v) = t \;]$$

where $\updownarrow_s(v)$ is the label source $s$ gives vertex $v$, $r_s$ its reliability weight, $t$ ranges over candidate types, and $1[\cdot]$ is the indicator (1 if true, else 0). Where sources disagree the more reliable one prevails; an allocortex-versus-isocortex call requires histology or Glasser support (gyral atlases corroborate but cannot override). The reliability weights are specified in Table S1.

**Topology audit of the released map.** Table S2 reports the audited values for the four rules on the released map; they are identical in the two hemispheres.

**Left–right agreement.** Because left–right symmetry was not enforced (each hemisphere was painted independently by the same rater under the same criteria; §2.1), we report the agreement between the two hemispheres as an internal consistency check. On the 32k fs_LR surface, where left and right vertices are homologous by construction, we compared the hemispheres vertex-by-vertex over the 29,286 vertices labelled (types 1–7) in both. They agree at 86.7% of vertices (Cohen's $\kappa = 0.83$, quadratic-weighted $\kappa_w = 0.96$), with disagreement overwhelmingly a single ordinal step (13.0% off by one type, 0.3% off by two or more; mean $|\Delta\tau| = 0.14$). The high weighted agreement indicates that the typing criteria were applied consistently across hemispheres, while the residual, near-diagonal disagreement concentrates at type borders and in the

**Supplementary Table S2.** Topological audit of the released cyto7 map. Values are identical in the left and right hemispheres. R1: number of skip edges (mesh edges with $|\Delta\tau| \geq 2$). R2: connected components of each low-complexity type and whether the component forms a single ring encircling the medial wall. R3: eulaminate II component count and number of enclosed holes. R4: number of isolated (non-annular) islands; each koniocortex island lies within an eulaminate-III island. The allocortex forms an open arc on the cut surface that closes off-surface through the hippocampal formation (§3.2, §4).

| Rule | Type | Audit result (LH = RH) |
|---|---|---|
| R1 (sequential gradient) | all types | 0 skip edges ($|\Delta\tau| \geq 2$) |
| R2 (core continuity) | allocortex | 1 component; open arc, closes off-surface |
| | agranular | 1 component; single ring encircling the medial wall |
| | dysgranular | 1 component; single ring encircling the medial wall |
| | eulaminate I | 1 component; single ring encircling the medial wall |
| R3 (eulaminate II matrix) | eulaminate II | 1 component enclosing exactly 3 holes |
| R4 (islands) | eulaminate III | 3 isolated (non-annular) islands |
| | koniocortex | 3 isolated (non-annular) islands |

limbic belt, where genuine interhemispheric asymmetries (notably the larger left entorhinal/medial-temporal allocortex; §3.1) are preserved rather than averaged away.

### Annex B – Benchmark metrics (cyto7 vs the area-level map)

**Confusion matrix** $O_{\text{ij}}$ = # *vertices* with cyto7 type $i$ and reference type $j$.

**Cohen's kappa (unweighted).** Chance-corrected agreement that treats every disagreement equally,

$$\kappa = \frac{p_o - p_e}{1 - p_e},$$

where $p_o = \sum_i O_{ii}/N$ is the observed agreement (the fraction of vertices assigned the same type by both maps) and $p_e = \sum_i O_{i\cdot}O_{\cdot i}/N^2$ is the agreement expected from the marginals. Unlike $\kappa_w$ below, it gives no partial credit for near-misses; the gap between the two ($\kappa = 0.36$ vs $\kappa_w = 0.70$) therefore reflects that most disagreements are a single ordinal step.

**Quadratically-weighted kappa.** With $K$ categories, weights $w\text{ij} = \frac{(i - j)^2}{(K - 1)^2}$,

$\kappa_w = 1 - (\Sigma_{\text{ij}}\ w_{\text{ij}}O_{\text{ij}})/(\Sigma_{\text{ij}}\ w_{\text{ij}}E_{\text{ij}})$, $E_{\text{ij}} = O_{i\cdot}O_{\cdot j}/N$

*where $O_{i\cdot}$ is the sum of all columns of row $i$ of the confusion matrix, and $O_{\cdot j}$ is the sum of all rows of column $j$.*

**Adjusted Rand index (ARI).** A label-free, chance-corrected measure of whether the two maps induce the same *partition* of the vertices: over all vertex pairs it counts agreement on co-membership (same vs different group), ignoring the type names, then subtracts the agreement expected by chance. It is computed from the same table $O$, with $C(x,\ 2) = x(x - 1)/2$ the number of pairs among $x$ items:

$$ARI = \frac{\Sigma_{ij}C(O_{ij},2) - ab/C(N,2)}{\frac{1}{2}(a+b) - ab/C(N,2)}$$

with $a = \Sigma_i C(O_{i\cdot},\ 2)$ and $b = \Sigma_j C(O_{\cdot j},\ 2)$. ARI = 1 for identical partitions, ≈ 0 at chance, and slightly negative if worse than chance. It is a separate metric from the weighted $\kappa$ above and complements it: $\kappa$ uses the type labels and their ordinal distance, whereas ARI ignores both and scores only the grouping (boundary) structure.

**Difference map.** Per vertex, signed ordinal difference $\Delta(v) = \tau_{cyto7}(v) - \tau_{\text{ref}}(v)$, categorized as agree / off-by-one (±) / off-by-≥2.

**Component analysis.** Connected components per type (mesh graph) with sizes and centroids; used to establish multi-focality (§3.2).

### Annex C – The anatomical support score

Components, each mapped to [0, 1]:

**Atlas concordance.**

$$c_{\text{atlas}}(v) = (\ \Sigma_s r_s(1 - |\tau(v) - \tau_s(v)|\ /\Delta_{\max})\ )/(\Sigma_s r_s)$$

i.e. reliability-weighted ordinal agreement with independent parcellations ($\Delta_{\max} = K - 1$).

*Implementation note. In the released build this general form is realised with a single reference for isocortex (the ordinal agreement 1 − |τ(v) − τ_vE(v)|/6 against the area-level type map) and, for the allocortex sliver, the fraction of two belt sources (FreeSurfer ex-vivo + Destrieux) marking the vertex as paleocortical belt. This uses fewer sources than the Annex B construction reference, which adds Glasser and Desikan at weights 3/2/1/1, and the difference is deliberate rather than pending. The two references answer different questions. Annex B's decides belt membership, where a wider consensus is an advantage. The concordance term measures how strongly a label is backed by sources that carry cytoarchitectural type, and Glasser, Destrieux and Desikan carry none, so counting them would raise the score without adding support of the kind the score is meant to record. Harmonising the two sets was tested rather than assumed: because the belt consensus exists only over the limbic belt and has no isocortical counterpart, it can reach just 6,421 of 304,972 vertices (2.1%), and rebuilding the term from the wider reference leaves the combined score correlated at 0.992 with the released one, moves no per-type median except allocortex, changes the disagreement AUC by less than 0.001, and moves no vertex between support tertiles. What it does do is raise the allocortex median from 0.32 to 0.56, and we retain the lower value because it is the honest one: the surface allocortex is typed on developmental grounds rather than from these atlases, and a support score should say so.*

**Topological consistency.**

On the mesh graph (vertices joined by triangle edges), the topology audit flags every vertex that participates in a rule violation: an R1 "skip" (a mesh edge with $|\tau(i) - \tau(j)| \geq 2$), an R2/R3 continuity-or-hole gap, or an R4 island violation; we call this the set of violating vertices. Let $d_{\text{viol}}(v)$ be the **graph-hop distance** (the number of edges in the shortest path between two nodes) from $v$ to the nearest violating vertex (the smallest number of mesh edges on any path between them, 0 at a violation) obtained by a multi-source breadth-first search seeded simultaneously from all violating vertices. The component is a fixed penalty within a tolerance of $h = 2$ hops:

$$c_{\text{topo}}(v) = \begin{cases} 0.5, & d_{\text{viol}}(v) \leq h \\ 1.0, & d_{\text{viol}}(v) > h \end{cases} \quad (h = 2 \text{ graph-hops})$$

i.e. any vertex within $h$ hops of a violation is down-weighted to 0.5, and every vertex farther away is treated as topologically clean (1.0). The distance is a count of edges, not millimetres (fsaverage vertices are ~1 mm apart, so 2 hops ≈ 2 mm). Because the released map has no R1 violations and clean R3/R4, the violating set is essentially empty and $c_{\text{topo}} \approx 1$ almost everywhere; this term protects against errors introduced by manual corrections on an intermediate/candidate map.

**Geometric reliability.** Two factors, each floored at 0.5 at the relevant edge and increasing linearly to 1.0 at a buffer distance. A medial-wall factor uses the geodesic distance $d_{\text{wall}}(v)$ (mm) from the medial wall (Dijkstra on the mesh, edge weights = Euclidean edge lengths); a boundary factor uses the graph-hop distance $h_{\text{bnd}}(v)$ to the nearest label boundary (any neighbour of a different type, medial wall included):

$$c_{\text{med}}(v) = clip(0.5 + 0.5d_{\text{wall}}(v)/D,\ 0.5,\ 1)$$

$$c_{\text{bnd}}(v) = clip(0.5 + 0.5h_{\text{bnd}}(v)/H,\ 0.5,\ 1)$$

$$c_{\text{geom}}(v) = min(c_{\text{med}}(v), c_{\text{bnd}}(v))$$

Where $D = 8\ mm$ and $H = 3\ hops$. Boundary- and wall-adjacent vertices are thus down-weighted to 0.5 and reach full reliability only $\geq$ 8 mm from the wall and $\geq$ 3 hops from any label edge, which is important for the thin, near-wall allocortex sliver.

**Provenance prior.** $c_{prior}$, a per-type base level (higher for expert-vetted isocortex, lower for the allocortex/agranular/dysgranular zone).

**Combination.** weighted geometric mean (main text §2.4, Box 2); weights are hyper-parameters reported with the release. A categorical high/medium/low map ($\geq$ 0.66 / $<$ 0.33) is provided for visualization. In the released build the provenance prior is allocortex 0.50, agranular 0.65, dysgranular 0.70, eulaminate I 0.90, eulaminate II 0.95, eulaminate III 0.95, koniocortex 0.90, and the four component weights are equal (all 1). Candidate additional components (label heterogeneity, curvature, and a borrowed Jülich cross-subject variability prior) were evaluated under a pre-specified keep-rule, and none improved calibration, so the released map retains these four components (see §3.4).

**Anti-circularity.** No functional/microstructural term enters the score; a separate descriptive T1w/T2w-agreement overlay is provided but is not part of the support score used to weight analyses.

**How well does the score predict disagreement?** We also asked how well the support score predicts *where* cyto7 and the area-level reference disagree, using only the components that do not depend on that benchmark (topology, geometry, provenance). It does so modestly (AUC $\approx$ 0.58), with the signal carried almost entirely by the geometric term (the topology term is uniformly high, because the released map is topologically clean). Read plainly, the support score is a transparent, mostly geometric reliability indicator (vertices near type borders and the medial wall are the least certain) rather than a finely calibrated error model, and its four components are not statistically independent of one another (the topology term is near constant by construction, because the map was repaired until the rules held). On a small ex-vivo histological patch (509 belt vertices), labelling error decreases monotonically with the score across its three tertiles: 0.153, 0.071 and 0.019, on 170, 126 and 213 vertices, which is 26, 9 and 4 mislabelled vertices out of 39 in total. This is an internal consistency check and not an external test, because the belt was partly built from these labels. The counts are small, the vertices are spatially clustered rather than 509 independent observations, and a few hundred contiguous vertices from one specimen cannot calibrate an error model, so the ordering is the only thing we read from it. The components are combined with each floored at 0.02 before the logarithm, which bounds the penalty a single zero component can impose.

**Candidate components tested and rejected.** To check whether the support score could be improved, we tested three candidate components under the pre-specified rule of §2.4: a candidate is adopted only if it raises the benchmark-independent disagreement AUC without duplicating an existing term. None qualified: local label heterogeneity was redundant with the geometric boundary term, cortical curvature was independent but degraded calibration, and the borrowed Jülich cross-subject variability prior did not predict disagreement at all (AUC $\approx$ 0.49, i.e. chance). The released map therefore keeps its four anatomy-only components (disagreement AUC $\approx$ 0.58). We note, as a forward-looking point, that a properly *external, per-subject* variability estimate (rather than the borrowed group prior tested here) may prove a promising future refinement.

### Annex D – Spatial statistics (spin test + FDR)

**Surface resampling.** All features and the cyto7 map itself were brought to the 32k fs_LR mesh with the `neuromaps fsaverage_to_fslr` transform (which calls Connectome Workbench `wb_command` with the standard fsaverage and fs_LR registration spheres). Continuous features use barycentric interpolation; the categorical type map uses nearest-neighbour resampling, since interpolating ordinal labels would create types that were never painted. Round-trip agreement per type (Dice, LH / RH) is: allocortex 0.65 / 0.70, agranular 0.81 / 0.85, dysgranular 0.964 / 0.964, eulaminate I 0.981 / 0.980, eulaminate II 0.984 / 0.984, eulaminate III 0.988 / 0.987, koniocortex 0.976 / 0.978. The two least well preserved types are the two smallest, which is expected when a thin band is carried to a four-times-coarser mesh, but they are also the types carrying the map's distinctive claims, so the structure–function results of §3.5 should be read as validating the resampled map rather than the released 164k one. Three R1 skip edges introduced by this step are documented in Annex A.

**Spatial-null convention.** One spatial-null convention is used throughout: the plus-one corrected two-tailed tail, $p = (1 + \#\{|\rho_{\text{null}}| \geq |\rho_{\text{obs}}|\})/(N + 1)$ with $N = 1000$, so the smallest attainable value is $1/1001 \approx 0.001$. Vertex-level tests draw their null from 1,000 Alexander-Bloch rotations of the type map. Parcel-level tests (the layer-marker genes of §3.7 and the disease-vulnerability maps of Annex H) draw theirs from 1,000 rotations of the 68 Desikan–Killiany parcel centroids generated with the ENIGMA toolbox's `rotate_parcellation` (`numpy` seed 0), rotating the per-parcel type score and recomputing the correlation at each rotation. The two differ only in what is rotated, so all reported spin $p$-values are on the same scale and are directly comparable.

**Sign conventions for principal components and gradients.** The sign of a principal component or of a diffusion-embedding gradient is mathematically arbitrary, so orientation has to be fixed without reference to the type map. AHBA gene-expression PC1 and the principal functional-connectivity gradient are used at the orientation published in their source distributions (`abagen` and Margulies et al. [2016] respectively, both via `neuromaps`) and were not reoriented here. Receptor PC1 is assembled in this work rather than taken published, so its sign is anchored to correlate positively with the across-tracer mean density, a neutral reference independent of cyto7. In no case was a sign selected by reference to the cyto7 map, and the predicted directions in Box 3 were fixed before testing.

**Spin null.** The cortical-type map is rotated on the sphere via the registration spheres; for each of $N = 1000$ rotations the Spearman $\rho$ against each feature is recomputed, giving a spatially-constrained null. $p_{spin}$ as in §2.6. The same rotations serve all features.

**FDR.** Benjamini–Hochberg across the feature family: reject $H_{(i)}$ for the largest $i$ with $p_{(i)} \leq (i/m)\alpha$. *Note: robustness variants of a single metric (e.g. alternate spectral-centre choices) are treated as one family member, not as independent tests.*

**Feature summary.** Table S3 collects, for every structural, functional and external-reference feature, the per-type median and the vertex-level Spearman statistics from this spin+FDR pipeline (applied throughout Annexes E–H).

Supplementary Table S3. Structure–function and external-reference summary by cyto7 type. For each feature: the per-type median (hemisphere-averaged) across the seven cyto7 types, then the vertex-level Spearman correlation $\rho$ with the ordinal type, its spin-test $p$ ($N = 1000$ rotations) and Benjamini–Hochberg FDR $q$ over the feature family. $\rho$, $p$ and $q$ are computed over all cortical vertices of both hemispheres; $*$ marks features surviving the FDR-controlled spin null ($q < 0.05$). Myelin, thickness and the functional gradient survive; in this group-average panel the five raw MEG band powers, receptor PC1 and BigBrain skewness do not. The intrinsic-timescale row reports the single-subject, source-reconstructed values (medians and $\rho = -0.46$, $q = 0.014$; Annex F), which survive the spin+FDR null; its $\rho$ is a per-subject mean, not a vertex-level coefficient like the other rows. The aperiodic-corrected $\beta$ power that tracks type in Annex F is distinct from the raw band powers shown here. AHBA gene-expression PC1 also survives but is interpreted as a fourth (transcriptomic) readout of the same gradient of laminar complexity rather than an independent test (Annex H). Units: myelin, T1w/T2w ratio; thickness, mm; timescale, ms; band powers, relative fraction; functional gradient and PC scores, arbitrary units.

| Feature | Allo | Agr | Dys | EulI | EulII | EulIII | Konio | $\rho$ | spin $p$ | $q$ |
|---|---|---|---|---|---|---|---|---|---|---|
| T1w/T2w myelin | +1.28 | +1.15 | +1.20 | +1.25 | +1.29 | +1.38 | +1.55 | +0.59 | 0.001 | 0.004* |
| Cortical thickness | 2.99 | 3.12 | 2.96 | 2.74 | 2.62 | 2.52 | 1.97 | −0.50 | 0.002 | 0.006* |
| Functional gradient | +0.25 | +0.58 | +2.52 | +4.21 | 0.00 | −4.22 | −4.68 | −0.58 | 0.001 | 0.004* |

| Feature | Allo | Agr | Dys | EulI | EulII | EulIII | Konio | $\rho$ | spin $p$ | $q$ |
|---|---|---|---|---|---|---|---|---|---|---|
| Intrinsic timescale (ms) | 41.5 | 46.8 | 40.9 | 34.7 | 29.5 | 24.6 | 21.9 | −0.46 | 0.004 | 0.014* |
| Delta power | 0.340 | 0.357 | 0.350 | 0.334 | 0.312 | 0.291 | 0.282 | −0.44 | 0.110 | 0.198 |
| Theta power | 0.236 | 0.247 | 0.259 | 0.246 | 0.257 | 0.244 | 0.252 | −0.18 | 0.623 | 0.623 |
| Alpha power | 0.287 | 0.247 | 0.257 | 0.289 | 0.266 | 0.330 | 0.318 | +0.37 | 0.257 | 0.371 |
| Beta power | 0.082 | 0.085 | 0.095 | 0.097 | 0.102 | 0.101 | 0.113 | +0.27 | 0.416 | 0.468 |
| Gamma power | 0.023 | 0.024 | 0.025 | 0.023 | 0.024 | 0.020 | 0.021 | −0.31 | 0.289 | 0.371 |
| AHBA gene expression PC1 | −1.78 | −1.77 | −1.24 | −0.45 | −0.14 | +0.49 | +1.24 | +0.67 | 0.001 | 0.002* |
| PET receptor PC1 | −0.26 | +1.34 | +1.80 | +1.40 | +1.07 | +0.46 | −0.44 | −0.33 | 0.063 | 0.063 |
| BigBrain profile skewness | +1.96 | +1.85 | +1.90 | +1.83 | +1.84 | +1.69 | +1.50 | −0.21 | 0.321 | 0.401 |

**FDR families.** Every confirmatory and exploratory test in this study belongs to exactly one family, and families are never pooled. Families were defined by measurement domain and statistical unit, before the results were seen, and the listing below is complete for them: 147 entries in 21 families, of which 117 are tests and 30 are descriptive entries carrying no null and no test: the 27 per-feature win fractions on the disagreement set and the three septile-comparator win fractions. Table S8's Status column marks each. Some robustness analyses belong to a declared family and are listed in Table S8 with the status *robustness*. The additional robustness and sensitivity analyses listed in Table S9 sit outside these families and receive no correction, because they ask whether a result is stable rather than whether an effect is present; they are listed separately so that what was run is on the record without being counted as inference. One exception is documented there: the comparator correlations of Annex J were given Benjamini–Hochberg control within each partition, and their $q$ values are released in `comparator_panel.csv`. Benjamini–Hochberg correction is applied within a family whenever the members are distinct hypotheses about distinct measures. It is not applied in four cases, for one stated reason in each. The three aggregate added-value variants (all vertices, off-by-one, off-by-two-or-more) are nested re-expressions of a single statistic on nested subsets, so correcting across them would correct a statistic against itself; the 27 descriptive per-feature win fractions are the per-arbiter decomposition of those aggregates, reported for transparency rather than as separate claims, and the confirmatory claim is the aggregate; the five tractography tests and the eight per-type tractography tests carry no inferential claim at all (Annex G); and the robustness families are sensitivity analyses rather than new hypotheses: most re-run a test already corrected in its own family, while `benchmark_controls` bears on the §3.3 benchmark itself rather than on the type axis, so pooling it with the type-axis families would be a category error. Where no correction is applied the table reports the raw $p$ and no $q$, and Table S8 marks the row accordingly. "Hypothesis status" repeats Table S8's Status column and records whether a test was specified in advance, not whether its result is retained.

**Supplementary Table S4.** FDR families. Every family, its size, whether Benjamini–Hochberg correction was applied within it, and the hypothesis status of its members. Sizes sum to the 147 entries of Table S8. `layer_markers` is mixed because the five single-gene tests were pre-specified and the three z-composites were not. `bigbrain_profile_panel` is likewise mixed: profile skewness was the pre-specified index and the other four features are exploratory. It is listed separately from `external_reference` because its five members were corrected together, which is what gives profile skewness $q = 0.401$ from $p = 0.321$; `external_reference` accordingly holds two members, not three. `osc_band_as_coded` has four members, not three or five, because it retains all four non-$\beta$ bands under their coded edges: $\theta$ and $\alpha$, which are estimable, and $\delta$ and $\gamma$, which are not and are reported without being interpreted (Annex F). $\beta$ is carried separately as the pre-specified single-member test in `dynamics_singlemember`.

| Family | $n$ | FDR applied | Hypothesis status |
|---|---|---|---|
| `structure_function` | 9 | yes | pre-specified |
| `added_value_global` | 9 | yes | pre-specified |
| `added_value_aggregate` | 3 | no | pre-specified |
| `added_value_localized` | 27 | no | pre-specified |
| `bigbrain_added_value` | 2 | yes | pre-specified |
| `bigbrain_profile_panel` | 5 | yes | mixed |
| `dynamics_singlemember` | 7 | yes | pre-specified |
| `external_reference` | 2 | yes | pre-specified |
| `fig9_predictions` | 3 | yes | pre-specified |

| Family | $n$ | FDR applied | Hypothesis status |
|---|---|---|---|
| `negative_control` | 2 | yes | pre-specified |
| `tractography` | 5 | no | pre-specified |
| `layer_markers` | 8 | yes | mixed |
| `osc_band_as_coded` | 4 | yes | exploratory |
| `receptor_maps` | 19 | yes | exploratory |
| `receptor_composites` | 8 | yes | exploratory |
| `disease` | 8 | yes | exploratory |
| `tractography_per_type` | 8 | no | exploratory |
| `disease_partial` | 8 | yes | robustness |
| `dynamics_robustness_variants` | 5 | no | robustness |
| `single robustness variant` | 1 | no | robustness |
| `benchmark_controls` | 4 | no | robustness |

### Annex E – Myelin progression regression

Per-vertex T1w/T2w values are assembled per type after a validity mask (myelin $> 0$), an optional gyral-crown restriction (curv $\leq 0$), a geodesic medial-wall buffer, and targeted tail-trimming (the same constrained as described by García-Cabezas et al. [2020], Fig. 8D).

**Type-median fit.** $\langle m \rangle_\tau \;=\; a\,\tau \;+\; b$ over $\tau \in \{\textit{dysgranular},\ \ldots,\ \textit{koniocortex}\}$; $R^2_{\text{med}}$.

**Per-vertex fit.** the same over all retained vertices; $R^2_{\text{vtx}}$. The two answer different questions and are not comparable (the per-median value removes within-type variance by construction).

Agranular and allocortex are excluded from the fit (motor and limbic myelin variance does not follow the sensory progression that the gradient of laminar complexity captures).

### Annex F – Source-reconstructed MEG dynamics and the Mahjoory relation

**Source reconstruction.** We took the preprocessed resting-state sensor MEG of the HCP young-adult cohort (Shafiei et al., 2022; $N = 89$, three runs each), projected it to cortical sources with an LCMV beamformer on a realistic single-shell (FieldTrip) forward model, and computed a Welch power spectrum per source. Each spectrum was fitted with `specparam` over 2–40 Hz (fixed aperiodic mode, pre-specified; knee and broadband 1–100 Hz variants as robustness checks), separating the aperiodic 1/f component (exponent, offset) from oscillatory peaks. From the broadband source signal we computed an intrinsic timescale as the autocorrelation decay (area under the ACF to the first zero crossing, primary; 1/e-lag and exponential-decay variants; and a knee-derived timescale $\tau_c = 1/(2\pi f_{\text{knee}})$). The power-weighted spectral centroid is

$$C(v) \;=\; (\ \Sigma_b\ P_b(v)\, f_b\ )\ /\ (\ \Sigma_b P_b(v)\ )$$

over bands $b \in \{\delta,\ \theta,\ \alpha,\ \beta,\ \gamma 1\}$ with centre frequencies $f_b = \{2.25,\ 6,\ 10,\ 21,\ 45\}$ Hz (band midpoints; log-spaced and geometric variants as a robustness check), and the pre-specified slow/fast ratio is $(\delta+\theta)/(\beta+\gamma)$. Fits were good (99.9% of sources $R^2 \geq 0.9$) and reproducible across runs (across-run reliability: exponent 0.87, intrinsic timescale 0.83–0.88). The reconstruction was validated against the neuromaps group maps: source $\alpha$ power matches *megalpha* at $\rho = 0.98$, and the broadband timescale matches *megtimescale* at $\rho = 0.85$.

**Result.** Each metric was tested against ordinal cyto7 type with the released spin+FDR machinery (Annex D), in its own dynamics family. The intrinsic timescale decreases toward koniocortex and survives: $\rho = -0.46$ (spin $p = 0.004$, $q = 0.014$) for the pre-specified area-under-the-ACF definition, which enters the dynamics FDR family as the single timescale member (§2.6). That coding is the operative one throughout, and is what Table S4, Table S8 and Figures 5, S1 and S2 all use; $q = 0.014$ reproduces from the seven raw $p$-values of that family in Table S8. Note also that a second, distinct test shares the name "intrinsic timescale" in Table S8: the vertex-level structure-function member, which does not survive ($q = 0.103$). The surviving result is the subject-level area-under-the-ACF measure reported here. The 1/e-lag ($-0.32$, spin $p = 0.025$), exponential-decay ($-0.38$, $p = 0.013$) and knee-derived ($-0.34$, $p = 0.027$) definitions are robustness checks reported outside the family; all agree in direction, so the gradient does not depend on the choice of definition. The spectral centroid increases ($\rho = +0.41$, $q = 0.014$), the slow/fast ratio decreases ($\rho = -0.36$, $q = 0.021$), and aperiodic-corrected $\beta$ power increases ($\rho = +0.22$, $q = 0.021$). The aperiodic exponent does not track type ($\rho = -0.12$, spin $p = 0.50$); it is also unstable across fitting modes (knee $+0.34$, spin $p = 0.040$; broadband $+0.10$). Both are robustness variants outside any FDR family, so neither is a corrected result and "surviving" has no defined meaning for them; what the knee value shows is that the sign reverses with the fitting mode. We read that as sensitivity to specification rather than as support for the null. Peak frequency and the offset are also null.

**Statistical unit and null.** The reported $\rho$ is the mean over the 89 subjects of each subject's vertex-level Spearman correlation between the metric and ordinal type (median $-0.471$ for the primary timescale, against a mean of $-0.459$). The three runs per subject are averaged within subject before correlating. The null is hierarchical: for each of the 1000 rotations the same rotated type map is applied to every subject, each subject's $\rho$ is recomputed, the group mean is taken, and $p$ is the plus-one-corrected fraction of rotations whose absolute group mean equals or exceeds the observed one. Subject-level consistency is reported beside the group statistic: 89 of 89 subjects show the predicted sign for the intrinsic timescale, 88 of 89 for the spectral centroid, 87 of 89 for the slow/fast ratio and 80 of 89 for aperiodic-corrected $\beta$ power. Excluding the allocortex leaves the primary timescale at $-0.451$ (spin $p = 0.005$).

**Aperiodic-corrected $\delta$ is not estimable, and $\gamma$ is poorly estimated.** With `specparam` fitted over 2–40 Hz, a $\delta$ peak sits at the edge of the fit range and $\gamma$ (30–60 Hz) mostly outside it: $\delta$ is non-zero at 0.2% of subject-vertices and constant within 56 of 89 subjects, whereas $\gamma$ is non-zero at 3.0% and constant within 4. Their tests appear in Table S8 in the `osc_band_as_coded` family, computed with the band edges as coded, but they are not interpretable as nulls: a non-significant result for a quantity that is zero at almost every subject-vertex carries no information about type. The band-power controls below are fitted with all five bands in the design matrix, as Table S9 records. We do not claim this is equivalent to a control on $\theta$, $\alpha$ and $\beta$ alone. A pseudo-inverse discards directions by numerical tolerance, not by how often a column is zero, and $\gamma$ in particular is constant within only 4 of the 89 subjects, so it can influence the fit for most of them. What the sparsity means is that the $\delta$ and $\gamma$ columns are poorly estimated, not that they are absent from the fit. A control restricted to the three reliable bands is a different model and we have not run it, so robustness to the band specification is untested.

**Control for shared spectral structure.** Because the timescale, centroid, slow/fast ratio and aperiodic slope are related summaries of one spectrum, the type-to-timescale association was retested with the others partialled out, per subject, under the same hierarchical spin null. It survives partialling the aperiodic exponent ($-0.436$, nominal $p = 0.010$), all five coded band powers ($-0.348$, $p = 0.002$) and both together ($-0.284$, $p = 0.009$, 88 of 89 subjects), a 38% attenuation from the uncontrolled $-0.459$. The reverse does not hold: with the timescale partialled out, the exponent's association with type weakens further ($-0.047$, $p = 0.817$). These $p$-values are nominal and are not FDR-controlled within a declared family (Table S9). Two limits apply. The covariate set includes $\delta$ and $\gamma$, which are poorly estimated under this fit, and the model restricted to the three reliable bands has not been run, so robustness to the band specification is untested. Variance inflation is below 1.5 for every covariate in the full model, computed over the subjects in which that covariate varies.

**Relation to Mahjoory et al. [2020].** They report a peak-frequency gradient by a linear model against geodesic distance, without a spin null. Our per-subject, source-reconstructed analysis reproduces the direction and resolves it on the cytoarchitectural axis under a spatially constrained spin null. An earlier pass on the group-average neuromaps maps with a single-sphere forward found only a directional trend (timescale $\rho \approx -0.48$, not surviving; the group-average nine-feature panel of Table S3); the realistic forward and per-subject spectra here (source $\alpha$ vs *megalpha* $\rho$: $0.51 \rightarrow 0.98$) recover the effect, so the absence of a detected exponent association is not attributable to the forward model. Per-metric statistics with the single-sphere comparison are in the released `meg_dynamics_v2_summary` table.

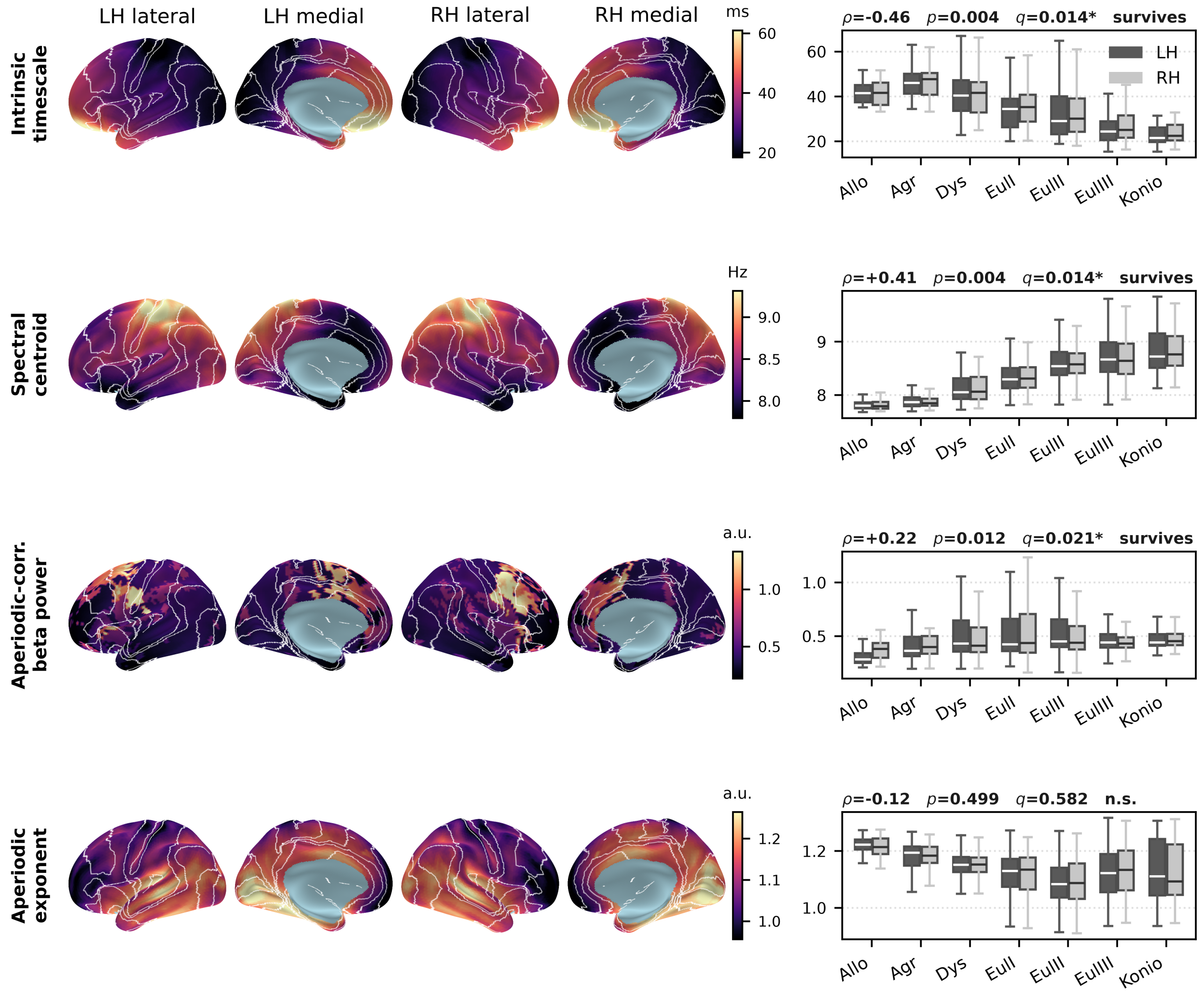


**Supplementary Figure S1. Resting-state MEG dynamics along the cyto7 axis (source-reconstructed).** Group-mean maps of the intrinsic timescale, spectral centroid, aperiodic-corrected $\beta$ power and the aperiodic 1/f exponent on the inflated 32k fs_LR surface (both hemispheres, lateral and medial; magma) with cyto7 type borders, alongside per-type box-and-whisker distributions annotated with the group mean of the per-subject Spearman $\rho$, the spin $p$ from the hierarchical subject-by-rotation null, and FDR $q$. The intrinsic timescale ($\rho = -0.46$), spectral centroid (+0.41) and $\beta$ power (+0.22) survive the spin+FDR null; the aperiodic exponent (−0.12) does not (§3.6, Annex F).

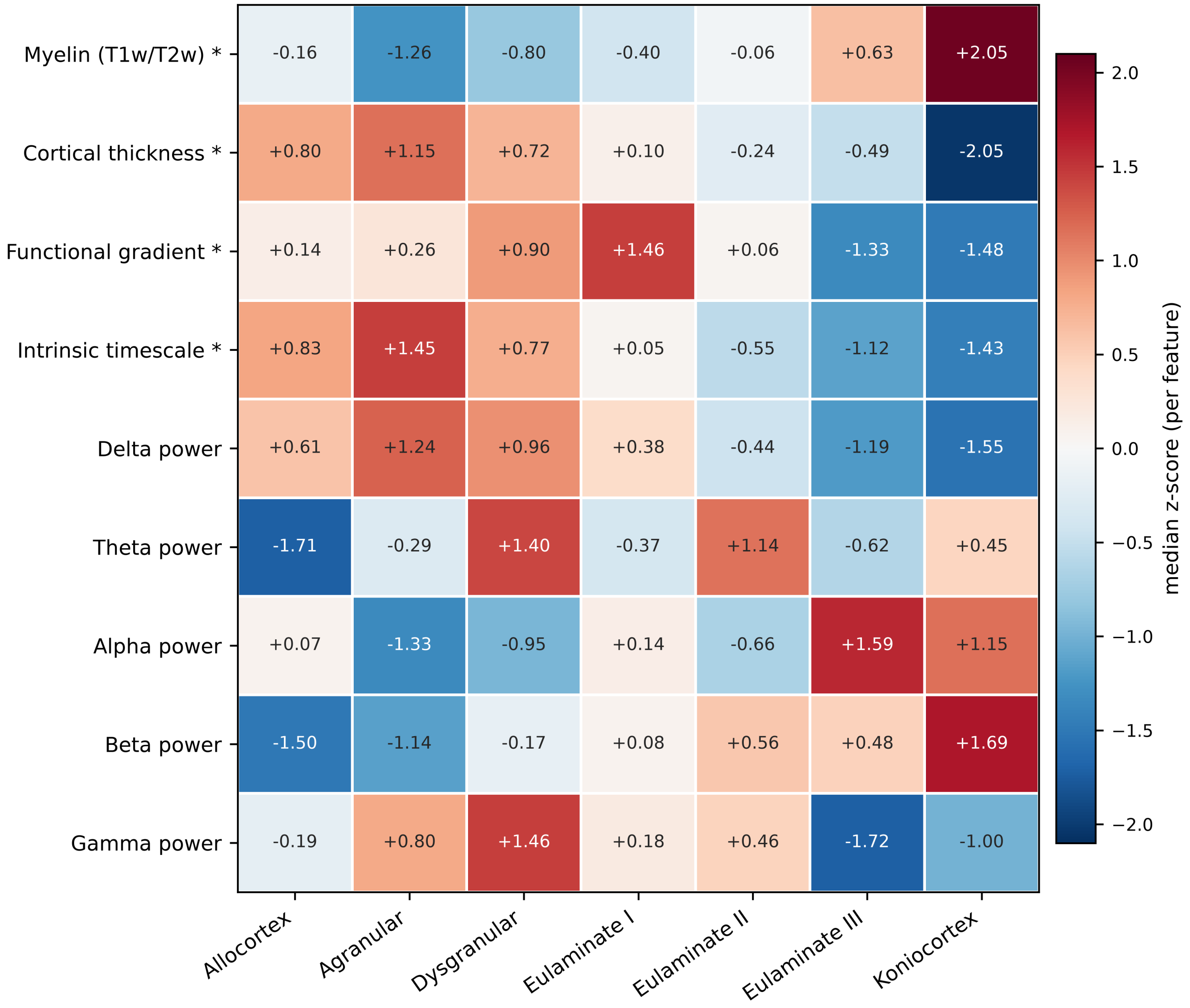


**Supplementary Figure S2. Feature fingerprint across cortical types.** Median of each structural and functional feature within each cyto7 type, z-scored across the seven types so that features on different native scales are comparable (red: above the feature's cross-type average, blue: below). Rows: intracortical myelin and cortical thickness, the principal functional gradient, then the intrinsic timescale and the MEG band powers; $*$ marks the features that survive the spin null with FDR control (myelin, thickness, the functional gradient and the intrinsic timescale). The intrinsic-timescale row shows the single-subject, source-reconstructed values (Annex F), which survive the null; the five MEG band powers remain the group-average maps. The colour encodes the z-scored median only; the reported correlations (main text and Fig. 5) are vertex-level Spearman coefficients over all cortical vertices under a spatial-autocorrelation null (a per-subject mean for the timescale), not fits to these seven medians.

### Annex G – Tractographic connectivity vs type-distance

cyto7 surface labels were projected to an fsaverage volumetric label map with FreeSurfer `mri_aparc2aseg` (cortical labels 1000+i [LH]/2000+i [RH], remapped to 1–7) and warped into the HCP-1065 tractography reference space with `mri_robust_register` + `mri_vol2vol --lta` (nearest-neighbour), giving a 0–7 label volume in the same RAS+mm frame as the streamlines. Connectivity was computed in DIPY by **endpoint** assignment (a streamline contributes to a type pair only when its two endpoints fall in those two regions); per-pair bundle geometry (endpoint membership, $\geq 5$ streamlines) gives mean length (arc length), tortuosity (arc length $\div$ end-to-end distance) and mean absolute direction along the RAS axes. Bundles are split at 80 mm into short/long range. That cutoff is deliberately high: population-averaged tractograms under-recover genuine short-association (U-)fibres, so a conventional cutoff would leave the short-range class dominated by the connections the template reconstructs least well.

**Type-distance falloff.** Splitting the type-by-type streamline counts by bundle length (Fig. S11a,b, shared log scale) shows short-range connections concentrating near the matrix diagonal, between cytoarchitecturally similar types, whereas long-range bundles are denser and spread more broadly across type-distances. For the 21 off-diagonal pairs, $\log(1+w_{\tau\tau'}) = \alpha\,|\tau-\tau'| + \beta$; short-range slope $\alpha \approx -1.23$ (label-permutation p = 0.003; Fig. S11c), about a threefold reduction per ordinal step. The $\log(1+w)$ form and the retention of empty cells both matter, and are not cosmetic: six of the 21 pairs carry no short-range streamlines at all, and they lie at the largest type-distances, so they are the evidence that connectivity decreases to nothing rather than nuisance zeros to be dropped. Fitting $\log w$ on the non-empty cells only, or including the seven diagonal cells, gives $-0.69$ and $-0.41$ respectively; the released matrix is `connectivity_short.csv` and the mapping from each quantity below to the file that reproduces it is in the release (`results/tables/tractography/PROVENANCE.md`). This result is exploratory. The label permutation reassigns type labels and so destroys the concentric spatial arrangement the rules impose, which makes it an easy null; replacing it with a rotation-based null that preserves that arrangement gives $p = 0.040$ for the identical statistic, against a null distribution centred at Pearson $r = -0.334$, so about a third of the observed $-0.765$ follows from the geometry of the map alone.

**Spatial-opportunity control.** Because short streamlines preferentially connect nearby cortex, the offset $\log(n_{\text{src}} \times n_{\text{tgt}})$ over-counts: it credits every possible endpoint pair, including pairs no short streamline could join. We therefore counted, on the fsaverage white surface in both hemispheres and from the released v9 annot, the vertex pairs whose three-dimensional Euclidean separation is at most 80 mm, the same cutoff that defines short range (exact dual-tree counting, no sampling; 40 mm and 120 mm reported as sensitivity). Euclidean separation is used because a streamline runs through white matter rather than along the cortical sheet. Opportunity is not proportional to the number of possible pairs: the fraction within reach ranges from 37% to 99% across type pairs and decreases with type-distance ($r = -0.65$ on the log fraction). Substituting it as the offset, the aggregate type-distance coefficient is $-1.15$ ($p = 1.7 \times 10^{-6}$, spin $p = 0.067$) against $-1.36$ (spin $p = 0.060$) under the old offset, and the published statistic re-expressed as a connection rate per available pair attenuates from a slope of $-1.23$ to $-0.80$ per ordinal step ($r = -0.765$ to $-0.597$) and no longer survives the rotation null ($p = 0.070$). The 21 off-diagonal cells are also not 21 independent observations: they summarise 33 named template bundles, and modelled at the bundle level the coefficient remains zero under either offset ($+0.01$, $p = 0.96$; $+0.13$, $p = 0.43$). Two specifications that add contact area on top of the opportunity offset do clear the null ($-1.47$, spin $p = 0.021$; $-1.68$, spin $p = 0.012$), but contact area is non-zero for only 6 of the 21 pairs and so acts largely as an adjacency indicator, the second coefficient is not identified in its own model ($p = 0.117$, VIF 8.8), and what changes between these specifications is chiefly the null mean ($-0.60$ to $-0.37$ to $+0.01$) rather than the observed effect. We therefore report the falloff as an ordinal, exploratory observation attributable to the spatial construction of the map, and draw no inferential claim from it.

**Proximity control.** The geodesic control reported previously measured centroid-to-centroid distance between type footprints, which is invalid for this map because most types are annular or multi-focal, so a type's centroid falls in its own hole: it placed agranular and dysgranular, which share 4,728 mm$^2$ of border, 54.3 mm apart, further than dysgranular is from koniocortex, with which it shares none. Replacing it with area-weighted nearest-neighbour distance between footprints (Euclidean and geodesic, median and 10th percentile) compresses the six bordering type pairs from a 4.5 to 54.3 mm spread to 4.9 to 9.5 mm, as a valid measure must. It also strengthens rather than weakens the collinearity: type-distance correlates with the valid proximity measures at $r = +0.92$ to $+0.97$ across the 21 pairs, against $r = +0.81$ for the superseded centroid measure. With seven ordinal types, type-distance and physical proximity are close to the same predictor and cannot be separated on these data.

**Boundary-surface control.** Inter-region contact area $A_{\tau\tau'}$ (shared voxel faces $\times$ face area) correlates with short-range connectivity (Pearson $r \approx 0.58$); after partialling contact area out, the type-distance effect remains (partial $r = -0.64$, $p = 0.002$). This control is reported for completeness but is not sufficient: sharing a border is not the same as being close, and the proximity and spatial-opportunity controls above are the ones that decide the question. On its own, the contact-area adjustment does not license an inferential claim, and none is drawn.

**Per-type connectivity.** Summarising the streamline matrices by type rather than by type-pair, and normalising each type's incident streamline count by its surface vertex count, the per-type short-range degree decreases from the limbic belt toward koniocortex (Spearman $\rho \approx -0.89$ across the seven types; short and long combined $\rho \approx -0.82$; label-permutation $p = 0.01$–$0.03$), while the mean bundle length and tortuosity increase ($\rho \approx +0.96$ and $+1.00$; Fig. S3). The raw, unnormalised degree shows no trend, so the effect is one of connection density rather than region size. The direction is consistent with the per-type T1w/T2w myelin medians (Table S3), but streamline counts are a connectivity-density proxy and not myelination (see Caveats), so this is reported as a connectional-property gradient rather than a claim about myelinated fibres.

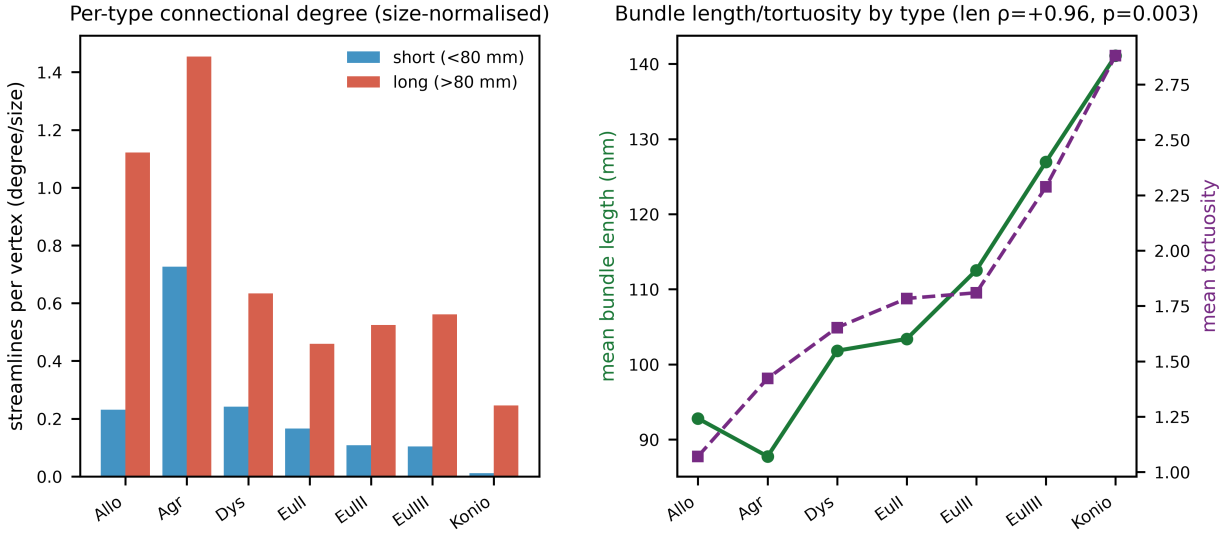


**Supplementary Figure S3. Per-type cortico-cortical connectivity.** Summaries of the HCP-1065 template streamline matrices by cyto7 type. After normalising each type's incident streamline count by its surface vertex count, the short-range degree decreases from the limbic belt toward koniocortex, while the mean bundle length and tortuosity increase; the raw, unnormalised degree shows no trend, so the effect reflects connection density rather than region size. Streamline counts are a connectivity-density proxy and not myelination, and the direction is cross-referenced to the per-type T1w/T2w myelin medians (Table S3).

**What the data support.** An ordinal, descriptive statement: short-range streamline counts are highest between cytoarchitecturally similar types and decrease with cytoarchitectural type-distance. That is consistent with the architectonic-type principle, whereby connection probability declines with cytoarchitectural dissimilarity [Barbas and Rempel-Clower, 1997, Beul et al., 2017, Hilgetag et al., 2019], rather than a test of it, and it is read with the caveats of the group-averaged template and the surface-to-volume projection set out in §4.9 of the main text. The per-type gradient above is reported on the same footing.

**Caveats.** Streamline counts are not axonal density; gyral/length biases and template-space projection apply; the result is a relative, ordinal prediction, computed on the population-averaged HCP-1065 template tractogram fitted to the released map.

### Annex H – External validation: molecular and histological references

Several references held entirely external to the atlas were tested against the ordinal cyto7 type through the same 32k fs_LR resampling and spin+FDR pipeline (Annex D); the transcriptomic, receptor and histological references are described first, and receptor diversity, cortical expansion and disease vulnerability follow. Per-type medians and statistics for the primary features are included in Table S3.

**Transcriptomic gradient (AHBA gene-expression PC1).** First principal component of Allen Human Brain Atlas cortical microarray expression (six donors in the source dataset, five analysed here), assembled with `abagen` [Arnatkeviciute et al., 2019]. $\rho$ vs type $= +0.67$ (spin $p = 0.001$, FDR $q = 0.002$): this survives the spin null, and the per-type medians increase monotonically (allocortex $-1.78 \rightarrow$ koniocortex $+1.24$). It correlates $r \approx 0.60$ with the functional gradient, so it is reported as a fourth (transcriptomic) readout of the one dominant sensorimotor–association axis, not an independent test. Caveats. Five of the six donors contribute; donor 15496 was excluded because its data are no longer retrievable from the Allen distribution URL the pipeline uses. Processing was `abagen` with bidirectional left–right mirroring, interpolation of unsampled regions, and `norm_matched` disabled; probe selection was left at the `abagen` default. The same configuration produced both the PC1 map and the individual marker genes. Mirroring copies samples across the midline and interpolation estimates expression where no sample exists, so both improve coverage without adding independent tissue: the 68 parcel values are not 68 independent observations, and a null for a hemisphere-pooled predictor against mirrored expression should respect that bilateral dependence. A left-hemisphere-only and a leave-one-donor-out sensitivity analysis would address this directly and are not yet run.

**Neurotransmitter receptors and transporters (PET).** The 19 receptor and transporter density maps of the Hansen collection [Hansen et al., 2022] were tested against cyto7 type both as a single principal component and map by map, through the same 32k fs_LR resampling and spin+FDR pipeline (Annex D), with cyto7 as the type map. The first principal component (PC1 $\approx$ 50% variance, assembled from five native-fsaverage maps and 14 MNI152 volumes projected to fs_LR) shows only a weak trend ($\rho = -0.33$, spin $p = 0.063$) that does not survive the null. Individual receptor systems do track the type axis, in both directions. We do not claim that this is why PC1 is flat: a principal component carries signed loadings and can represent opposing patterns, so the explanation would need the loadings themselves, which we do not report. The sign of a principal component is in any case arbitrary. Examined individually (Benjamini–Hochberg FDR across the 19-map family, kept separate from the structure–function and external-validation families so the other $q$-values are unchanged), four maps survive, and the directions are specific: the $\mu$-opioid receptor (MOR, $\rho = -0.55$), serotonin 5-$HT_{1A}$ ($\rho = -0.54$) and cannabinoid $CB_1$ ($\rho = -0.41$) increase toward the limbic belt, whereas the noradrenaline transporter increases toward koniocortex (NET, $\rho = +0.44$; all $q = 0.043$). The remaining maps show weaker, receptor-specific gradients of both signs (Supplementary Fig. S4). Grouping by pharmacological class makes the organising axis explicit: a metabotropic composite (the mean z-scored density of the metabotropic receptors) decreases with type ($\rho = -0.41$), the ionotropic composite is approximately flat, and their contrast, an ionotropic-to-metabotropic index, increases from the limbic belt to koniocortex ($\rho = +0.43$, spin $q = 0.016$; Supplementary Fig. S5). A direct excitatory-versus-inhibitory (glutamate-versus-GABA) composite points the expected way but is under-powered in this neuromodulator-dominated panel and does not survive the null. These per-receptor and composite gradients are the data-level counterpart of the molecular and cellular gradients associated with cortical type (Zaldívar-Díez and García-Cabezas, 2026, Fig. 9), recovered here from receptor densities held entirely external to the atlas. Caveats: most maps are volume-to-surface projections, the panel is dominated by neuromodulatory systems, and the composites treat the ordinal type scale as equally spaced.

**BigBrain histology (single specimen; Amunts et al., 2013).** Two attempts, both inconclusive. (i) The microstructural-profile-covariance gradient (Hist-G1; via BigBrainWarp, Paquola et al., 2021) barely tracks the type axis ($\rho \approx 0.21$, spin $p = 0.37$), so the histology-based added-value test on it was uninformative (49.9%, $\approx$ chance). We report that as a measured null: it is not evidence against the refinement, and we have not established whether low power, the single specimen, the projection or the scalar index is responsible (§3.3, §4.9). (ii) The raw intracortical intensity profiles (50 equivolumetric depths, fs_LR): a laminar complexity index was *pre-specified* (profile skewness) before analysis, with a validity gate on that index. The five profile features form their own FDR family, `bigbrain_profile_panel` (Table S4), with skewness pre-specified and

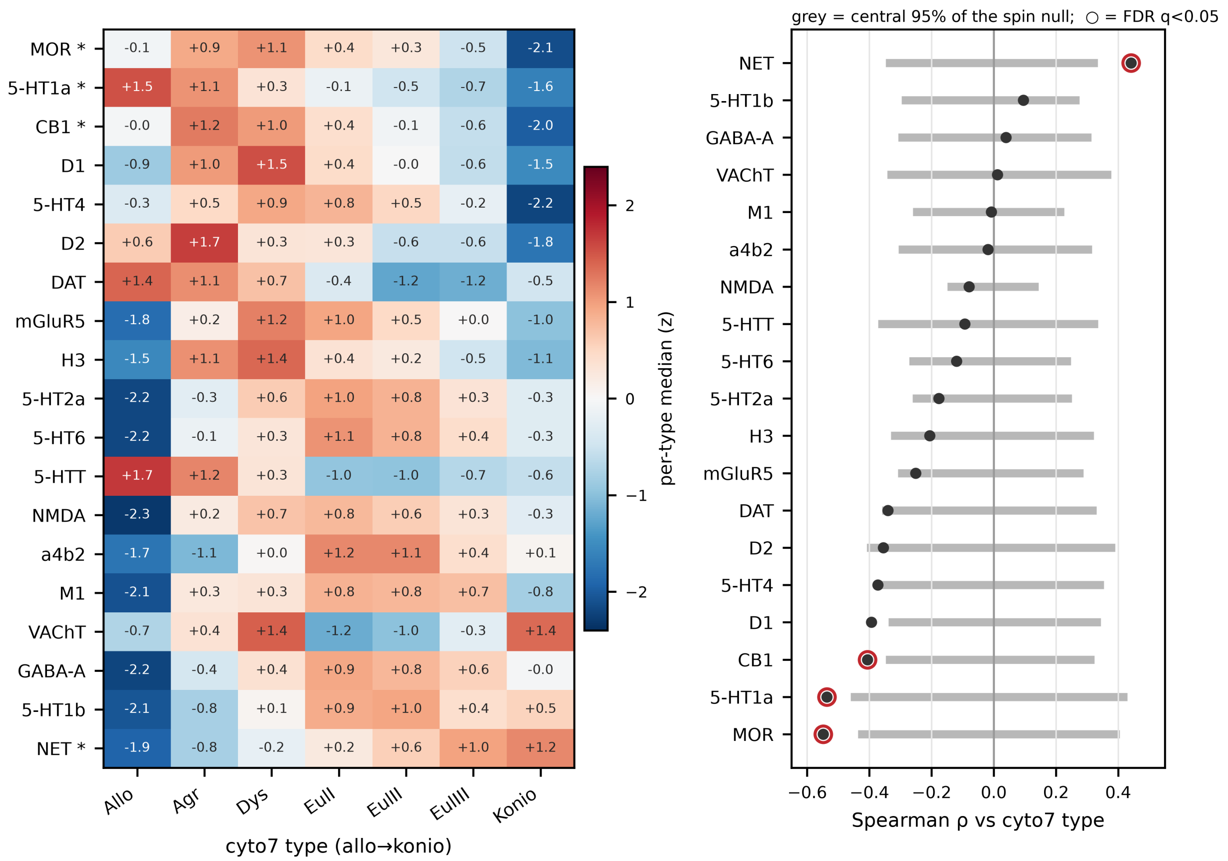


**Supplementary Figure S4. Neurotransmitter receptors and transporters along the cyto7 axis.** *Left:* z-scored per-type median density of each of the 19 Hansen PET maps across the seven cyto7 types (allocortex → koniocortex), rows ordered by their vertex-level Spearman $\rho$ with ordinal type; $*$ marks maps surviving the spin null with FDR control across the 19-map family. *Right:* forest plot of the vertex-level Spearman $\rho$ (points) with the central 95% of the spin-null distribution (grey bars), which is a null interval and not a confidence interval for the observed $\rho$; maps with FDR $q < 0.05$ are circled. Four maps survive: MOR, 5-$HT_{1A}$ and $CB_1$ decrease toward koniocortex, while the noradrenaline transporter (NET) increases.

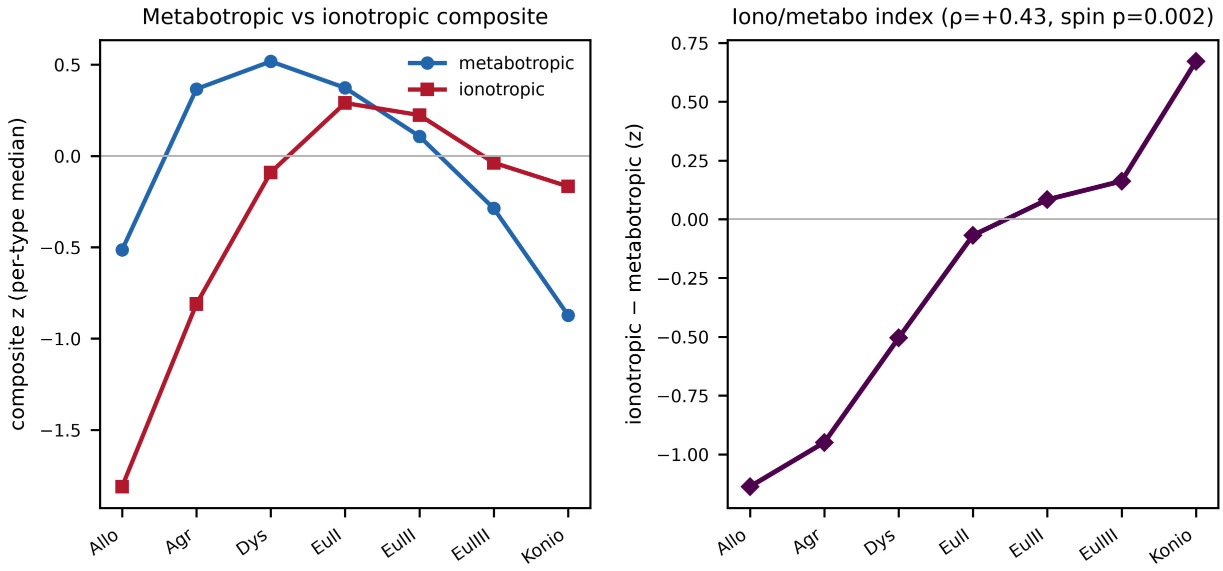


**Supplementary Figure S5. Ionotropic and metabotropic receptor composites along the cyto7 axis.** Class composites (mean z-scored density of the member maps) by cyto7 type: the metabotropic composite decreases with laminar complexity ($\rho = -0.41$) while the ionotropic composite is approximately flat, so their contrast, an ionotropic-to-metabotropic index, increases from the limbic belt to koniocortex ($\rho = +0.43$, spin $q = 0.016$). This reproduces, from receptor densities held external to the atlas, the direction of the molecular and cellular gradients associated with cortical type (Zaldívar-Díez and García-Cabezas, 2026, Fig. 9).

the other four exploratory. Per-feature $\rho$ vs type: mean $-0.28$; SD $+0.44$ (spin $q = 0.015$); skewness $-0.21$ ($q = 0.401$); kurtosis $-0.24$; layer-IV-peak prominence $+0.08$. Only SD survives the null, but it was not the pre-specified index, so it is reported as exploratory. The pre-specified skewness did not survive, so the profile-validation gate failed. The added-value test is a separate analysis and was run, using Hist-G1 as arbiter (49.9%, $\approx$ chance; §3.3). Mean intensity profiles nonetheless differ in shape by type (koniocortex shows the expected mid-cortical density peak). **Verdict:** single-specimen BigBrain profiles at this projection do not resolve the seven-type scale. Specimen variability, the surface projection, the sensitivity of the test and the reduction of a profile to a single scalar are each possible contributors, and we have not established which (§3.3, §4.9). See Supplementary Fig. S6.

**Receptor diversity.** A per-vertex receptor-diversity index was computed as the Shannon entropy of the 19 min–max-normalised Hansen maps (equivalently an effective number of receptors, $\exp H$). It decreases with cortical type in the predicted direction ($\rho = -0.38$, spin $p = 0.046$) but does not survive the Fig. 9-family FDR ($q = 0.069$); the effective number of receptors changes only modestly (16.9 at allocortex to 16.1 at koniocortex). A coefficient-of-variation variant agrees in direction.

**Evolutionary and developmental expansion.** Bilateral evolutionary cortical expansion (Xu et al., 2020, via neuromaps) decreases with type ($\rho = -0.27$, spin $p = 0.028$; allocortex-excluded $p = 0.010$). Developmental expansion (Hill et al., 2010) is available for the right hemisphere only in this release and, tested on that hemisphere, does not reach significance ($\rho = -0.11$), so it is reported as exploratory.

**Disease vulnerability.** Eight ENIGMA case–control cortical-thickness effect-size maps (Cohen's $d$, Desikan–Killiany parcels; via the ENIGMA toolbox, Larivière et al., 2021) were correlated with a continuous per-parcel cyto7 type score $c(p) = \sum_t t\, f_t(p)$, where $f_t(p)$ is the surface-area fraction of type $t$ in parcel $p$. With the ENIGMA convention that negative $d$ denotes case atrophy, a positive $\rho$ means more atrophy in lower-type cortex, that is, vulnerability decreasing with type as the Structural Model predicts. Inference used a parcel-level spin (rotation of the parcellated type score) with Benjamini–Hochberg FDR across the eight-disorder

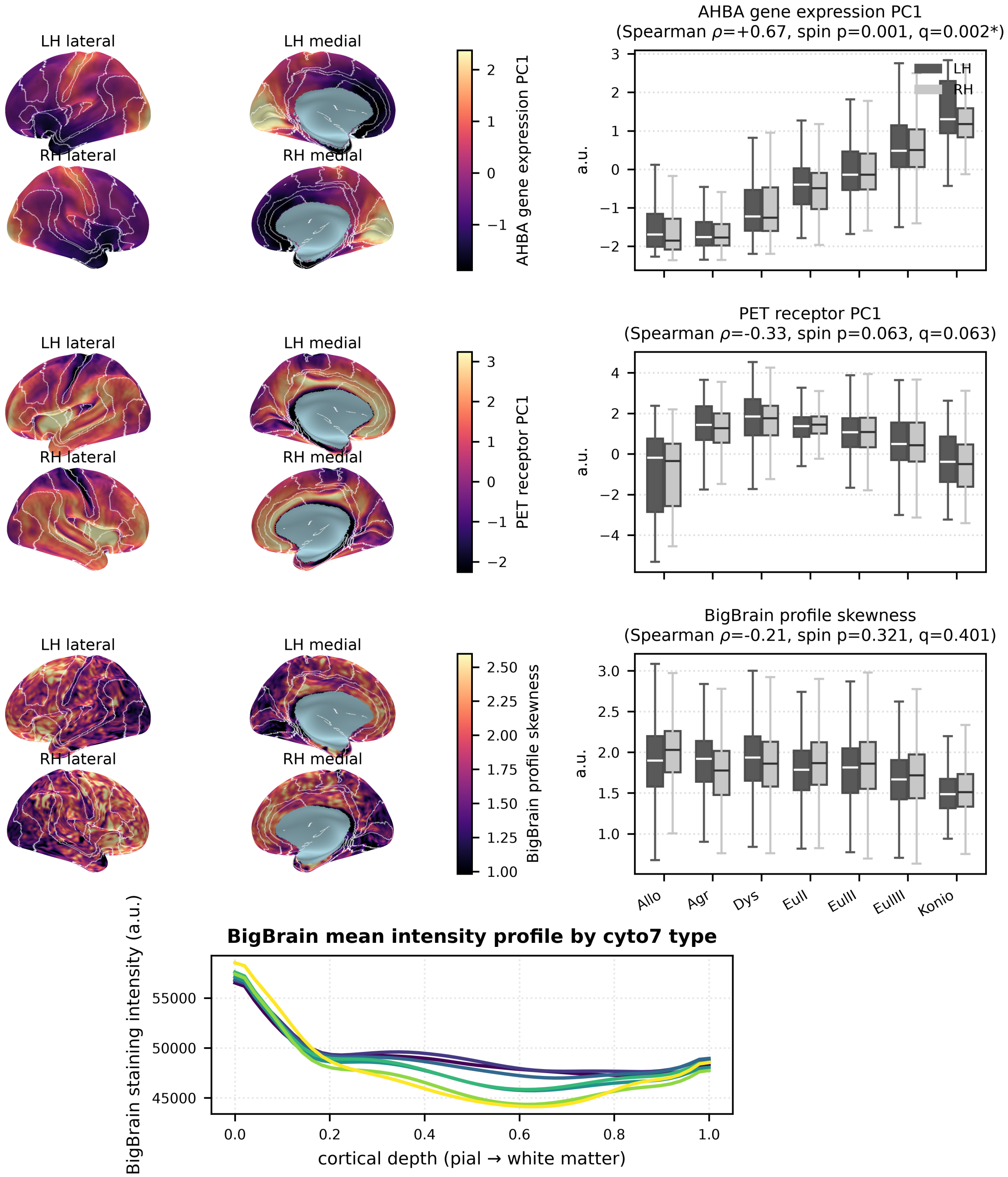


**Supplementary Figure S6. External validation: molecular and histological references.** Gene-expression PC1 (AHBA), receptor PC1 (19 PET tracers), and the BigBrain laminar complexity index (pre-specified profile skewness) on the inflated 32k fs_LR surface with cyto7 type borders (magma), each with per-type box-and-whisker distributions and the vertex-level Spearman $\rho$/spin $p$/FDR $q$; the rightmost panel shows the BigBrain mean intensity profile per cyto7 type. Gene PC1 increases across types and survives ($\rho \approx +0.67$, $q = 0.002$); receptor PC1 does not survive the spin null ($q = 0.063$); the BigBrain index is inconclusive in a single specimen (§4.9).

family; a partial Spearman correlation additionally controlled for the per-parcel principal functional gradient. Major depression ($\rho = +0.60$), ADHD (+0.41), obsessive–compulsive disorder (+0.32) and 22q11.2 deletion (+0.36) survive; depression and ADHD also survive after partialling out the functional gradient (partial $q < 0.05$), whereas obsessive–compulsive disorder does not, its signal being carried by the shared gradient. Epilepsy is significant in the opposite direction ($\rho = -0.42$), and schizophrenia, autism and bipolar disorder are null. Because these are parcel-level, cross-cohort clinical maps and a single type per parcel is lossy, the result is reported as convergent context rather than a vertex-level or predictive claim (Supplementary Fig. S7).

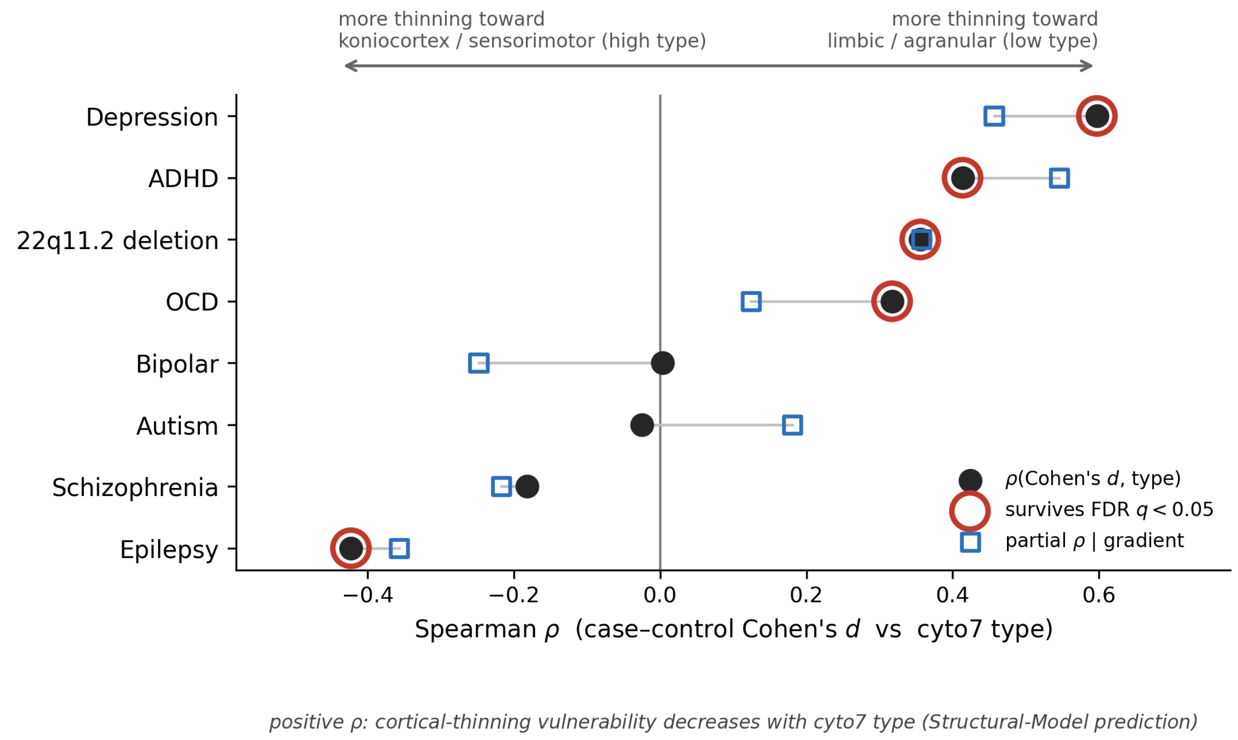


**Supplementary Figure S7. Disease vulnerability versus cyto7 type (ENIGMA).** For each of eight ENIGMA case–control cortical-thickness maps, the Spearman correlation between the parcelwise effect size (Cohen's $d$) and a continuous per-parcel cyto7 type score (Desikan–Killiany parcels); the open square is the correlation after partialling out the principal functional gradient, and ringed markers survive the parcel-level spin null with FDR control. Direction is labelled on the axis: a positive correlation places thinning toward lower-type (limbic/agranular) cortex, a negative correlation toward koniocortex/sensorimotor cortex. Depression (+0.60), ADHD, obsessive–compulsive disorder and 22q11.2 deletion show the predicted limbic-concentrated pattern (depression and ADHD surviving control for the functional gradient). Epilepsy runs opposite (−0.42): the ENIGMA all-epilepsies aggregate is weighted toward post-central, peri-Rolandic and sensorimotor thinning rather than the mesial-temporal atrophy of temporal-lobe epilepsy, which places its thinning at the koniocortex end. Schizophrenia, autism and bipolar disorder are null.

### Annex I – Cortical types across classical parcellations

The crossed parcellation was built by intersecting the released v9 cyto7 labels with each base atlas on the 164k fsaverage surface, retaining every parcel-by-type cell whose vertex count across both hemispheres exceeds 200, absorbing each sub-threshold cell whole into the largest qualifying node of the same anatomical parcel (preferring one it adjoins, so that no anatomical boundary is crossed), and keeping one node per cell irrespective of how many patches it occupies, with per-node component counts reported so that contiguous nodes can be recovered; nodes are numbered by base parcel alphabetically and then by ascending cyto7 ordinal, and because the retention threshold is evaluated bilaterally a node may fall below it in one hemisphere, so per-hemisphere vertex counts are given in the node tables.

Each Desikan–Killiany (34 parcels) and von Economo–Koskinas (43 parcels) area was cross-tabulated with cyto7 type at vertex resolution. **Result:** most classical parcels are cytoarchitecturally mixed: the median parcel is $\approx 67\%$ a single cyto7 type and spans $\approx 2$ types ($\geq 10\%$) on average. Only the primary sensory and limbic anchors are near-pure: 4/34 Desikan and 7/43 von Economo parcels are a single type at $\geq 90\%$ (e.g. V1/pericalcarine koniocortex 97%, entorhinal allocortex 98%, von Economo areas HA/HB/TD), while 12/34 Desikan and 8/43 von Economo parcels span $\geq 3$ types. Full per-parcel composition is in **Table S5**, mapped in **Supplementary Fig. S8**. The same table gives each parcel's *majority* cyto7 type, a practical per-node laminar prior for region-based whole-brain models (one neural-mass node per parcel; §4.7), with the caveat that assigning a single type per parcel discards the within-area heterogeneity ($\approx$ a third of vertices on average) that the vertex-level map preserves.

**Supplementary Table S5. Per-parcel majority cyto7 type** for the Desikan–Killiany (34 parcels) and von Economo–Koskinas (43 parcels) atlases. *Dominant fraction* is the surface-area share of the parcel occupied by its majority cyto7 type; *n types* counts the cyto7 types occupying $\geq 10\%$ of the parcel. Parcels are grouped by atlas and ordered by majority type (allocortex → koniocortex).

| Atlas | Parcel | Majority cyto7 type | Dom. frac. | $n$ types ≥10% |
|---|---|---|---|---|
| Desikan | entorhinal | Allocortex | 0.98 | 1 |
| Desikan | parahippocampal | Agranular | 0.41 | 4 |
| Desikan | caudalanteriorcingulate | Agranular | 0.89 | 2 |
| Desikan | temporalpole | Agranular | 0.58 | 3 |
| Desikan | rostralanteriorcingulate | Agranular | 0.76 | 2 |
| Desikan | posteriorcingulate | Dysgranular | 0.88 | 2 |
| Desikan | isthmuscingulate | Dysgranular | 0.68 | 3 |
| Desikan | insula | Dysgranular | 0.30 | 3 |
| Desikan | medialorbitofrontal | Eulaminate I | 0.39 | 4 |
| Desikan | lateralorbitofrontal | Eulaminate I | 0.38 | 4 |
| Desikan | inferiortemporal | Eulaminate I | 0.76 | 2 |
| Desikan | fusiform | Eulaminate I | 0.51 | 3 |
| Desikan | inferiorparietal | Eulaminate I | 0.55 | 2 |
| Desikan | middletemporal | Eulaminate I | 0.37 | 3 |
| Desikan | superiorfrontal | Eulaminate II | 0.49 | 2 |
| Desikan | caudalmiddlefrontal | Eulaminate II | 0.67 | 2 |
| Desikan | rostralmiddlefrontal | Eulaminate II | 0.67 | 2 |
| Desikan | parsopercularis | Eulaminate II | 0.79 | 2 |
| Desikan | parsorbitalis | Eulaminate II | 0.87 | 2 |
| Desikan | frontalpole | Eulaminate II | 1.00 | 1 |
| Desikan | precuneus | Eulaminate II | 0.56 | 3 |
| Desikan | superiorparietal | Eulaminate II | 0.61 | 2 |
| Desikan | superiortemporal | Eulaminate III | 0.68 | 2 |
| Desikan | paracentral | Eulaminate III | 0.59 | 4 |
| Desikan | parstriangularis | Eulaminate III | 0.59 | 2 |
| Desikan | bankssts | Eulaminate III | 0.68 | 2 |
| Desikan | supramarginal | Eulaminate III | 0.72 | 2 |
| Desikan | precentral | Eulaminate III | 0.45 | 3 |
| Desikan | lateraloccipital | Eulaminate III | 0.68 | 2 |
| Desikan | lingual | Eulaminate III | 0.72 | 3 |
| Desikan | cuneus | Eulaminate III | 0.91 | 1 |
| Desikan | postcentral | Eulaminate III | 0.59 | 2 |
| Desikan | transversetemporal | Koniocortex | 0.81 | 2 |
| Desikan | pericalcarine | Koniocortex | 0.97 | 1 |
| von Economo | HA | Allocortex | 1.00 | 1 |
| von Economo | HB | Allocortex | 1.00 | 1 |
| von Economo | HC | Allocortex | 0.42 | 2 |
| von Economo | LA2 | Agranular | 0.67 | 2 |
| von Economo | FLMN | Agranular | 0.56 | 2 |
| von Economo | TG | Dysgranular | 0.50 | 3 |
| von Economo | LD | Dysgranular | 0.65 | 2 |
| von Economo | LE | Dysgranular | 0.75 | 2 |
| von Economo | LA1 | Dysgranular | 0.68 | 2 |

*Table S5 (continued)*

| Atlas | Parcel | Majority cyto7 type | Dom. frac. | $n$ types ≥10% |
|---|---|---|---|---|
| von Economo | FJK | Dysgranular | 0.58 | 2 |
| von Economo | LC3 | Dysgranular | 1.00 | 1 |
| von Economo | FH | Dysgranular | 0.81 | 2 |
| von Economo | LC2 | Dysgranular | 0.91 | 1 |
| von Economo | IA | Dysgranular | 0.63 | 2 |
| von Economo | TF | Eulaminate I | 0.63 | 2 |
| von Economo | LC1 | Eulaminate I | 0.52 | 2 |
| von Economo | FF | Eulaminate I | 0.34 | 4 |
| von Economo | FG | Eulaminate I | 0.87 | 2 |
| von Economo | FC | Eulaminate I | 0.92 | 1 |
| von Economo | PG | Eulaminate I | 0.70 | 2 |
| von Economo | FE | Eulaminate II | 0.67 | 2 |
| von Economo | FD | Eulaminate II | 0.79 | 2 |
| von Economo | FCBm | Eulaminate II | 0.77 | 2 |
| von Economo | FB | Eulaminate II | 0.73 | 2 |
| von Economo | PH | Eulaminate II | 0.63 | 3 |
| von Economo | FDdelta | Eulaminate II | 1.00 | 1 |
| von Economo | PE | Eulaminate II | 0.80 | 2 |
| von Economo | IB | Eulaminate III | 0.35 | 4 |
| von Economo | TE | Eulaminate III | 0.43 | 3 |
| von Economo | TB | Eulaminate III | 0.46 | 3 |
| von Economo | TA | Eulaminate III | 0.64 | 1 |
| von Economo | FDT | Eulaminate III | 0.63 | 3 |
| von Economo | PD | Eulaminate III | 0.58 | 2 |
| von Economo | PF | Eulaminate III | 0.68 | 3 |
| von Economo | FA | Eulaminate III | 0.67 | 2 |
| von Economo | OA | Eulaminate III | 0.83 | 2 |
| von Economo | OB | Eulaminate III | 0.79 | 2 |
| von Economo | PC | Eulaminate III | 0.76 | 2 |
| von Economo | OC | Koniocortex | 0.67 | 2 |
| von Economo | PA | Koniocortex | 0.62 | 2 |
| von Economo | PB | Koniocortex | 0.67 | 2 |
| von Economo | TC | Koniocortex | 0.62 | 2 |
| von Economo | TD | Koniocortex | 1.00 | 1 |

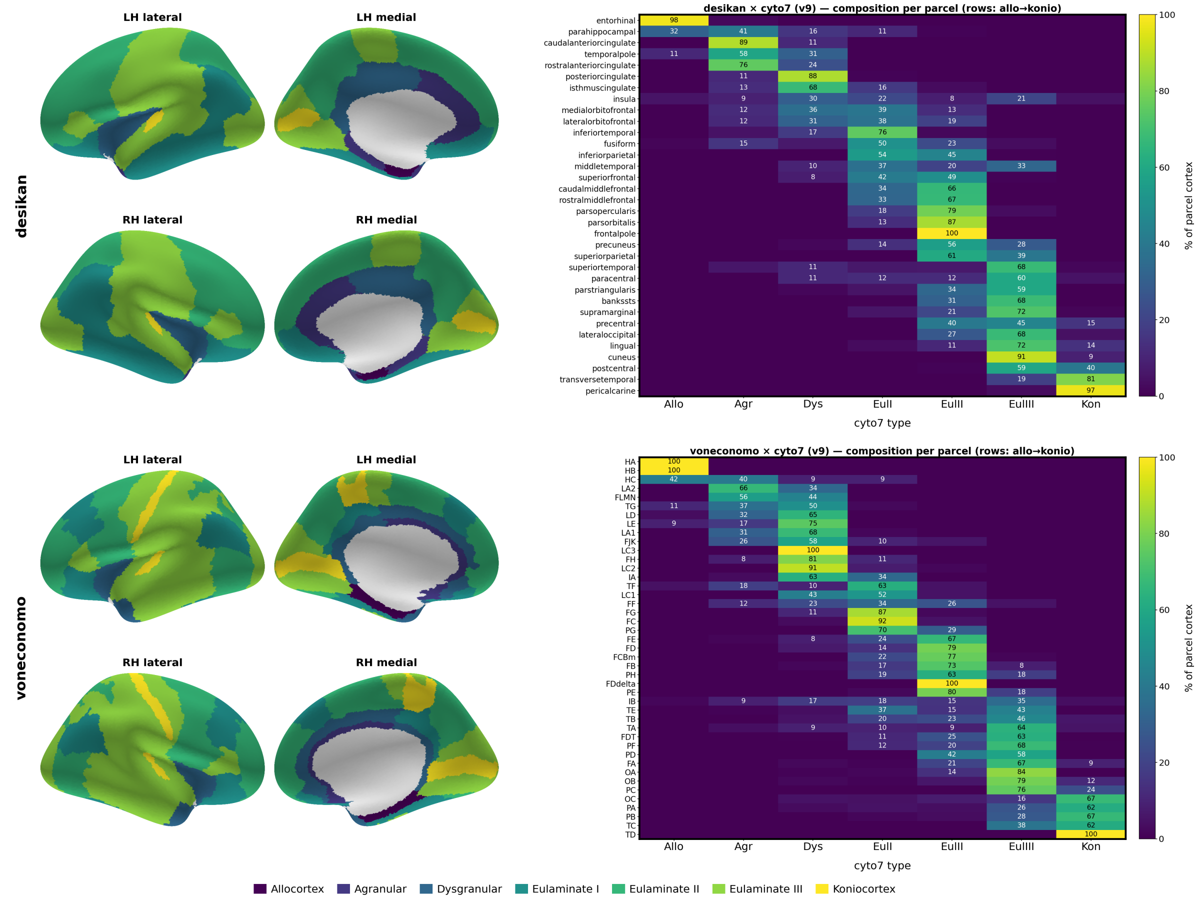


**Supplementary Figure S8. Cortical types across classical parcellations.** *Left blocks:* the Desikan–Killiany (top) and von Economo–Koskinas (bottom) parcellations displayed on the fsaverage inflated surface (LH and RH, lateral and medial views), with each parcel coloured by its majority cyto7 type (viridis palette, allocortex → koniocortex; medial wall in grey). *Right:* the corresponding parcel × cyto7-type composition matrices (both hemispheres combined, post-cleanup); rows are parcels sorted allocortex → koniocortex, columns are the seven cyto7 types, and each cell gives the percentage of that parcel's labelled cortex assigned to the type (values $\geq 8\%$ annotated). Most parcels are cytoarchitecturally mixed rather than one-to-one with a cortical type: the median dominant-type fraction is 0.67 for both atlases, with on average 2.4 (Desikan) and 2.0 (von Economo) types per parcel at $\geq 10\%$ occupancy, and only 4/34 Desikan and 7/43 von Economo parcels reaching $\geq 90\%$ of a single type. The near-pure parcels are the primary sensory koniocortical anchors (e.g. pericalcarine/V1 97%, transverse temporal/A1 81%) and the limbic allocortical anchors (e.g. entorhinal 98%). The per-parcel majority-type lookup is listed in Table S5.

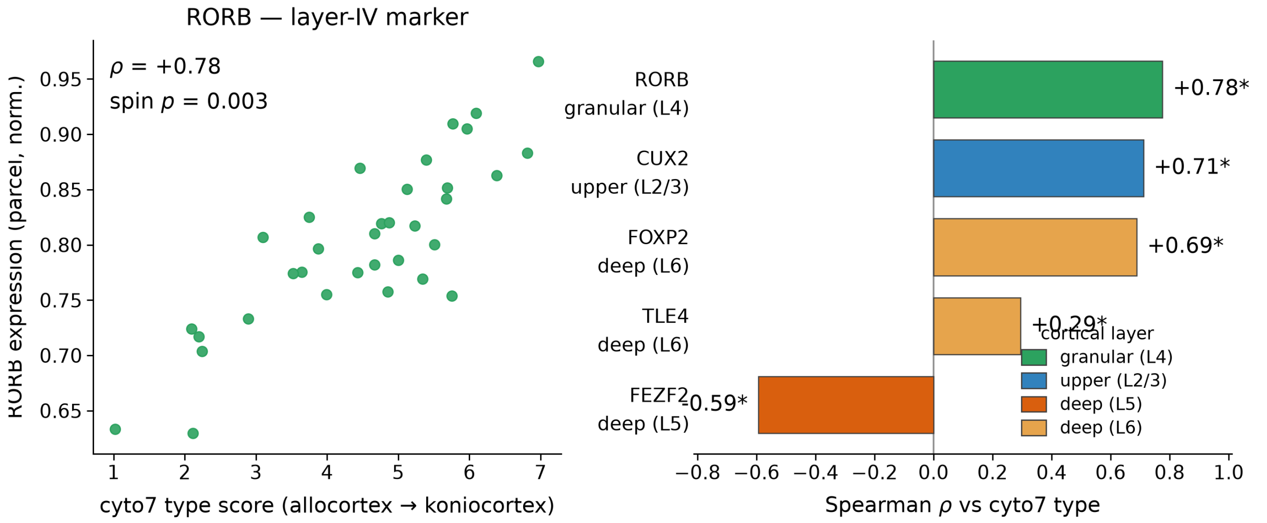


**Supplementary Figure S9. Laminar marker genes track cytoarchitectural type.** Left: parcel-level expression of the layer-IV marker *RORB* (AHBA via `abagen`; 68 Desikan–Killiany parcels, hemisphere-averaged for display) against the continuous cyto7 type score, increasing from allocortex to koniocortex ($\rho = +0.78$, parcel spin $p = 0.003$). Right: Spearman correlation with ordinal type for each layer-marker gene, coloured by the cortical layer it marks; * marks survival of the parcel spin null with FDR control within the layer-marker family. The family has eight members: the five single genes, which were pre-specified with their directions predicted a priori from the layer each marker labels, and three z-composites, which were not pre-specified and are exploratory. RORB is the one confirmatory test; the other four genes and the three composites are exploratory (Table S4, Table S8), on the reasoning that markers of layers that elaborate with laminar complexity should increase and markers of layer V should not: RORB $\rho = +0.78$ ($q = 0.005$, the one specified-in-advance primary), CUX2 +0.71 ($q = 0.005$), FOXP2 +0.69 ($q = 0.005$), TLE4 +0.29 ($q = 0.049$) and FEZF2 −0.59 ($q = 0.005$); the composites are upper +0.71, granular +0.78 and deep +0.33. All eight survive. The upper (CUX2), layer-IV (RORB) and deep L6 (FOXP2, TLE4) markers increase with laminar complexity while the layer-V marker FEZF2 decreases ($\rho = -0.59$), the laminar specificity expected if the axis reads genuine layer identity. RORB was pre-specified as a molecular test of the layer-IV definitional criterion; layer assignments are literature-derived, not from these data. AHBA caveats: 5 of 6 donors, transcript not protein, mostly left hemisphere, coarse Desikan parcels.

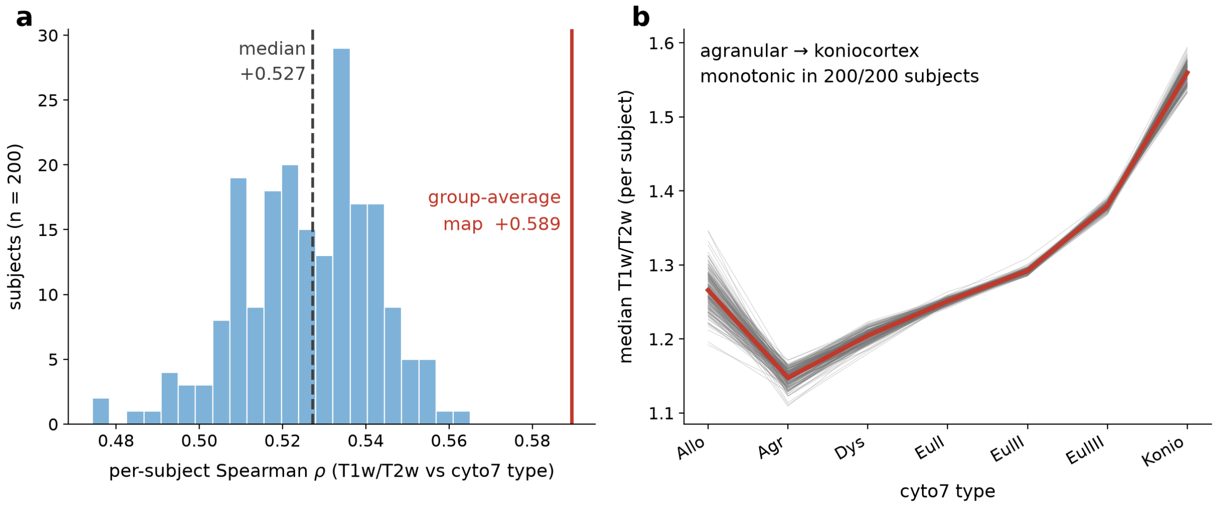


**Supplementary Figure S10. The cyto7 type axis generalises to individual subjects.** The group map projected onto 200 individual HCP subjects (individual bias-corrected T1w/T2w on fs_LR 32k). (a) Distribution of the per-subject Spearman correlation between individual T1w/T2w and ordinal cyto7 type (median +0.53, dashed); every subject is positive and all fall below the group-average map value (+0.59, red), which is therefore an upper bound on the individual fit, not a typical one. (b) Each subject's median T1w/T2w per cyto7 type (grey; across-subject median in red): the increase from agranular to koniocortex is strictly monotonic in all 200 subjects, with allocortex off the ramp (elevated, hence excluded from the trend fits; §2.6).

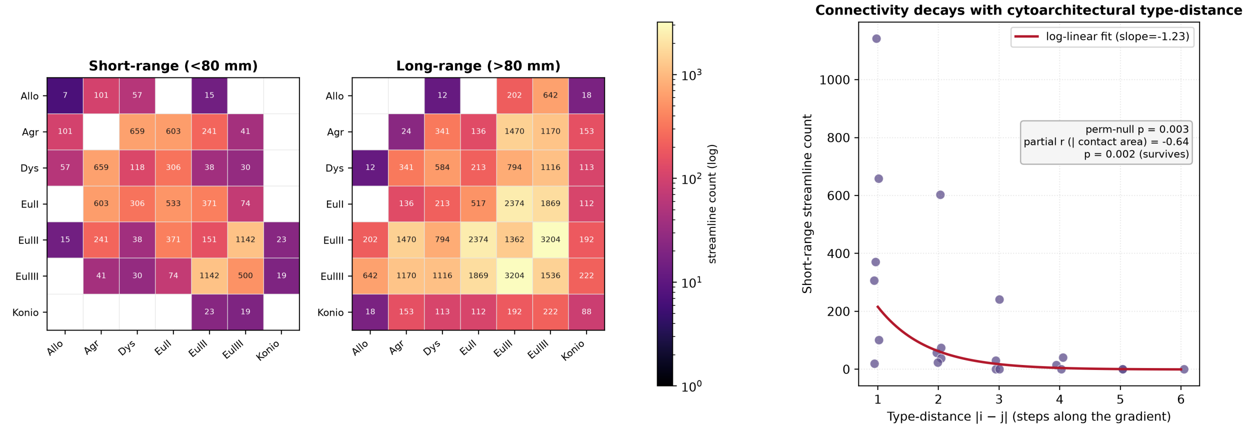


**Supplementary Figure S11. Short-range cortico-cortical connectivity by cytoarchitectural type-distance (exploratory).** *a,b* Type-by-type streamline counts from the population-averaged HCP-1065 template tractogram, split into short-range ($<$ 80 mm, *a*) and long-range ($>$ 80 mm, *b*) bundles; rows and columns are the seven cyto7 types and cells are streamline counts on a **shared logarithmic colour scale** (magma). Short-range connections concentrate near the diagonal, between cytoarchitecturally similar types, whereas long-range connections are broader and denser. *c* Short-range streamline count versus cytoarchitectural type-distance $|i - j|$ (steps along the gradient of laminar complexity), with a log-linear fit (slope $\approx -1.23$). The falloff survives a label-permutation null ($p = 0.003$) and a contact-area partial correlation ($r = -0.64$, $p = 0.002$). The raw-count statistic then only marginally clears a spin null, which slides the whole map on the sphere and so leaves its spatial arrangement intact ($p = 0.040$) against a null already centred at $r = -0.334$, so roughly a third of the observed effect follows from the geometry of the map alone; the opportunity-corrected statistic does not clear it ($p = 0.070$), and at the level of the template bundles the association is absent ($\beta = +0.01$, $p = 0.96$). It is therefore reported as exploratory. The result is stated as a relative, ordinal prediction given the group-averaged template tractogram and surface-to-volume projection (§4).

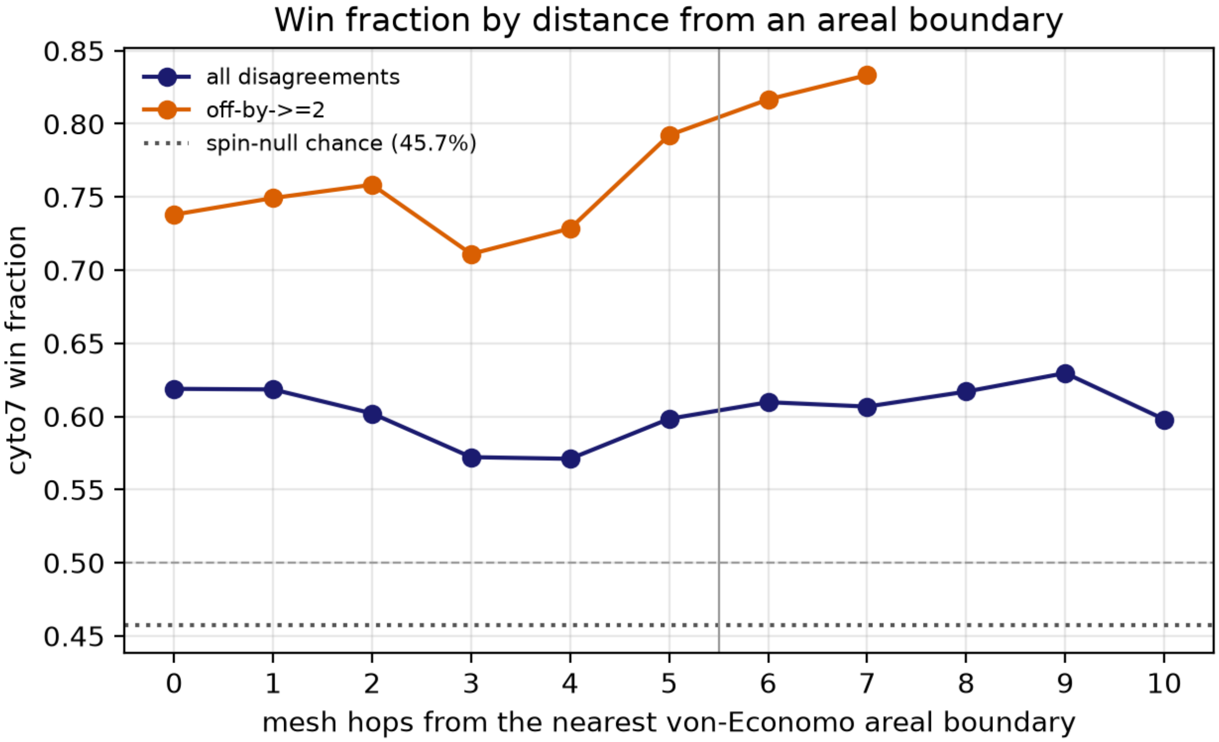


**Supplementary Figure S12. The vertex-level advantage over the area-level map holds inside areas, not only at their borders.** Win fraction (the proportion of disagreement vertices at which the cyto7 label sits closer to its own type's median on the independent multimodal profile than the area-level label does) as a function of graph-hop distance from the nearest von Economo areal border, in 1-hop bins out to ten hops and pooled beyond. The profile is flat: 61.9% at the border and 59.8% at ten hops, against a spin-null chance level of roughly 45%. Aggregated, the areal interior (more than five hops, $n$ = 4,844) gives 58.7% (chance 44.8%, spin $p$ = 0.007) against 60.4% in the boundary band ($n$ = 22,454). Had the advantage been confined to areal borders, the profile would decay toward chance with distance, which is the pattern expected if the vertex map were only smoothing areal edges (§3.3).

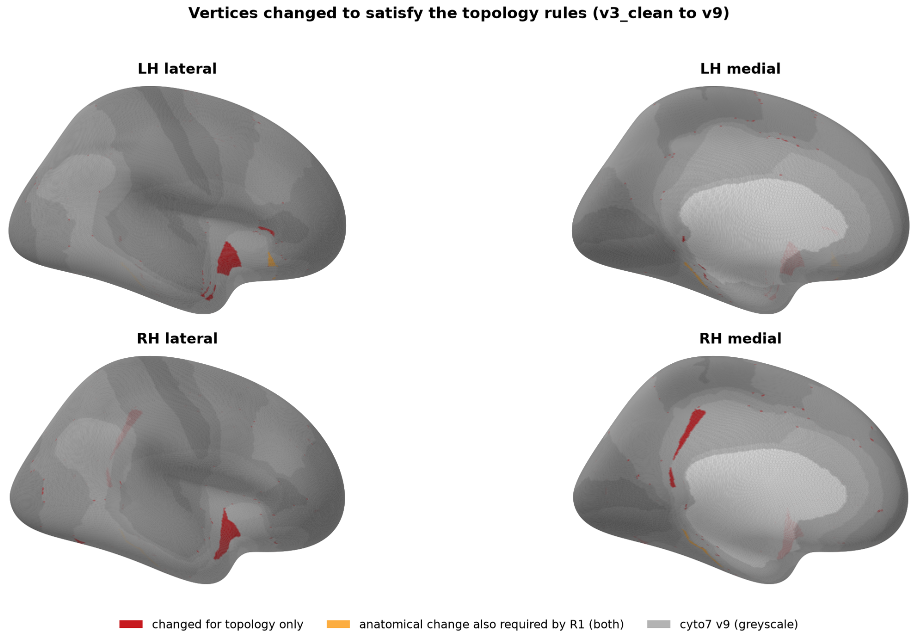


**Supplementary Figure S13. Where the topology repairs are.** Vertices relabelled solely to satisfy the sequential-gradient rule R1 (2,368 vertices, 0.78% of labelled cortex, 960 mm$^2$), shown on the inflated fsaverage surface over a greyed cyto7 map, both hemispheres in lateral and medial views. Eighty-four per cent are dysgranular and they concentrate in the peri-insular and superior-temporal belt; 28 Desikan parcels are touched. The rule strictly required 42 vertices, so most of this extent is a drawn gradient rather than a minimal repair (Annex A). Removing all of them with a two-hop margin changes no primary correlation by more than $\Delta\rho = 0.011$.

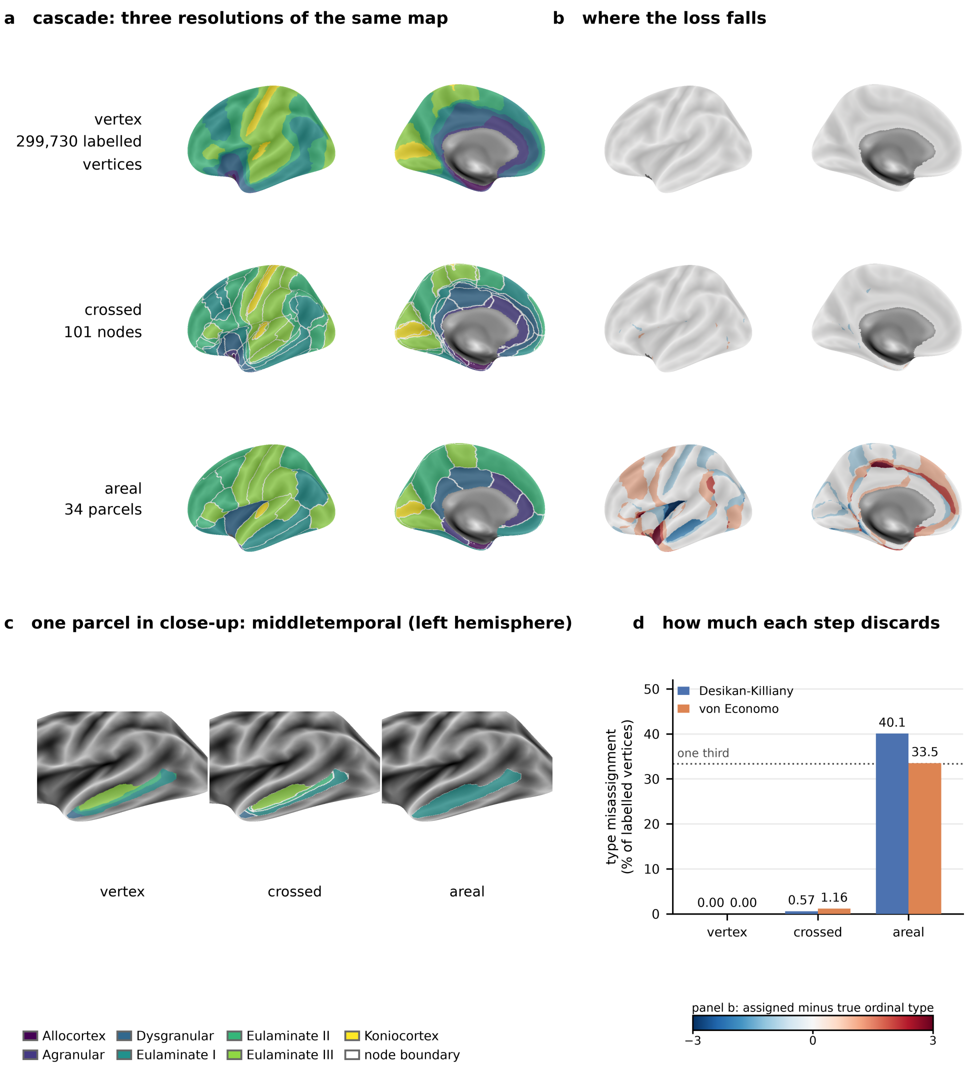


**Supplementary Figure S14. The same cytoarchitectural map at three resolutions, and what each coarsening step discards.** *(a)* The released cyto7 map on the inflated fsaverage surface (left hemisphere, lateral and medial) at three resolutions: the vertex map (304,972 labelled vertices across both hemispheres), the crossed parcellation (101 Desikan-Killiany nodes, §4.7), and one majority cyto7 type per Desikan-Killiany parcel (34 parcels). Node and parcel boundaries are drawn in thin white. The three surfaces look nearly alike, which is the point of panel b. *(b)* The same three resolutions shown as the type misassignment map: the signed ordinal difference between the type a vertex is assigned and its true cyto7 type, on the diverging scale of Figure 2b (differences beyond three ordinal steps, which reach five and are rare, are clipped to the ends of the scale). The vertex row is blank by construction, the crossed row is almost blank, and the areal row is not. *(c)* One parcel in close-up: middletemporal in the left hemisphere, the Desikan-Killiany parcel spanning the most cyto7 types at above 10% of its area (eulaminate I 36.4%, eulaminate III 33.7%, eulaminate II 17.4%, dysgranular 12.5%) and the largest such parcel at 4,104 mm$^2$. Its eulaminate III band survives the crossing and disappears when the parcel takes a single majority type. *(d)* Type misassignment as a percentage of labelled vertices at each resolution, for both base atlases: zero at vertex level by construction; 0.57% (Desikan-Killiany) and 1.16% (von Economo) for the crossed parcellation, which is the residue absorbed by the 200-vertex floor; and 40.1% and 33.5% for one type per parcel. The dotted line marks one third. Per-node and per-resolution values are in `misassignment_fractions.csv`.

### Annex J – Comparison against quantised continuous maps

**What this tests.** Every validation correlation reported here could in principle be produced by any map that orders cortex along the unimodal-transmodal axis, in which case the anatomical labour would buy interpretability but no predictive content. We tested that directly by building three seven-level equal-count septiles on the same 58,731 vertices and the same medial-wall mask as the released analyses: of T1w/T2w myelin, of the principal functional-connectivity gradient, and of the BigBrain microstructural-profile-covariance gradient (Hist-G1). Equal-count rather than equal-width quantisation is used so that a comparator matches cyto7's ordinal structure without importing its type-size distribution. Each was then correlated with the same feature panel as cyto7, under the same spin nulls and with Benjamini–Hochberg control within each partition, and reduced to Desikan–Killiany parcels by the same area-weighted composition rule used for cyto7 in the parcel-level tests. The comparators are analysis objects and are not part of the release. The panel covers the twelve vertex-level features and the five laminar marker genes, which are the measures available on a common vertex or parcel grid for every partition. It does not cover the per-subject MEG dynamics, the receptor composites or the disease maps: the first are defined per subject rather than on the group grid, and the last two are parcel-level families whose members are individually exploratory. We did not run the MEG comparison and make no claim about how it would come out. The same fixed septile map could be applied to every subject exactly as `cyto7` is, and at the group level the timescale comparison already favours the myelin septile, so there is no basis for assuming the per-subject result would go the other way.

**Vertex level: the comparators attain the higher correlation.** On the twelve vertex-level features a septile attains the higher $|\rho|$ in every case, and cyto7 in none. Part of this is by construction and part is not, and the two should be separated. The three diagonal entries below (myelin under the myelin septile, the functional gradient under the gradient septile, Hist-G1 under the Hist-G1 septile) are arithmetic: a septile reproduces its own source map at $|\rho| = 0.99$. The other nine are empirical results, not guaranteed by construction, and they are consistent with T1w/T2w myelin carrying most of the macroscale axis, so that a feature correlated with myelin tends to favour a myelin septile. We report them as an outcome rather than as an inevitability.

**Supplementary Table S6.** Comparator panel, vertex level. Absolute Spearman correlation with ordinal level for cyto7 and for the three septiles, over the twelve vertex-level features. The strongest value in each row is bold. Signs are omitted because the gradient septile inverts the axis; signed values, spin $p$ and within-partition $q$ are in `comparator_panel.csv`.

| Feature | cyto7 | myelin | gradient | Hist-G1 |
|---|---|---|---|---|
| T1w/T2w myelin | 0.59 | **0.99** | 0.62 | 0.34 |
| Cortical thickness | 0.50 | **0.56** | 0.39 | 0.40 |
| Intrinsic timescale | 0.48 | **0.61** | 0.53 | 0.38 |
| Delta power (0.5-4 Hz) | 0.44 | **0.61** | 0.48 | 0.46 |
| Theta power (4-8 Hz) | 0.18 | 0.39 | 0.18 | **0.58** |
| Alpha power (8-12 Hz) | 0.37 | 0.57 | 0.36 | **0.60** |
| Beta power (12-30 Hz) | 0.27 | 0.16 | **0.29** | 0.21 |
| Gamma power (30-60 Hz) | 0.31 | 0.52 | 0.28 | **0.53** |
| Principal functional gradient | 0.58 | 0.63 | **0.99** | 0.23 |
| AHBA gene expression PC1 | 0.67 | **0.84** | 0.59 | 0.44 |
| BigBrain Hist-G1 | 0.21 | 0.35 | 0.23 | **0.99** |
| PET receptor PC1 | 0.33 | **0.60** | 0.47 | 0.25 |

**Laminar markers: cyto7 attains the higher correlation on four of five.** The difference appears at the measures that bear on the defining criterion, layer-IV granularity, rather than on the generic hierarchy. Across the five laminar marker genes cyto7 attains the higher $|\rho|$ for four, and is weaker only for TLE4. The FEZF2 margin is the widest, 0.60 against 0.17 for the myelin septile. Direction does not explain it: the comparison is of absolute correlations, so a myelin septile can represent a feature running against myelin with the opposite sign, and the margin is an empirical result rather than a consequence of FEZF2 marking layer V. Against the Hist-G1 septile, the one comparator built from histology rather than from an MRI proxy, cyto7 is higher on all five. None of these comparisons tests whether the differences between maps are statistically reliable; that needs a paired null preserving the dependence among the competing maps, which we have not run.

**Supplementary Table S7.** Comparator panel, laminar marker genes. Absolute Spearman correlation with ordinal level across the 68 Desikan–Killiany parcels, for cyto7 and the three septiles. The cyto7 level is the hemisphere-pooled type score, evaluated at each of the 68 hemisphere-specific parcels; the "34" of earlier drafts referred to the 34 named parcel identities, not to the number of observations. The strongest value in each row is bold. cyto7's values here are recomputed on the composed parcellation and agree with the released fsaverage values to within 0.012 (`comparator_panel_parcel.csv`).

| Gene | cyto7 | myelin | gradient | Hist-G1 |
|---|---|---|---|---|
| *RORB* | **0.77** | 0.63 | 0.34 | 0.68 |
| *CUX2* | **0.72** | 0.56 | 0.29 | 0.68 |
| *FEZF2* | **0.60** | 0.17 | 0.04 | 0.26 |
| *FOXP2* | **0.69** | 0.49 | 0.27 | 0.62 |
| *TLE4* | 0.29 | **0.50** | 0.32 | 0.12 |

**The win-fraction benchmark cannot separate them.** Substituted for cyto7 as the challenger against the area-level map on the same 27,298 disagreement vertices, the myelin septile scores 0.703 and the gradient septile 0.714, both above cyto7's 0.601. Each, however, is scored partly against the arbiter it was built from, at 0.909 and 0.926 respectively. Because each arbiter contributes the same vertex count, the pooled win fraction is exactly the mean of the three per-arbiter fractions, so dropping each partition's own defining arbiter is an exact operation and not an approximation. The same arbiter must be dropped from cyto7's side as well, or the two are not scored on the same features: without myelin, cyto7 is 0.594 against the myelin septile's 0.599; without the functional gradient, 0.592 against the gradient septile's 0.608. The gaps are 0.005 and 0.016 in the septiles' favour. We have not estimated their uncertainty or stated a practical-equivalence criterion, so we read this as the benchmark failing to separate the maps rather than as either winning. The Hist-G1 septile, built from none of the three arbiters, reaches 0.502 and is below chance on two of them. The benchmark therefore measures alignment with the macroscale hierarchy, which any map encoding that axis attains, and it is not a test that discriminates among such maps.

**The granularity-matched null.** The published chance level of 46% comes from rotating cyto7 and rebuilding its per-type medians from the rotated labels, which computes those medians over anatomically arbitrary tissue and so collapses them toward the global median. A plausible seven-class alternative is a harder comparison. We built 200 spatially constrained seven-class partitions by region-growing from random seeds with target sizes taken from cyto7 itself, grown within each hemisphere because the fs_LR mesh is disconnected at the midline; each partition comes out at exactly two connected components per class, one per hemisphere, with a worst-case class-size error of 11.2%. This matches `cyto7` for types 1 to 5 but not for eulaminate III and koniocortex, which form three islands per hemisphere (Table S2), so the null's classes are more compact than the two most fragmented `cyto7` types. What the null matches is class-size distribution, class count and contiguity; it does not match component counts, ordinal adjacency or boundary complexity, and the 11.2% worst-case size error is the limit of "matched" here. A size-rebalancing pass was tried and discarded because it restored sizes at the cost of contiguity, and a ragged partition would make the baseline artificially easy to beat. Their win fractions average 0.542 (s.d. 0.030, range 0.455 to 0.621), so the rotation baseline of 0.46 sits at the 0.5th percentile of this distribution and was too generous. cyto7's observed 0.601 sits at the 98.5th percentile, $p = 0.020$. The advantage survives a granularity-matched alternative, with the margin reduced from fourteen points to six. This test and the three comparator win fractions are tabulated in Table S8 as the `benchmark_controls` family.

**Reproduction of the published statistics.** Two inputs had to be reconstructed exactly before the comparators could be scored against them, and both were checked against released values. The disagreement set recomputes to 27,298 vertices, matching §3.3, only when the area-level reference is the decoded type-per-area map rather than the raw areal labels. cyto7's win fraction recomputes to 0.6010, matching the published 60.1%, only when the three arbiter judgements are pooled across vertices rather than voted per vertex; a majority vote over the three arbiters would give 0.628. The rotation permutation was recovered once at seed

0 and validated by reproducing the released null file before being reused for every partition, so no released null was regenerated.

**Supplementary Table S8.** Outcome table: the declared FDR families. Every test in the declared FDR families, with its hypothesis status, statistical unit, null, FDR family, raw $p$, within-family $q$ and effect size. Status records whether a test was specified in advance, not whether its result is retained: the connectivity analysis of Annex G was specified in advance but is reported as exploratory there, because it does not survive a spin null or a bundle-level model. An asterisk marks a test that survives the null in its Null column: at $q < 0.05$ within its family where Benjamini–Hochberg correction was applied, and at $p < 0.05$ where it was not. Families are never pooled. Tests reported without a $q$ had no FDR applied within their family and are marked accordingly; Table S4 lists every family with its size, whether correction was applied and why. Rows whose status is *descriptive* carry no null and no test and are never asterisked: the per-feature win fractions on the disagreement set are reported this way, because the vertices are not independent and the chance baseline for a win fraction is not 0.5 but the level of the relevant empirical null, which differs between comparisons. The aggregate win fractions are tested against their own null distributions and are asterisked. A few robustness analyses belong to a declared family and are listed here with the status *robustness*; those that belong to no declared family are disclosed separately in Table S9, whose caption states the correction policy applied to them and the one exception to it.

| Measure | Family ($n$) | Status | Unit ($n_{\text{obs}}$) | Null | $p_{\text{raw}}$ | $q$ | |
|---|---|---|---|---|---|---|---|
| T1w/T2w myelin<br>Spearman $\rho = +0.5894$ | structure_function_9 (9) | confirmatory | vertex (32k fs_LR), n=58731 | vertex spin (N=1000, seed 0) | 0.001 | 0.004 | * |
| Cortical thickness<br>Spearman $\rho = -0.5018$ | structure_function_9 (9) | confirmatory | vertex (32k fs_LR), n=58731 | vertex spin (N=1000, seed 0) | 0.002 | 0.006 | * |
| Intrinsic timescale<br>Spearman $\rho = -0.4848$ | structure_function_9 (9) | confirmatory | vertex (32k fs_LR), n=58731 | vertex spin (N=1000, seed 0) | 0.046 | 0.103 | |
| Delta power (0.5-4 Hz)<br>Spearman $\rho = -0.4427$ | structure_function_9 (9) | confirmatory | vertex (32k fs_LR), n=58731 | vertex spin (N=1000, seed 0) | 0.110 | 0.198 | |
| Theta power (4-8 Hz)<br>Spearman $\rho = -0.1777$ | structure_function_9 (9) | confirmatory | vertex (32k fs_LR), n=58731 | vertex spin (N=1000, seed 0) | 0.623 | 0.623 | |
| Alpha power (8-12 Hz)<br>Spearman $\rho = +0.3692$ | structure_function_9 (9) | confirmatory | vertex (32k fs_LR), n=58731 | vertex spin (N=1000, seed 0) | 0.257 | 0.371 | |
| Beta power (12-30 Hz)<br>Spearman $\rho = +0.2660$ | structure_function_9 (9) | confirmatory | vertex (32k fs_LR), n=58731 | vertex spin (N=1000, seed 0) | 0.416 | 0.468 | |
| Gamma power (30-60 Hz)<br>Spearman $\rho = -0.3082$ | structure_function_9 (9) | confirmatory | vertex (32k fs_LR), n=58731 | vertex spin (N=1000, seed 0) | 0.289 | 0.371 | |
| Principal functional gradient<br>Spearman $\rho = -0.5761$ | structure_function_9 (9) | confirmatory | vertex (32k fs_LR), n=58731 | vertex spin (N=1000, seed 0) | 0.001 | 0.004 | * |
| AHBA gene expression PC1<br>Spearman $\rho = +0.6657$ | external_reference_2 (2) | confirmatory | vertex (32k fs_LR), n=57957 | vertex spin (N=1000, seed 0) | 0.001 | 0.002 | * |
| PET receptor PC1 (Hansen-2022 collection)<br>Spearman $\rho = -0.3292$ | external_reference_2 (2) | confirmatory | vertex (32k fs_LR), n=58731 | vertex spin (N=1000, seed 0) | 0.063 | 0.063 | |
| BigBrain Profile mean intensity<br>Spearman $\rho = -0.2826$ | bigbrain_profile_ panel_5 (5) | exploratory | vertex (32k fs_LR), n=58731 | vertex spin (N=1000, seed 0) | 0.209 | 0.348 | |
| BigBrain Profile SD<br>Spearman $\rho = +0.4357$ | bigbrain_profile_ panel_5 (5) | exploratory | vertex (32k fs_LR), n=58731 | vertex spin (N=1000, seed 0) | 0.003 | 0.015 | * |
| BigBrain Profile skewness (pre-specified)<br>Spearman $\rho = -0.2091$ | bigbrain_profile_ panel_5 (5) | confirmatory | vertex (32k fs_LR), n=58731 | vertex spin (N=1000, seed 0) | 0.321 | 0.401 | |
| BigBrain Profile kurtosis<br>Spearman $\rho = -0.2422$ | bigbrain_profile_ panel_5 (5) | exploratory | vertex (32k fs_LR), n=58731 | vertex spin (N=1000, seed 0) | 0.133 | 0.332 | |

| Measure | Family ($n$) | Status | Unit ($n_{\mathrm{obs}}$) | Null | $p_{\mathrm{raw}}$ | $q$ | |
|---|---|---|---|---|---|---|---|
| BigBrain Layer-IV peak prominence<br>Spearman $\rho = +0.0848$ | bigbrain__profile__panel_5 (5) | exploratory | vertex (32k fs_LR), n=58731 | vertex spin (N=1000, seed 0) | 0.674 | 0.674 | |
| Aperiodic exponent (fixed, 2-40 Hz, pre-specified)<br>mean over 89 subjects of the per-subject vertex-level Spearman $\rho$ = -0.1192 | dynamics__singlemember_7 (7) | confirmatory | subject (N=89) x vertex (4k fs_LR), n=89 subjects x 8004 vertices | hierarchical subject x vertex spin (N=1000, seed 0) | 0.499 | 0.582 | |
| Peak frequency<br>mean over 89 subjects of the per-subject vertex-level Spearman $\rho$ = +0.0280 | dynamics__singlemember_7 (7) | confirmatory | subject (N=89) x vertex (4k fs_LR), n=89 subjects x 8004 vertices | hierarchical subject x vertex spin (N=1000, seed 0) | 0.824 | 0.824 | |
| Spectral offset<br>mean over 89 subjects of the per-subject vertex-level Spearman $\rho$ = -0.1618 | dynamics__singlemember_7 (7) | confirmatory | subject (N=89) x vertex (4k fs_LR), n=89 subjects x 8004 vertices | hierarchical subject x vertex spin (N=1000, seed 0) | 0.213 | 0.298 | |
| Intrinsic timescale (knee-derived)<br>mean over 89 subjects of the per-subject vertex-level Spearman $\rho$ = -0.3434 | dynamics__robustness__variants (outside FDR) (5) | robustness | subject (N=89) x vertex (4k fs_LR), n=89 subjects x 8004 vertices | hierarchical subject x vertex spin (N=1000, seed 0) | 0.027 | – | |
| Intrinsic timescale (1/e lag)<br>mean over 89 subjects of the per-subject vertex-level Spearman $\rho$ = -0.3147 | dynamics__robustness__variants (outside FDR) (5) | robustness | subject (N=89) x vertex (4k fs_LR), n=89 subjects x 8004 vertices | hierarchical subject x vertex spin (N=1000, seed 0) | 0.025 | – | |
| Intrinsic timescale (area under the ACF, pre-specified)<br>mean over 89 subjects of the per-subject vertex-level Spearman $\rho$ = -0.4592 | dynamics__singlemember_7 (7) | confirmatory | subject (N=89) x vertex (4k fs_LR), n=89 subjects x 8004 vertices | hierarchical subject x vertex spin (N=1000, seed 0) | 0.004 | 0.014 | * |
| Intrinsic timescale (exponential decay)<br>mean over 89 subjects of the per-subject vertex-level Spearman $\rho$ = -0.3797 | dynamics__robustness__variants (outside FDR) (5) | robustness | subject (N=89) x vertex (4k fs_LR), n=89 subjects x 8004 vertices | hierarchical subject x vertex spin (N=1000, seed 0) | 0.013 | – | |
| Aperiodic exponent (knee mode)<br>mean over 89 subjects of the per-subject vertex-level Spearman $\rho$ = +0.3388 | dynamics__robustness__variants (outside FDR) (5) | robustness | subject (N=89) x vertex (4k fs_LR), n=89 subjects x 8004 vertices | hierarchical subject x vertex spin (N=1000, seed 0) | 0.040 | – | |
| Aperiodic exponent (broadband 1-100 Hz)<br>mean over 89 subjects of the per-subject vertex-level Spearman $\rho$ = +0.0954 | dynamics__robustness__variants (outside FDR) (5) | robustness | subject (N=89) x vertex (4k fs_LR), n=89 subjects x 8004 vertices | hierarchical subject x vertex spin (N=1000, seed 0) | 0.469 | – | |
| Aperiodic-corrected delta power<br>mean over 89 subjects of the per-subject vertex-level Spearman $\rho$ = -0.0298 | osc__band__as__coded__4 (4) | exploratory | subject (N=89) x vertex (4k fs_LR), n=89 subjects x 8004 vertices | hierarchical subject x vertex spin (N=1000, seed 0) | 0.284 | 0.567 | |

| Measure | Family ($n$) | Status | Unit ($n_{\text{obs}}$) | Null | $p_{\text{raw}}$ | $q$ | |
|---|---|---|---|---|---|---|---|
| Aperiodic-corrected theta power mean over 89 subjects of the per-subject vertex-level Spearman $\rho$ = -0.0923 | osc__band__as__coded__4 (4) | exploratory | subject (N=89) x vertex (4k fs__LR), n=89 subjects x 8004 vertices | hierarchical subject x vertex spin (N=1000, seed 0) | 0.493 | 0.591 | |
| Aperiodic-corrected alpha power mean over 89 subjects of the per-subject vertex-level Spearman $\rho$ = +0.2602 | osc__band__as__coded__4 (4) | exploratory | subject (N=89) x vertex (4k fs__LR), n=89 subjects x 8004 vertices | hierarchical subject x vertex spin (N=1000, seed 0) | 0.242 | 0.567 | |
| Aperiodic-corrected beta power mean over 89 subjects of the per-subject vertex-level Spearman $\rho$ = +0.2222 | dynamics__singlemember__7 (7) | confirmatory | subject (N=89) x vertex (4k fs__LR), n=89 subjects x 8004 vertices | hierarchical subject x vertex spin (N=1000, seed 0) | 0.012 | 0.021 | * |
| Aperiodic-corrected gamma power mean over 89 subjects of the per-subject vertex-level Spearman $\rho$ = -0.0273 | osc__band__as__coded__4 (4) | exploratory | subject (N=89) x vertex (4k fs__LR), n=89 subjects x 8004 vertices | hierarchical subject x vertex spin (N=1000, seed 0) | 0.591 | 0.591 | |
| Slow/fast band-power ratio mean over 89 subjects of the per-subject vertex-level Spearman $\rho$ = -0.3564 | dynamics__singlemember__7 (7) | confirmatory | subject (N=89) x vertex (4k fs__LR), n=89 subjects x 8004 vertices | hierarchical subject x vertex spin (N=1000, seed 0) | 0.009 | 0.021 | * |
| Power-weighted spectral centroid mean over 89 subjects of the per-subject vertex-level Spearman $\rho$ = +0.4088 | dynamics__singlemember__7 (7) | confirmatory | subject (N=89) x vertex (4k fs__LR), n=89 subjects x 8004 vertices | hierarchical subject x vertex spin (N=1000, seed 0) | 0.004 | 0.014 | * |
| MOR (opioid, metabotropic) Spearman $\rho$ = -0.5483 | receptor__maps__19 (19) | exploratory | vertex (32k fs__LR), n=58731 | vertex spin (N=1000, seed 0) | 0.005 | 0.043 | * |
| 5-HT1a (serotonin, metabotropic) Spearman $\rho$ = -0.5370 | receptor__maps__19 (19) | exploratory | vertex (32k fs__LR), n=58731 | vertex spin (N=1000, seed 0) | 0.007 | 0.043 | * |
| CB1 (endocannabinoid, metabotropic) Spearman $\rho$ = -0.4058 | receptor__maps__19 (19) | exploratory | vertex (32k fs__LR), n=58731 | vertex spin (N=1000, seed 0) | 0.009 | 0.043 | * |
| D1 (dopamine, metabotropic) Spearman $\rho$ = -0.3935 | receptor__maps__19 (19) | exploratory | vertex (32k fs__LR), n=58731 | vertex spin (N=1000, seed 0) | 0.017 | 0.065 | |
| 5-HT4 (serotonin, metabotropic) Spearman $\rho$ = -0.3730 | receptor__maps__19 (19) | exploratory | vertex (32k fs__LR), n=58731 | vertex spin (N=1000, seed 0) | 0.048 | 0.152 | |
| D2 (dopamine, metabotropic) Spearman $\rho$ = -0.3552 | receptor__maps__19 (19) | exploratory | vertex (32k fs__LR), n=58731 | vertex spin (N=1000, seed 0) | 0.108 | 0.256 | |
| DAT (dopamine, transporter) Spearman $\rho$ = -0.3400 | receptor__maps__19 (19) | exploratory | vertex (32k fs__LR), n=58731 | vertex spin (N=1000, seed 0) | 0.061 | 0.165 | |
| mGluR5 (glutamate, metabotropic) Spearman $\rho$ = -0.2509 | receptor__maps__19 (19) | exploratory | vertex (32k fs__LR), n=58731 | vertex spin (N=1000, seed 0) | 0.127 | 0.268 | |
| H3 (histamine, metabotropic) Spearman $\rho$ = -0.2057 | receptor__maps__19 (19) | exploratory | vertex (32k fs__LR), n=58731 | vertex spin (N=1000, seed 0) | 0.277 | 0.478 | |
| 5-HT2a (serotonin, metabotropic) Spearman $\rho$ = -0.1763 | receptor__maps__19 (19) | exploratory | vertex (32k fs__LR), n=58731 | vertex spin (N=1000, seed 0) | 0.226 | 0.429 | |
| 5-HT6 (serotonin, metabotropic) Spearman $\rho$ = -0.1197 | receptor__maps__19 (19) | exploratory | vertex (32k fs__LR), n=58731 | vertex spin (N=1000, seed 0) | 0.443 | 0.647 | |
| 5-HTT (serotonin, transporter) Spearman $\rho$ = -0.0930 | receptor__maps__19 (19) | exploratory | vertex (32k fs__LR), n=58731 | vertex spin (N=1000, seed 0) | 0.665 | 0.843 | |

| Measure | Family ($n$) | Status | Unit ($n_{\mathrm{obs}}$) | Null | $p_{\mathrm{raw}}$ | $q$ | |
|---|---|---|---|---|---|---|---|
| NMDA (glutamate, ionotropic) Spearman $\rho$ = -0.0796 | receptor_maps_19 (19) | exploratory | vertex (32k fs_LR), n=58731 | vertex spin (N=1000, seed 0) | 0.348 | 0.550 | |
| a4b2 (acetylcholine, ionotropic) Spearman $\rho$ = -0.0186 | receptor_maps_19 (19) | exploratory | vertex (32k fs_LR), n=58731 | vertex spin (N=1000, seed 0) | 0.915 | 0.951 | |
| M1 (acetylcholine, metabotropic) Spearman $\rho$ = -0.0076 | receptor_maps_19 (19) | exploratory | vertex (32k fs_LR), n=58731 | vertex spin (N=1000, seed 0) | 0.951 | 0.951 | |
| VAChT (acetylcholine, transporter) Spearman $\rho$ = +0.0115 | receptor_maps_19 (19) | exploratory | vertex (32k fs_LR), n=58731 | vertex spin (N=1000, seed 0) | 0.943 | 0.951 | |
| GABA-A (GABA, ionotropic) Spearman $\rho$ = +0.0389 | receptor_maps_19 (19) | exploratory | vertex (32k fs_LR), n=58731 | vertex spin (N=1000, seed 0) | 0.818 | 0.951 | |
| 5-HT1b (serotonin, metabotropic) Spearman $\rho$ = +0.0953 | receptor_maps_19 (19) | exploratory | vertex (32k fs_LR), n=58731 | vertex spin (N=1000, seed 0) | 0.514 | 0.698 | |
| NET (noradrenaline, transporter) Spearman $\rho$ = +0.4413 | receptor_maps_19 (19) | exploratory | vertex (32k fs_LR), n=58731 | vertex spin (N=1000, seed 0) | 0.003 | 0.043 | * |
| Receptor composite: metabotropic Spearman $\rho$ = -0.4082 | receptor_composites_ 8 (8) | exploratory | vertex (32k fs_LR), n=58731 | vertex spin (N=1000, seed 0) | 0.016 | 0.043 | * |
| Receptor composite: ionotropic Spearman $\rho$ = -0.0346 | receptor_composites_ 8 (8) | exploratory | vertex (32k fs_LR), n=58731 | vertex spin (N=1000, seed 0) | 0.844 | 0.844 | |
| Receptor composite: serotonin Spearman $\rho$ = -0.2899 | receptor_composites_ 8 (8) | exploratory | vertex (32k fs_LR), n=58731 | vertex spin (N=1000, seed 0) | 0.108 | 0.216 | |
| Receptor composite: dopamine Spearman $\rho$ = -0.4603 | receptor_composites_ 8 (8) | exploratory | vertex (32k fs_LR), n=58731 | vertex spin (N=1000, seed 0) | 0.008 | 0.032 | * |
| Receptor composite: acetylcholine Spearman $\rho$ = -0.0255 | receptor_composites_ 8 (8) | exploratory | vertex (32k fs_LR), n=58731 | vertex spin (N=1000, seed 0) | 0.807 | 0.844 | |
| Receptor composite: reuptake_ innervation Spearman $\rho$ = +0.0370 | receptor_composites_ 8 (8) | exploratory | vertex (32k fs_LR), n=58731 | vertex spin (N=1000, seed 0) | 0.831 | 0.844 | |
| Receptor composite: iono_minus_ metabo_index Spearman $\rho$ = +0.4263 | receptor_composites_ 8 (8) | exploratory | vertex (32k fs_LR), n=58731 | vertex spin (N=1000, seed 0) | 0.002 | 0.016 | * |
| Receptor composite: glutamate_ minus_GABA Spearman $\rho$ = -0.1634 | receptor_composites_ 8 (8) | exploratory | vertex (32k fs_LR), n=58731 | vertex spin (N=1000, seed 0) | 0.184 | 0.294 | |
| Receptor diversity (Shannon entropy of 19 maps) Spearman $\rho$ = -0.3812 | fig9_predictions_3 (3) | confirmatory | vertex (32k fs_LR), n=58731 | vertex spin (N=1000, seed 0) | 0.046 | 0.069 | |
| Receptor diversity (coefficient-of-variation robustness check) Spearman $\rho$ = +0.4143 | no FDR family (robustness variant) (1) | robustness | vertex (32k fs_LR), n=58731 | vertex spin (N=1000, seed 0) | 0.033 | – | |
| RORB [gene, layer IV (granular)] Spearman $\rho$ = +0.7761 | layer_markers_8 (8) | confirmatory | parcel (Desikan-68), n=68 | parcel spin (N=1000, seed 0) | 0.003 | 0.005 | * |
| CUX2 [gene, layer II/III (upper)] Spearman $\rho$ = +0.7112 | layer_markers_8 (8) | exploratory | parcel (Desikan-68), n=68 | parcel spin (N=1000, seed 0) | 0.003 | 0.005 | * |
| FEZF2 [gene, layer V (deep)] Spearman $\rho$ = -0.5932 | layer_markers_8 (8) | exploratory | parcel (Desikan-68), n=68 | parcel spin (N=1000, seed 0) | 0.003 | 0.005 | * |
| FOXP2 [gene, layer VI (deep)] Spearman $\rho$ = +0.6890 | layer_markers_8 (8) | exploratory | parcel (Desikan-68), n=68 | parcel spin (N=1000, seed 0) | 0.004 | 0.005 | * |

| Measure | Family ($n$) | Status | Unit ($n_{obs}$) | Null | $p_{raw}$ | $q$ | |
|---|---|---|---|---|---|---|---|
| TLE4 [gene, layer VI (deep)]<br>Spearman $\rho$ = +0.2943 | layer_markers_8 (8) | exploratory | parcel (Desikan-68), n=68 | parcel spin (N=1000, seed 0) | 0.044 | 0.049 | * |
| upper(CUX2) [composite, layer -]<br>Spearman $\rho$ = +0.7112 | layer_markers_8 (8) | exploratory | parcel (Desikan-68), n=68 | parcel spin (N=1000, seed 0) | 0.003 | 0.005 | * |
| granular(RORB) [composite, layer -]<br>Spearman $\rho$ = +0.7761 | layer_markers_8 (8) | exploratory | parcel (Desikan-68), n=68 | parcel spin (N=1000, seed 0) | 0.003 | 0.005 | * |
| deep(FEZF2/FOXP2/TLE4) [composite, layer -]<br>Spearman $\rho$ = +0.3347 | layer_markers_8 (8) | exploratory | parcel (Desikan-68), n=68 | parcel spin (N=1000, seed 0) | 0.049 | 0.049 | * |
| Evolutionary cortical expansion (Xu 2020)<br>Spearman $\rho$ = -0.2659 | fig9_predictions_3 (3) | confirmatory | vertex (32k fs_LR), n=58731 | vertex spin (N=1000, seed 0) | 0.028 | 0.069 | |
| Developmental cortical expansion (Hill 2010)<br>Spearman $\rho$ = -0.1146 | fig9_predictions_3 (3) | confirmatory | vertex (32k fs_LR), n=58731 | vertex spin (N=1000, seed 0) | 0.416 | 0.416 | |
| ENIGMA cortical-thickness effect size: 22q<br>Spearman $\rho$ = +0.3557 | disease_8 (8) | exploratory | parcel (Desikan-68), n=68 | parcel spin (N=1000, seed 0) | 0.022 | 0.035 | * |
| ENIGMA 22q, partialling the functional gradient<br>partial Spearman $\rho$ = +0.3571 | disease_partial_8 (8) | robustness | parcel (Desikan-68), n=68 | parcel spin (N=1000, seed 0) | 0.026 | 0.069 | |
| ENIGMA cortical-thickness effect size: adhd<br>Spearman $\rho$ = +0.4136 | disease_8 (8) | exploratory | parcel (Desikan-68), n=68 | parcel spin (N=1000, seed 0) | 0.005 | 0.019 | * |
| ENIGMA adhd, partialling the functional gradient<br>partial Spearman $\rho$ = +0.5458 | disease_partial_8 (8) | robustness | parcel (Desikan-68), n=68 | parcel spin (N=1000, seed 0) | 0.003 | 0.016 | * |
| ENIGMA cortical-thickness effect size: asd<br>Spearman $\rho$ = -0.0249 | disease_8 (8) | exploratory | parcel (Desikan-68), n=68 | parcel spin (N=1000, seed 0) | 0.949 | 0.988 | |
| ENIGMA asd, partialling the functional gradient<br>partial Spearman $\rho$ = +0.1804 | disease_partial_8 (8) | robustness | parcel (Desikan-68), n=68 | parcel spin (N=1000, seed 0) | 0.625 | 0.625 | |
| ENIGMA cortical-thickness effect size: bipolar<br>Spearman $\rho$ = +0.0033 | disease_8 (8) | exploratory | parcel (Desikan-68), n=68 | parcel spin (N=1000, seed 0) | 0.988 | 0.988 | |
| ENIGMA bipolar, partialling the functional gradient<br>partial Spearman $\rho$ = -0.2483 | disease_partial_8 (8) | robustness | parcel (Desikan-68), n=68 | parcel spin (N=1000, seed 0) | 0.154 | 0.246 | |
| ENIGMA cortical-thickness effect size: depression<br>Spearman $\rho$ = +0.5972 | disease_8 (8) | exploratory | parcel (Desikan-68), n=68 | parcel spin (N=1000, seed 0) | 0.004 | 0.019 | * |
| ENIGMA depression, partialling the functional gradient<br>partial Spearman $\rho$ = +0.4567 | disease_partial_8 (8) | robustness | parcel (Desikan-68), n=68 | parcel spin (N=1000, seed 0) | 0.004 | 0.016 | * |
| ENIGMA cortical-thickness effect size: epilepsy<br>Spearman $\rho$ = -0.4229 | disease_8 (8) | exploratory | parcel (Desikan-68), n=68 | parcel spin (N=1000, seed 0) | 0.019 | 0.035 | * |

| Measure | Family ($n$) | Status | Unit ($n_{\mathrm{obs}}$) | Null | $p_{\mathrm{raw}}$ | $q$ | |
|---|---|---|---|---|---|---|---|
| ENIGMA epilepsy, partialling the functional gradient<br>partial Spearman $\rho$ = -0.3570 | disease_partial_8 (8) | robustness | parcel (Desikan-68), n=68 | parcel spin (N=1000, seed 0) | 0.043 | 0.086 | |
| ENIGMA cortical-thickness effect size: ocd<br>Spearman $\rho$ = +0.3172 | disease_8 (8) | exploratory | parcel (Desikan-68), n=68 | parcel spin (N=1000, seed 0) | 0.007 | 0.019 | * |
| ENIGMA ocd, partialling the functional gradient<br>partial Spearman $\rho$ = +0.1240 | disease_partial_8 (8) | robustness | parcel (Desikan-68), n=68 | parcel spin (N=1000, seed 0) | 0.342 | 0.456 | |
| ENIGMA cortical-thickness effect size: schizophrenia<br>Spearman $\rho$ = -0.1816 | disease_8 (8) | exploratory | parcel (Desikan-68), n=68 | parcel spin (N=1000, seed 0) | 0.502 | 0.670 | |
| ENIGMA schizophrenia, partialling the functional gradient<br>partial Spearman $\rho$ = -0.2176 | disease_partial_8 (8) | robustness | parcel (Desikan-68), n=68 | parcel spin (N=1000, seed 0) | 0.422 | 0.482 | |
| Short-range connectivity vs type-distance (log-linear slope)<br>log-linear slope = -1.2306 | tractography_5 (5) | confirmatory | type-pair (7x7), n=21 | label permutation of the 7 type ranks | 0.003 | – | * |
| Short-range connectivity vs type-distance, contact area partialled out<br>partial Pearson r = -0.6358 | tractography_5 (5) | confirmatory | type-pair (7x7), n=21 | parametric p on the partial Pearson r | 0.002 | – | * |
| Per-type connectional property: degree_short<br>Spearman $\rho$ = +0.1071 | tractography_per_type_8 (no FDR applied) (8) | exploratory | type (7), n=7 | label permutation of the 7 type ranks (N=5040, exhaustive) | 0.840 | – | |
| Per-type connectional property: degree_long<br>Spearman $\rho$ = +0.4286 | tractography_per_type_8 (no FDR applied) (8) | exploratory | type (7), n=7 | label permutation of the 7 type ranks (N=5040, exhaustive) | 0.354 | – | |
| Per-type connectional property: degree_total<br>Spearman $\rho$ = +0.1786 | tractography_per_type_8 (no FDR applied) (8) | exploratory | type (7), n=7 | label permutation of the 7 type ranks (N=5040, exhaustive) | 0.713 | – | |
| Per-type connectional property: degree_per_vertex<br>Spearman $\rho$ = -0.8214 | tractography_per_type_8 (no FDR applied) (8) | exploratory | type (7), n=7 | label permutation of the 7 type ranks (N=5040, exhaustive) | 0.034 | – | * |
| Per-type connectional property: degree_short_per_vertex<br>Spearman $\rho$ = -0.8929 | tractography_per_type_8 (no FDR applied) (8) | exploratory | type (7), n=7 | label permutation of the 7 type ranks (N=5040, exhaustive) | 0.012 | – | * |
| Per-type connectional property: degree_long_per_vertex<br>Spearman $\rho$ = -0.8214 | tractography_per_type_8 (no FDR applied) (8) | exploratory | type (7), n=7 | label permutation of the 7 type ranks (N=5040, exhaustive) | 0.034 | – | * |
| Per-type connectional property: mean_length_mm<br>Spearman $\rho$ = +0.9643 | tractography_per_type_8 (no FDR applied) (8) | exploratory | type (7), n=7 | label permutation of the 7 type ranks (N=5040, exhaustive) | 0.003 | – | * |
| Per-type connectional property: mean_tortuosity<br>Spearman $\rho$ = +1.0000 | tractography_per_type_8 (no FDR applied) (8) | exploratory | type (7), n=7 | label permutation of the 7 type ranks (N=5040, exhaustive) | 0.0004 | – | * |

| Measure | Family ($n$) | Status | Unit ($n_{\mathrm{obs}}$) | Null | $p_{\mathrm{raw}}$ | $q$ | |
|---|---|---|---|---|---|---|---|
| Added value over the area-level map, global: T1w/T2w myelin delta $\|\rho\|$ = +0.1164 (cyto7 +0.608 vs von-Economo +0.492) | added_value_global_9 (9) | confirmatory | vertex (32k fs_LR), n=58731 | vertex spin (N=1000, seed 0) | 0.001 | 0.006 | * |
| Added value over the area-level map, global: Cortical thickness delta $\|\rho\|$ = +0.0960 (cyto7 -0.509 vs von-Economo -0.413) | added_value_global_9 (9) | confirmatory | vertex (32k fs_LR), n=58731 | vertex spin (N=1000, seed 0) | 0.002 | 0.006 | * |
| Added value over the area-level map, global: Intrinsic timescale delta $\|\rho\|$ = +0.1582 (cyto7 -0.487 vs von-Economo -0.329) | added_value_global_9 (9) | confirmatory | vertex (32k fs_LR), n=58731 | vertex spin (N=1000, seed 0) | 0.034 | 0.077 | |
| Added value over the area-level map, global: Delta power (0.5-4 Hz) delta $\|\rho\|$ = +0.1398 (cyto7 -0.448 vs von-Economo -0.308) | added_value_global_9 (9) | confirmatory | vertex (32k fs_LR), n=58731 | vertex spin (N=1000, seed 0) | 0.077 | 0.138 | |
| Added value over the area-level map, global: Theta power (4-8 Hz) delta $\|\rho\|$ = +0.0662 (cyto7 -0.213 vs von-Economo -0.147) | added_value_global_9 (9) | confirmatory | vertex (32k fs_LR), n=58731 | vertex spin (N=1000, seed 0) | 0.537 | 0.537 | |
| Added value over the area-level map, global: Alpha power (8-12 Hz) delta $\|\rho\|$ = +0.1074 (cyto7 +0.371 vs von-Economo +0.263) | added_value_global_9 (9) | confirmatory | vertex (32k fs_LR), n=58731 | vertex spin (N=1000, seed 0) | 0.226 | 0.298 | |
| Added value over the area-level map, global: Beta power (12-30 Hz) delta $\|\rho\|$ = +0.0948 (cyto7 +0.235 vs von-Economo +0.141) | added_value_global_9 (9) | confirmatory | vertex (32k fs_LR), n=58731 | vertex spin (N=1000, seed 0) | 0.446 | 0.501 | |
| Added value over the area-level map, global: Gamma power (30-60 Hz) delta $\|\rho\|$ = +0.0968 (cyto7 -0.325 vs von-Economo -0.228) | added_value_global_9 (9) | confirmatory | vertex (32k fs_LR), n=58731 | vertex spin (N=1000, seed 0) | 0.232 | 0.298 | |
| Added value over the area-level map, global: Principal functional gradient delta $\|\rho\|$ = +0.1620 (cyto7 -0.583 vs von-Economo -0.421) | added_value_global_9 (9) | confirmatory | vertex (32k fs_LR), n=58731 | vertex spin (N=1000, seed 0) | 0.002 | 0.006 | * |
| Win fraction on the disagreement set (all): T1w/T2w myelin win fraction = 0.6156 | added_value_localized_27 (descriptive, not tested) (27) | descriptive | vertex (32k fs_LR), n=27298 | none; reported descriptively | – | – | |
| Win fraction on the disagreement set (off1): T1w/T2w myelin win fraction = 0.5790 | added_value_localized_27 (descriptive, not tested) (27) | descriptive | vertex (32k fs_LR), n=23168 | none; reported descriptively | – | – | |
| Win fraction on the disagreement set (off2): T1w/T2w myelin win fraction = 0.8206 | added_value_localized_27 (descriptive, not tested) (27) | descriptive | vertex (32k fs_LR), n=4130 | none; reported descriptively | – | – | |

| Measure | Family ($n$) | Status | Unit ($n_{\text{obs}}$) | Null | $p_{\text{raw}}$ | $q$ |
|---|---|---|---|---|---|---|
| Win fraction on the disagreement set (all): Cortical thickness win fraction = 0.5682 | added_value_ localized_27 (descriptive, not tested) (27) | descriptive | vertex (32k fs_LR), n=27298 | none; reported descriptively | – | – |
| Win fraction on the disagreement set (off1): Cortical thickness win fraction = 0.5508 | added_value_ localized_27 (descriptive, not tested) (27) | descriptive | vertex (32k fs_LR), n=23168 | none; reported descriptively | – | – |
| Win fraction on the disagreement set (off2): Cortical thickness win fraction = 0.6656 | added_value_ localized_27 (descriptive, not tested) (27) | descriptive | vertex (32k fs_LR), n=4130 | none; reported descriptively | – | – |
| Win fraction on the disagreement set (all): Intrinsic timescale win fraction = 0.6377 | added_value_ localized_27 (descriptive, not tested) (27) | descriptive | vertex (32k fs_LR), n=27298 | none; reported descriptively | – | – |
| Win fraction on the disagreement set (off1): Intrinsic timescale win fraction = 0.6171 | added_value_ localized_27 (descriptive, not tested) (27) | descriptive | vertex (32k fs_LR), n=23168 | none; reported descriptively | – | – |
| Win fraction on the disagreement set (off2): Intrinsic timescale win fraction = 0.7530 | added_value_ localized_27 (descriptive, not tested) (27) | descriptive | vertex (32k fs_LR), n=4130 | none; reported descriptively | – | – |
| Win fraction on the disagreement set (all): Delta power (0.5-4 Hz) win fraction = 0.6062 | added_value_ localized_27 (descriptive, not tested) (27) | descriptive | vertex (32k fs_LR), n=27298 | none; reported descriptively | – | – |
| Win fraction on the disagreement set (off1): Delta power (0.5-4 Hz) win fraction = 0.5902 | added_value_ localized_27 (descriptive, not tested) (27) | descriptive | vertex (32k fs_LR), n=23168 | none; reported descriptively | – | – |
| Win fraction on the disagreement set (off2): Delta power (0.5-4 Hz) win fraction = 0.6961 | added_value_ localized_27 (descriptive, not tested) (27) | descriptive | vertex (32k fs_LR), n=4130 | none; reported descriptively | – | – |
| Win fraction on the disagreement set (all): Theta power (4-8 Hz) win fraction = 0.5192 | added_value_ localized_27 (descriptive, not tested) (27) | descriptive | vertex (32k fs_LR), n=27298 | none; reported descriptively | – | – |
| Win fraction on the disagreement set (off1): Theta power (4-8 Hz) win fraction = 0.4988 | added_value_ localized_27 (descriptive, not tested) (27) | descriptive | vertex (32k fs_LR), n=23168 | none; reported descriptively | – | – |
| Win fraction on the disagreement set (off2): Theta power (4-8 Hz) win fraction = 0.6332 | added_value_ localized_27 (descriptive, not tested) (27) | descriptive | vertex (32k fs_LR), n=4130 | none; reported descriptively | – | – |

| Measure | Family ($n$) | Status | Unit ($n_{\mathrm{obs}}$) | Null | $p_{\mathrm{raw}}$ | $q$ |
|---|---|---|---|---|---|---|
| Win fraction on the disagreement set (all): Alpha power (8-12 Hz) win fraction = 0.5435 | added_value_localized_27 (descriptive, not tested) (27) | descriptive | vertex (32k fs_LR), n=27298 | none; reported descriptively | – | – |
| Win fraction on the disagreement set (off1): Alpha power (8-12 Hz) win fraction = 0.5497 | added_value_localized_27 (descriptive, not tested) (27) | descriptive | vertex (32k fs_LR), n=23168 | none; reported descriptively | – | – |
| Win fraction on the disagreement set (off2): Alpha power (8-12 Hz) win fraction = 0.5085 | added_value_localized_27 (descriptive, not tested) (27) | descriptive | vertex (32k fs_LR), n=4130 | none; reported descriptively | – | – |
| Win fraction on the disagreement set (all): Beta power (12-30 Hz) win fraction = 0.4824 | added_value_localized_27 (descriptive, not tested) (27) | descriptive | vertex (32k fs_LR), n=27298 | none; reported descriptively | – | – |
| Win fraction on the disagreement set (off1): Beta power (12-30 Hz) win fraction = 0.4532 | added_value_localized_27 (descriptive, not tested) (27) | descriptive | vertex (32k fs_LR), n=23168 | none; reported descriptively | – | – |
| Win fraction on the disagreement set (off2): Beta power (12-30 Hz) win fraction = 0.6462 | added_value_localized_27 (descriptive, not tested) (27) | descriptive | vertex (32k fs_LR), n=4130 | none; reported descriptively | – | – |
| Win fraction on the disagreement set (all): Gamma power (30-60 Hz) win fraction = 0.5203 | added_value_localized_27 (descriptive, not tested) (27) | descriptive | vertex (32k fs_LR), n=27298 | none; reported descriptively | – | – |
| Win fraction on the disagreement set (off1): Gamma power (30-60 Hz) win fraction = 0.5203 | added_value_localized_27 (descriptive, not tested) (27) | descriptive | vertex (32k fs_LR), n=23168 | none; reported descriptively | – | – |
| Win fraction on the disagreement set (off2): Gamma power (30-60 Hz) win fraction = 0.5203 | added_value_localized_27 (descriptive, not tested) (27) | descriptive | vertex (32k fs_LR), n=4130 | none; reported descriptively | – | – |
| Win fraction on the disagreement set (all): Principal functional gradient win fraction = 0.6193 | added_value_localized_27 (descriptive, not tested) (27) | descriptive | vertex (32k fs_LR), n=27298 | none; reported descriptively | – | – |
| Win fraction on the disagreement set (off1): Principal functional gradient win fraction = 0.5972 | added_value_localized_27 (descriptive, not tested) (27) | descriptive | vertex (32k fs_LR), n=23168 | none; reported descriptively | – | – |
| Win fraction on the disagreement set (off2): Principal functional gradient win fraction = 0.7436 | added_value_localized_27 (descriptive, not tested) (27) | descriptive | vertex (32k fs_LR), n=4130 | none; reported descriptively | – | – |

| Measure | Family ($n$) | Status | Unit ($n_{\mathrm{obs}}$) | Null | $p_{\mathrm{raw}}$ | $q$ | |
|---|---|---|---|---|---|---|---|
| Aggregate win fraction on the disagreement set (all)<br>win fraction = 0.6010 across myelin/thickness/gradient | added_value_aggregate_3 (3) | confirmatory | vertex (32k fs_LR), n=27298 | vertex spin (N=1000, seed 0), win fraction recomputed per rotation | 0.001 | – | * |
| Aggregate win fraction on the disagreement set (off-by-1)<br>win fraction = 0.5757 across myelin/thickness/gradient | added_value_aggregate_3 (3) | confirmatory | vertex (32k fs_LR), n=23168 | vertex spin (N=1000, seed 0), win fraction recomputed per rotation | 0.002 | – | * |
| Aggregate win fraction on the disagreement set (off-by-$\geq$2)<br>win fraction = 0.7433 across myelin/thickness/gradient | added_value_aggregate_3 (3) | confirmatory | vertex (32k fs_LR), n=4130 | vertex spin (N=1000, seed 0), win fraction recomputed per rotation | 0.001 | – | * |
| Added value over the area-level map with BigBrain histology as arbiter<br>delta $\|\rho\|$ = +0.0515 (cyto7 +0.209 vs von-Economo +0.158) | bigbrain_added_value_2 (2) | confirmatory | vertex (32k fs_LR), n=58731 | vertex spin (N=1000, seed 0) | 0.343 | 0.685 | |
| Win fraction on the disagreement set with BigBrain histology as arbiter<br>win fraction = 0.4990 (chance 0.519) | bigbrain_added_value_2 (2) | confirmatory | vertex (32k fs_LR), n=27298 | vertex spin (N=1000, seed 0), win fraction recomputed per rotation | 0.740 | 0.740 | |
| Negative control: Sulcal depth (pre-specified expected-null)<br>Spearman $\rho$ = -0.0392 | negative_control_2 (2) | confirmatory | vertex (32k fs_LR), n=58731 | vertex spin (N=1000, seed 0) | 0.661 | 0.661 | |
| Negative control: Cortical curvature (\|mean-curv\| proxy)<br>Spearman $\rho$ = -0.0515 | negative_control_2 (2) | confirmatory | vertex (32k fs_LR), n=58731 | vertex spin (N=1000, seed 0) | 0.529 | 0.661 | |
| Short-range connectivity vs type-distance, topology-preserving null<br>Pearson $\rho$ = -0.7649 (null mean -0.3337) | tractography_5 (5) | confirmatory | type pair (7x7 off-diagonal), n=21, n=21 | type-map rotation (N=1000, seed 0), scale-free statistic | 0.040 | – | |
| Short-range connectivity vs type-distance, bundle level, endpoint-pair offset<br>beta = +0.0089 per ordinal step | tractography_5 (5) | confirmatory | bundle x type pair, n=924, n=924 | negative binomial, tract-clustered errors | 0.957 | – | |
| Short-range connectivity vs type-distance, bundle level, 80 mm opportunity offset<br>beta = +0.1263 per ordinal step | tractography_5 (5) | confirmatory | bundle x type pair, n=924, n=924 | negative binomial, tract-clustered errors | 0.433 | – | |
| Win fraction vs the granularity-matched null (200 size-matched partitions)<br>win fraction = 0.6010 (matched-null mean 0.5422, s.d. 0.0298, 98.5th percentile) | benchmark_controls_4 (4) | robustness | vertex (32k fs_LR), n=27298 | 200 contiguous size-matched seven-class partitions | 0.020 | – | * |
| Win fraction of the myelin septile as challenger against the area-level map<br>win fraction = 0.7026 | benchmark_controls_4 (4) | descriptive | vertex (32k fs_LR), n=27298 | none (descriptive comparator on the same disagreement set) | – | – | |

| Measure | Family ($n$) | Status | Unit ($n_{\mathrm{obs}}$) | Null | $p_{\mathrm{raw}}$ | $q$ |
|---|---|---|---|---|---|---|
| Win fraction of the gradient septile as challenger against the area-level map win fraction = 0.7139 | benchmark_controls_ 4 (4) | descriptive | vertex (32k fs_LR), n=27298 | none (descriptive comparator on the same disagreement set) | – | – |
| Win fraction of the BigBrain Hist-G1 septile as challenger against the area-level map win fraction = 0.5016 | benchmark_controls_ 4 (4) | descriptive | vertex (32k fs_LR), n=27298 | none (descriptive comparator on the same disagreement set) | – | – |

**Supplementary Table S9.** Robustness and sensitivity analyses reported outside the declared FDR families. Each row re-estimates a result reported in the text under a different exclusion, covariate set, partition or model specification. **What this table is not:** it is not part of the census. The 147 entries of the 21 declared families are in Table S8; these 107 analyses are additional to them and are counted nowhere in that total. **No $q$ is reported in this table**, because these analyses ask whether a result is *stable* rather than whether an effect is *present*, and correcting a sensitivity analysis against the test it perturbs would in most cases correct a statistic against itself. That is a reporting choice, not a claim that multiplicity never applies to a robustness check: where such a check is used to support a conclusion of its own it should be corrected, and the comparator correlations are the one case here where it was. Those were given Benjamini–Hochberg control within each partition when they were computed (Annex J); their $q$ values are not shown here but are released in `comparator_panel.csv`. Raw $p$ is given wherever the analysis has a null of its own, and omitted where it has none. **Rows are not tests.** The 260 output rows behind this listing reduce to 107 distinct tests under three rules: the arm of a comparison that *is* the published result is a member of a declared family and is not counted again; per-subject rows are inputs to one group-level test rather than one test each; and a row with no null is a description. Per group: *Precentral (M1) exclusion* (8 rows → 4): 4 features x 2 masks; the with-M1 arm is the published test. *Agranular exclusion, myelin regression* (3 rows → 1): 3 nested arms; arm A is Table S8's myelin row and arm B is counted under allocortex exclusion. *Allocortex-exclusion re-estimates* (22 rows → 13): one re-estimate per measure; three overlapping files de-duplicated, published arm excluded. *Belt exclusions (Glasser-decisive, whole belt)* (25 rows → 25): one re-estimate per measure per belt definition; nothing reduces. *Partialling controls* (100 rows → 7): 89 of the rows are the 89 subjects feeding the group tests, not tests; 4 unadjusted outcomes are already in the dynamics family. *Tractography specifications* (14 rows → 13): 2 levels x 7 fitted specifications, each its own null; bundle/R1 is already in Table S8. *Hemisphere holdout and areal-interior win fractions* (40 rows → 8): published and per-arbiter rows are in declared families, 2 rows are derived means with no null, and the 24 hop-distance rows are a curve with no p. *Comparator correlations (Tables S6, S7)* (48 rows → 36): 12 features x 4 partitions; the cyto7 partition is the published test.

| What was tested | § | The analysis it varies | Unit ($n_{\mathrm{obs}}$) | Null | $p_{\mathrm{raw}}$ |
|---|---|---|---|---|---|
| T1w/T2w myelin, precentral territory excluded Spearman $\rho$ = +0.5671 | 3.5 | re-estimate without M1 (published $\rho$ +0.5894) | vertex (32k fs_LR), n=54696 | vertex spin (N=1000, seed 0) | 0.002 |
| Cortical thickness, precentral territory excluded Spearman $\rho$ = -0.5374 | 3.5 | re-estimate without M1 (published $\rho$ -0.5018) | vertex (32k fs_LR), n=54696 | vertex spin (N=1000, seed 0) | 0.002 |
| Principal functional gradient, precentral territory excluded Spearman $\rho$ = -0.5433 | 3.5 | re-estimate without M1 (published $\rho$ -0.5761) | vertex (32k fs_LR), n=54696 | vertex spin (N=1000, seed 0) | 0.002 |
| AHBA gene expression PC1, precentral territory excluded Spearman $\rho$ = +0.6550 | 3.5 | re-estimate without M1 (published $\rho$ +0.6657) | vertex (32k fs_LR), n=53922 | vertex spin (N=1000, seed 0) | 0.002 |
| T1w/T2w myelin, agranular excluded as well as allocortex Spearman $\rho$ = +0.5851 | 3.5 | re-estimate over types 3-7. The other two arms of the same comparison: types 2-7 (allocortex excluded) $\rho$ +0.6039, n=57952, listed below; types 1-7 (all types) $\rho$ +0.5894, n=58731, which is the published correlation and is Table S8's myelin row | vertex (32k fs_LR), n=55686 | vertex spin (N=1000, seed 0) | 0.001 |
| T1w/T2w myelin, allocortex excluded Spearman $\rho$ = +0.6039 | 2.6 / 3.5 | re-estimate on labelled cortex minus allocortex (declared family: structure_function_9) | vertex (32k fs_LR), n=57952 | vertex spin (N=1000, seed 0) | 0.001 |

| What was tested | § | The analysis it varies | Unit ($n_{\text{obs}}$) | Null | $p_{\text{raw}}$ |
|---|---|---|---|---|---|
| Cortical thickness, allocortex excluded Spearman $\rho$ = -0.5014 | 2.6 / 3.5 | re-estimate on labelled cortex minus allocortex (declared family: structure_function_9) | vertex (32k fs_LR), n=57952 | vertex spin (N=1000, seed 0) | 0.002 |
| Intrinsic timescale, allocortex excluded Spearman $\rho$ = -0.4879 | 2.6 / 3.5 | re-estimate on labelled cortex minus allocortex (declared family: structure_function_9) | vertex (32k fs_LR), n=57952 | vertex spin (N=1000, seed 0) | 0.050 |
| Delta power (0.5-4 Hz), allocortex excluded Spearman $\rho$ = -0.4477 | 2.6 / 3.5 | re-estimate on labelled cortex minus allocortex (declared family: structure_function_9) | vertex (32k fs_LR), n=57952 | vertex spin (N=1000, seed 0) | 0.113 |
| Theta power (4-8 Hz), allocortex excluded Spearman $\rho$ = -0.2048 | 2.6 / 3.5 | re-estimate on labelled cortex minus allocortex (declared family: structure_function_9) | vertex (32k fs_LR), n=57952 | vertex spin (N=1000, seed 0) | 0.575 |
| Alpha power (8-12 Hz), allocortex excluded Spearman $\rho$ = +0.3662 | 2.6 / 3.5 | re-estimate on labelled cortex minus allocortex (declared family: structure_function_9) | vertex (32k fs_LR), n=57952 | vertex spin (N=1000, seed 0) | 0.269 |
| Beta power (12-30 Hz), allocortex excluded Spearman $\rho$ = +0.2425 | 2.6 / 3.5 | re-estimate on labelled cortex minus allocortex (declared family: structure_function_9) | vertex (32k fs_LR), n=57952 | vertex spin (N=1000, seed 0) | 0.465 |
| Gamma power (30-60 Hz), allocortex excluded Spearman $\rho$ = -0.3190 | 2.6 / 3.5 | re-estimate on labelled cortex minus allocortex (declared family: structure_function_9) | vertex (32k fs_LR), n=57952 | vertex spin (N=1000, seed 0) | 0.274 |
| Principal functional gradient, allocortex excluded Spearman $\rho$ = -0.5822 | 2.6 / 3.5 | re-estimate on labelled cortex minus allocortex (declared family: structure_function_9) | vertex (32k fs_LR), n=57952 | vertex spin (N=1000, seed 0) | 0.001 |
| AHBA gene expression PC1, allocortex excluded Spearman $\rho$ = +0.6601 | 2.6 / 3.5 | re-estimate on labelled cortex minus allocortex (declared family: external_reference_2) | vertex (32k fs_LR), n=57494 | vertex spin (N=1000, seed 0) | 0.001 |
| PET receptor PC1 (Hansen-2022 collection), allocortex excluded Spearman $\rho$ = -0.3628 | 2.6 / 3.5 | re-estimate on labelled cortex minus allocortex (declared family: external_reference_2) | vertex (32k fs_LR), n=57952 | vertex spin (N=1000, seed 0) | 0.038 |
| BigBrain profile skewness (pre-specified, histology), allocortex excluded Spearman $\rho$ = -0.2057 | 2.6 / 3.5 | re-estimate on labelled cortex minus allocortex (declared family: bigbrain_profile_panel_5) | vertex (32k fs_LR), n=57952 | vertex spin (N=1000, seed 0) | 0.346 |
| MEG intrinsic timescale (int_area), allocortex excluded mean per-subject Spearman $\rho$ = -0.4510 | 2.6 / 3.5 | re-estimate on the 4k mesh minus allocortex | subject (N=89) x vertex (4k fs_LR), n=89 subjects x 7143 vertices | hierarchical subject x vertex spin (N=1000, seed 0) | 0.005 |
| T1w/T2w myelin, limbic belt excluded, Set A (whole belt) Spearman $\rho$ = +0.5856 | 4.3 | re-estimate with the belt removed (Set A (whole belt)) | vertex (32k fs_LR), n=50682 | vertex spin (N=1000, seed 0) | 0.001 |
| Cortical thickness, limbic belt excluded, Set A (whole belt) Spearman $\rho$ = -0.4349 | 4.3 | re-estimate with the belt removed (Set A (whole belt)) | vertex (32k fs_LR), n=50682 | vertex spin (N=1000, seed 0) | 0.008 |
| Intrinsic timescale, limbic belt excluded, Set A (whole belt) Spearman $\rho$ = -0.4343 | 4.3 | re-estimate with the belt removed (Set A (whole belt)) | vertex (32k fs_LR), n=50682 | vertex spin (N=1000, seed 0) | 0.124 |
| Delta power (0.5-4 Hz), limbic belt excluded, Set A (whole belt) Spearman $\rho$ = -0.3722 | 4.3 | re-estimate with the belt removed (Set A (whole belt)) | vertex (32k fs_LR), n=50682 | vertex spin (N=1000, seed 0) | 0.261 |

| What was tested | § | The analysis it varies | Unit ($n_{\mathrm{obs}}$) | Null | $p_{\mathrm{raw}}$ |
|---|---|---|---|---|---|
| Theta power (4-8 Hz), limbic belt excluded, Set A (whole belt)<br>Spearman $\rho$ = -0.2090 | 4.3 | re-estimate with the belt removed (Set A (whole belt)) | vertex (32k fs_LR), n=50682 | vertex spin (N=1000, seed 0) | 0.587 |
| Alpha power (8-12 Hz), limbic belt excluded, Set A (whole belt)<br>Spearman $\rho$ = +0.2678 | 4.3 | re-estimate with the belt removed (Set A (whole belt)) | vertex (32k fs_LR), n=50682 | vertex spin (N=1000, seed 0) | 0.488 |
| Beta power (12-30 Hz), limbic belt excluded, Set A (whole belt)<br>Spearman $\rho$ = +0.1822 | 4.3 | re-estimate with the belt removed (Set A (whole belt)) | vertex (32k fs_LR), n=50682 | vertex spin (N=1000, seed 0) | 0.600 |
| Gamma power (30-60 Hz), limbic belt excluded, Set A (whole belt)<br>Spearman $\rho$ = -0.2375 | 4.3 | re-estimate with the belt removed (Set A (whole belt)) | vertex (32k fs_LR), n=50682 | vertex spin (N=1000, seed 0) | 0.496 |
| Principal functional gradient, limbic belt excluded, Set A (whole belt)<br>Spearman $\rho$ = -0.6177 | 4.3 | re-estimate with the belt removed (Set A (whole belt)) | vertex (32k fs_LR), n=50682 | vertex spin (N=1000, seed 0) | 0.001 |
| T1w/T2w myelin, limbic belt excluded, Set B (Glasser-decisive)<br>Spearman $\rho$ = +0.6060 | 4.3 | re-estimate with the belt removed (Set B (Glasser-decisive)) | vertex (32k fs_LR), n=57730 | vertex spin (N=1000, seed 0) | 0.001 |
| Cortical thickness, limbic belt excluded, Set B (Glasser-decisive)<br>Spearman $\rho$ = -0.5075 | 4.3 | re-estimate with the belt removed (Set B (Glasser-decisive)) | vertex (32k fs_LR), n=57730 | vertex spin (N=1000, seed 0) | 0.002 |
| Intrinsic timescale, limbic belt excluded, Set B (Glasser-decisive)<br>Spearman $\rho$ = -0.4846 | 4.3 | re-estimate with the belt removed (Set B (Glasser-decisive)) | vertex (32k fs_LR), n=57730 | vertex spin (N=1000, seed 0) | 0.051 |
| Delta power (0.5-4 Hz), limbic belt excluded, Set B (Glasser-decisive)<br>Spearman $\rho$ = -0.4464 | 4.3 | re-estimate with the belt removed (Set B (Glasser-decisive)) | vertex (32k fs_LR), n=57730 | vertex spin (N=1000, seed 0) | 0.113 |
| Theta power (4-8 Hz), limbic belt excluded, Set B (Glasser-decisive)<br>Spearman $\rho$ = -0.2084 | 4.3 | re-estimate with the belt removed (Set B (Glasser-decisive)) | vertex (32k fs_LR), n=57730 | vertex spin (N=1000, seed 0) | 0.571 |
| Alpha power (8-12 Hz), limbic belt excluded, Set B (Glasser-decisive)<br>Spearman $\rho$ = +0.3730 | 4.3 | re-estimate with the belt removed (Set B (Glasser-decisive)) | vertex (32k fs_LR), n=57730 | vertex spin (N=1000, seed 0) | 0.256 |
| Beta power (12-30 Hz), limbic belt excluded, Set B (Glasser-decisive)<br>Spearman $\rho$ = +0.2408 | 4.3 | re-estimate with the belt removed (Set B (Glasser-decisive)) | vertex (32k fs_LR), n=57730 | vertex spin (N=1000, seed 0) | 0.469 |
| Gamma power (30-60 Hz), limbic belt excluded, Set B (Glasser-decisive)<br>Spearman $\rho$ = -0.3196 | 4.3 | re-estimate with the belt removed (Set B (Glasser-decisive)) | vertex (32k fs_LR), n=57730 | vertex spin (N=1000, seed 0) | 0.276 |
| Principal functional gradient, limbic belt excluded, Set B (Glasser-decisive)<br>Spearman $\rho$ = -0.5871 | 4.3 | re-estimate with the belt removed (Set B (Glasser-decisive)) | vertex (32k fs_LR), n=57730 | vertex spin (N=1000, seed 0) | 0.001 |
| AHBA gene expression PC1, limbic belt excluded, Set A (whole belt)<br>Spearman $\rho$ = +0.5821 | 4.3 | re-estimate with the belt removed (Set A (whole belt)) | vertex (32k fs_LR), n=50682 | vertex spin (N=1000, seed 0) | 0.004 |

| What was tested | § | The analysis it varies | Unit ($n_{\mathrm{obs}}$) | Null | $p_{\mathrm{raw}}$ |
|---|---|---|---|---|---|
| PET receptor PC1 (Hansen-2022 collection), limbic belt excluded, Set A (whole belt)<br>Spearman $\rho$ = -0.3652 | 4.3 | re-estimate with the belt removed (Set A (whole belt)) | vertex (32k fs_LR), n=50682 | vertex spin (N=1000, seed 0) | 0.070 |
| BigBrain profile skewness (pre-specified, histology), limbic belt excluded, Set A (whole belt)<br>Spearman $\rho$ = -0.2080 | 4.3 | re-estimate with the belt removed (Set A (whole belt)) | vertex (32k fs_LR), n=50682 | vertex spin (N=1000, seed 0) | 0.384 |
| AHBA gene expression PC1, limbic belt excluded, Set B (Glasser-decisive)<br>Spearman $\rho$ = +0.6648 | 4.3 | re-estimate with the belt removed (Set B (Glasser-decisive)) | vertex (32k fs_LR), n=57269 | vertex spin (N=1000, seed 0) | 0.001 |
| PET receptor PC1 (Hansen-2022 collection), limbic belt excluded, Set B (Glasser-decisive)<br>Spearman $\rho$ = -0.3585 | 4.3 | re-estimate with the belt removed (Set B (Glasser-decisive)) | vertex (32k fs_LR), n=57730 | vertex spin (N=1000, seed 0) | 0.034 |
| BigBrain profile skewness (pre-specified, histology), limbic belt excluded, Set B (Glasser-decisive)<br>Spearman $\rho$ = -0.2239 | 4.3 | re-estimate with the belt removed (Set B (Glasser-decisive)) | vertex (32k fs_LR), n=57730 | vertex spin (N=1000, seed 0) | 0.278 |
| MEG intrinsic timescale (int_area), limbic belt excluded, Set A (whole belt)<br>mean per-subject Spearman $\rho$ = -0.3531 | 4.3 | re-estimate with the belt removed (Set A (whole belt)) | subject (N=89) x vertex (4k fs_LR), n=89 subjects x 6242 vertices | hierarchical subject x vertex spin (N=1000, seed 0) | 0.066 |
| Intrinsic timescale (area under the ACF), adjusted for the aperiodic exponent<br>mean partial Spearman $\rho$ = -0.4361 | Annex F | model M1: 1 covariate(s) removed per subject | subject (N=89) x vertex (4k fs_LR), n=89 subjects | hierarchical subject x vertex spin (N=1000, seed 0) | 0.010 |
| Intrinsic timescale (area under the ACF), adjusted for all five band powers<br>mean partial Spearman $\rho$ = -0.3479 | Annex F | model M2: 5 covariate(s) removed per subject | subject (N=89) x vertex (4k fs_LR), n=89 subjects | hierarchical subject x vertex spin (N=1000, seed 0) | 0.002 |
| Intrinsic timescale (area under the ACF), adjusted for the aperiodic exponent and all five band powers<br>mean partial Spearman $\rho$ = -0.2844 | Annex F | model M3: 6 covariate(s) removed per subject | subject (N=89) x vertex (4k fs_LR), n=89 subjects | hierarchical subject x vertex spin (N=1000, seed 0) | 0.009 |
| Power-weighted spectral centroid, adjusted for the aperiodic exponent and all five band powers<br>mean partial Spearman $\rho$ = +0.2817 | Annex F | model M3: 6 covariate(s) removed per subject | subject (N=89) x vertex (4k fs_LR), n=89 subjects | hierarchical subject x vertex spin (N=1000, seed 0) | 0.010 |
| Slow/fast band-power ratio, adjusted for the aperiodic exponent and all five band powers<br>mean partial Spearman $\rho$ = -0.2551 | Annex F | model M3: 6 covariate(s) removed per subject | subject (N=89) x vertex (4k fs_LR), n=89 subjects | hierarchical subject x vertex spin (N=1000, seed 0) | 0.020 |
| Aperiodic exponent (fixed, 2-40 Hz), adjusted for the intrinsic timescale<br>mean partial Spearman $\rho$ = -0.0466 | Annex F | model R1: 1 covariate(s) removed per subject | subject (N=89) x vertex (4k fs_LR), n=89 subjects | hierarchical subject x vertex spin (N=1000, seed 0) | 0.817 |
| Aperiodic exponent (fixed, 2-40 Hz), adjusted for int_area, osc_delta, osc_theta, osc_alpha, osc_beta, osc_gamma1<br>mean partial Spearman $\rho$ = -0.0815 | Annex F | model R2: 6 covariate(s) removed per subject | subject (N=89) x vertex (4k fs_LR), n=89 subjects | hierarchical subject x vertex spin (N=1000, seed 0) | 0.501 |

| What was tested | § | The analysis it varies | Unit ($n_{\mathrm{obs}}$) | Null | $p_{\mathrm{raw}}$ |
|---|---|---|---|---|---|
| Short-range connectivity vs type-distance, aggregate level, model R0 coefficient per SD = -2.0321 | Annex G | offset log(n_i n_j), RR7's offset; distance covariate: none | type pair (7x7 off-diagonal), n=21 | type-map rotation (N=1000, seed 0), coefficient per SD | 0.060 |
| Short-range connectivity vs type-distance, aggregate level, model R1 coefficient per SD = -1.7182 | Annex G | offset log(pairs <= 80 mm); distance covariate: none | type pair (7x7 off-diagonal), n=21 | type-map rotation (N=1000, seed 0), coefficient per SD | 0.067 |
| Short-range connectivity vs type-distance, aggregate level, model R2 coefficient per SD = -2.1862 | Annex G | offset log(pairs <= 80 mm); distance covariate: contact area | type pair (7x7 off-diagonal), n=21 | type-map rotation (N=1000, seed 0), coefficient per SD | 0.021 |
| Short-range connectivity vs type-distance, aggregate level, model R3 coefficient per SD = -2.5036 | Annex G | offset log(pairs <= 80 mm); distance covariate: contact area + NN median (Euclidean) | type pair (7x7 off-diagonal), n=21 | type-map rotation (N=1000, seed 0), coefficient per SD | 0.012 |
| Short-range connectivity vs type-distance, aggregate level, model R1-40 coefficient per SD = -1.1268 | Annex G | offset log(pairs <= 40 mm); distance covariate: none; reported as a further variant | type pair (7x7 off-diagonal), n=21 | type-map rotation (N=1000, seed 0), coefficient per SD | 0.113 |
| Short-range connectivity vs type-distance, aggregate level, model R1-120 coefficient per SD = -2.0226 | Annex G | offset log(pairs <= 120 mm); distance covariate: none; reported as a further variant | type pair (7x7 off-diagonal), n=21 | type-map rotation (N=1000, seed 0), coefficient per SD | 0.058 |
| Short-range connectivity vs type-distance, aggregate level, model R3-geo coefficient per SD = -2.4361 | Annex G | offset log(pairs <= 80 mm); distance covariate: contact area + NN median (geodesic); reported as a further variant | type pair (7x7 off-diagonal), n=21 | type-map rotation (N=1000, seed 0), coefficient per SD | 0.013 |
| Short-range connectivity vs type-distance, bundle level, model R0 coefficient per SD = +0.0154 | Annex G | offset log(n_i n_j), RR7's offset; distance covariate: none | bundle x type pair, n=924 | type-map rotation (N=1000, seed 0), coefficient per SD | 0.900 |
| Short-range connectivity vs type-distance, bundle level, model R2 coefficient per SD = +0.3239 | Annex G | offset log(pairs <= 80 mm); distance covariate: contact area | bundle x type pair, n=924 | type-map rotation (N=1000, seed 0), coefficient per SD | 0.219 |
| Short-range connectivity vs type-distance, bundle level, model R3 coefficient per SD = +0.7736 | Annex G | offset log(pairs <= 80 mm); distance covariate: contact area + NN median (Euclidean) | bundle x type pair, n=924 | type-map rotation (N=1000, seed 0), coefficient per SD | 0.055 |
| Short-range connectivity vs type-distance, bundle level, model R1-40 coefficient per SD = +0.7236 | Annex G | offset log(pairs <= 40 mm); distance covariate: none; reported as a further variant | bundle x type pair, n=924 | type-map rotation (N=1000, seed 0), coefficient per SD | 0.145 |
| Short-range connectivity vs type-distance, bundle level, model R1-120 coefficient per SD = +0.0143 | Annex G | offset log(pairs <= 120 mm); distance covariate: none; reported as a further variant | bundle x type pair, n=924 | type-map rotation (N=1000, seed 0), coefficient per SD | 0.900 |
| Short-range connectivity vs type-distance, bundle level, model R3-geo coefficient per SD = +0.5225 | Annex G | offset log(pairs <= 80 mm); distance covariate: contact area + NN median (geodesic); reported as a further variant | bundle x type pair, n=924 | type-map rotation (N=1000, seed 0), coefficient per SD | 0.136 |
| Win fraction, holdout: fit LH, score RH win fraction = 0.5969 | 3.3 | disagreement subset: all; chance 0.4495 | vertex (32k fs_LR), n=13777 | vertex spin (N=1000, seed 0), win fraction recomputed per rotation | 0.001 |
| Win fraction, holdout: fit RH, score LH win fraction = 0.6046 | 3.3 | disagreement subset: all; chance 0.4516 | vertex (32k fs_LR), n=13521 | vertex spin (N=1000, seed 0), win fraction recomputed per rotation | 0.001 |

| What was tested | § | The analysis it varies | Unit ($n_{\text{obs}}$) | Null | $p_{\text{raw}}$ |
|---|---|---|---|---|---|
| Win fraction, holdout: fit LH, score RH<br>win fraction = 0.7289 | 3.3 | disagreement subset: off2; chance 0.4261 | vertex (32k fs_LR), n=2163 | vertex spin (N=1000, seed 0), win fraction recomputed per rotation | 0.002 |
| Win fraction, holdout: fit RH, score LH<br>win fraction = 0.7563 | 3.3 | disagreement subset: off2; chance 0.4287 | vertex (32k fs_LR), n=1967 | vertex spin (N=1000, seed 0), win fraction recomputed per rotation | 0.001 |
| Win fraction, areal interior (> 5 hops)<br>win fraction = 0.5872 | 3.3 | disagreement subset: all; chance 0.4485 | vertex (32k fs_LR), n=4844 | vertex spin (N=1000, seed 0), win fraction recomputed per rotation | 0.007 |
| Win fraction, areal border band (<= 5 hops)<br>win fraction = 0.6040 | 3.3 | disagreement subset: all; chance 0.4592 | vertex (32k fs_LR), n=22454 | vertex spin (N=1000, seed 0), win fraction recomputed per rotation | 0.001 |
| Win fraction, areal interior (> 5 hops)<br>win fraction = 0.8194 | 3.3 | disagreement subset: off2; chance 0.4282 | vertex (32k fs_LR), n=24 | vertex spin (N=1000, seed 0), win fraction recomputed per rotation | 0.001 |
| Win fraction, areal border band (<= 5 hops)<br>win fraction = 0.7428 | 3.3 | disagreement subset: off2; chance 0.4335 | vertex (32k fs_LR), n=4106 | vertex spin (N=1000, seed 0), win fraction recomputed per rotation | 0.001 |
| T1w/T2w myelin vs the myelin septile<br>Spearman $\rho$ = +0.9897 | Tables S6 / S7 | granularity-matched challenger partition (vertex level) | vertex (32k fs_LR) | vertex spin (N=1000, seed 0) | 0.001 |
| T1w/T2w myelin vs the gradient septile<br>Spearman $\rho$ = -0.6150 | Tables S6 / S7 | granularity-matched challenger partition (vertex level) | vertex (32k fs_LR) | vertex spin (N=1000, seed 0) | 0.002 |
| T1w/T2w myelin vs the BigBrain Hist-G1 septile<br>Spearman $\rho$ = +0.3448 | Tables S6 / S7 | granularity-matched challenger partition (vertex level) | vertex (32k fs_LR) | vertex spin (N=1000, seed 0) | 0.094 |
| Cortical thickness vs the myelin septile<br>Spearman $\rho$ = -0.5585 | Tables S6 / S7 | granularity-matched challenger partition (vertex level) | vertex (32k fs_LR) | vertex spin (N=1000, seed 0) | 0.002 |
| Cortical thickness vs the gradient septile<br>Spearman $\rho$ = +0.3893 | Tables S6 / S7 | granularity-matched challenger partition (vertex level) | vertex (32k fs_LR) | vertex spin (N=1000, seed 0) | 0.030 |
| Cortical thickness vs the BigBrain Hist-G1 septile<br>Spearman $\rho$ = -0.4043 | Tables S6 / S7 | granularity-matched challenger partition (vertex level) | vertex (32k fs_LR) | vertex spin (N=1000, seed 0) | 0.009 |
| Intrinsic timescale vs the myelin septile<br>Spearman $\rho$ = -0.6101 | Tables S6 / S7 | granularity-matched challenger partition (vertex level) | vertex (32k fs_LR) | vertex spin (N=1000, seed 0) | 0.011 |
| Intrinsic timescale vs the gradient septile<br>Spearman $\rho$ = +0.5279 | Tables S6 / S7 | granularity-matched challenger partition (vertex level) | vertex (32k fs_LR) | vertex spin (N=1000, seed 0) | 0.009 |
| Intrinsic timescale vs the BigBrain Hist-G1 septile<br>Spearman $\rho$ = -0.3779 | Tables S6 / S7 | granularity-matched challenger partition (vertex level) | vertex (32k fs_LR) | vertex spin (N=1000, seed 0) | 0.333 |
| Delta power (0.5-4 Hz) vs the myelin septile<br>Spearman $\rho$ = -0.6077 | Tables S6 / S7 | granularity-matched challenger partition (vertex level) | vertex (32k fs_LR) | vertex spin (N=1000, seed 0) | 0.008 |
| Delta power (0.5-4 Hz) vs the gradient septile<br>Spearman $\rho$ = +0.4830 | Tables S6 / S7 | granularity-matched challenger partition (vertex level) | vertex (32k fs_LR) | vertex spin (N=1000, seed 0) | 0.016 |
| Delta power (0.5-4 Hz) vs the BigBrain Hist-G1 septile<br>Spearman $\rho$ = -0.4555 | Tables S6 / S7 | granularity-matched challenger partition (vertex level) | vertex (32k fs_LR) | vertex spin (N=1000, seed 0) | 0.225 |

| What was tested | § | The analysis it varies | Unit ($n_{\mathrm{obs}}$) | Null | $p_{\mathrm{raw}}$ |
|---|---|---|---|---|---|
| Theta power (4-8 Hz) vs the myelin septile<br>Spearman $\rho$ = -0.3850 | Tables S6 / S7 | granularity-matched challenger partition (vertex level) | vertex (32k fs_LR) | vertex spin (N=1000, seed 0) | 0.283 |
| Theta power (4-8 Hz) vs the gradient septile<br>Spearman $\rho$ = +0.1757 | Tables S6 / S7 | granularity-matched challenger partition (vertex level) | vertex (32k fs_LR) | vertex spin (N=1000, seed 0) | 0.555 |
| Theta power (4-8 Hz) vs the BigBrain Hist-G1 septile<br>Spearman $\rho$ = -0.5768 | Tables S6 / S7 | granularity-matched challenger partition (vertex level) | vertex (32k fs_LR) | vertex spin (N=1000, seed 0) | 0.040 |
| Alpha power (8-12 Hz) vs the myelin septile<br>Spearman $\rho$ = +0.5741 | Tables S6 / S7 | granularity-matched challenger partition (vertex level) | vertex (32k fs_LR) | vertex spin (N=1000, seed 0) | 0.025 |
| Alpha power (8-12 Hz) vs the gradient septile<br>Spearman $\rho$ = -0.3570 | Tables S6 / S7 | granularity-matched challenger partition (vertex level) | vertex (32k fs_LR) | vertex spin (N=1000, seed 0) | 0.163 |
| Alpha power (8-12 Hz) vs the BigBrain Hist-G1 septile<br>Spearman $\rho$ = +0.6043 | Tables S6 / S7 | granularity-matched challenger partition (vertex level) | vertex (32k fs_LR) | vertex spin (N=1000, seed 0) | 0.021 |
| Beta power (12-30 Hz) vs the myelin septile<br>Spearman $\rho$ = +0.1601 | Tables S6 / S7 | granularity-matched challenger partition (vertex level) | vertex (32k fs_LR) | vertex spin (N=1000, seed 0) | 0.688 |
| Beta power (12-30 Hz) vs the gradient septile<br>Spearman $\rho$ = -0.2906 | Tables S6 / S7 | granularity-matched challenger partition (vertex level) | vertex (32k fs_LR) | vertex spin (N=1000, seed 0) | 0.273 |
| Beta power (12-30 Hz) vs the BigBrain Hist-G1 septile<br>Spearman $\rho$ = -0.2105 | Tables S6 / S7 | granularity-matched challenger partition (vertex level) | vertex (32k fs_LR) | vertex spin (N=1000, seed 0) | 0.648 |
| Gamma power (30-60 Hz) vs the myelin septile<br>Spearman $\rho$ = -0.5200 | Tables S6 / S7 | granularity-matched challenger partition (vertex level) | vertex (32k fs_LR) | vertex spin (N=1000, seed 0) | 0.031 |
| Gamma power (30-60 Hz) vs the gradient septile<br>Spearman $\rho$ = +0.2751 | Tables S6 / S7 | granularity-matched challenger partition (vertex level) | vertex (32k fs_LR) | vertex spin (N=1000, seed 0) | 0.272 |
| Gamma power (30-60 Hz) vs the BigBrain Hist-G1 septile<br>Spearman $\rho$ = -0.5279 | Tables S6 / S7 | granularity-matched challenger partition (vertex level) | vertex (32k fs_LR) | vertex spin (N=1000, seed 0) | 0.051 |
| Principal functional gradient vs the myelin septile<br>Spearman $\rho$ = -0.6259 | Tables S6 / S7 | granularity-matched challenger partition (vertex level) | vertex (32k fs_LR) | vertex spin (N=1000, seed 0) | 0.001 |
| Principal functional gradient vs the gradient septile<br>Spearman $\rho$ = +0.9897 | Tables S6 / S7 | granularity-matched challenger partition (vertex level) | vertex (32k fs_LR) | vertex spin (N=1000, seed 0) | 0.001 |
| Principal functional gradient vs the BigBrain Hist-G1 septile<br>Spearman $\rho$ = -0.2322 | Tables S6 / S7 | granularity-matched challenger partition (vertex level) | vertex (32k fs_LR) | vertex spin (N=1000, seed 0) | 0.193 |
| gene_pc1 vs the myelin septile<br>Spearman $\rho$ = +0.8412 | Tables S6 / S7 | granularity-matched challenger partition (vertex level) | vertex (32k fs_LR) | vertex spin (N=1000, seed 0) | 0.001 |
| gene_pc1 vs the gradient septile<br>Spearman $\rho$ = -0.5860 | Tables S6 / S7 | granularity-matched challenger partition (vertex level) | vertex (32k fs_LR) | vertex spin (N=1000, seed 0) | 0.002 |
| gene_pc1 vs the BigBrain Hist-G1 septile<br>Spearman $\rho$ = +0.4409 | Tables S6 / S7 | granularity-matched challenger partition (vertex level) | vertex (32k fs_LR) | vertex spin (N=1000, seed 0) | 0.089 |

| What was tested | § | The analysis it varies | Unit ($n_{\mathrm{obs}}$) | Null | $p_{\mathrm{raw}}$ |
|---|---|---|---|---|---|
| histg1 vs the myelin septile Spearman $\rho$ = +0.3509 | Tables S6 / S7 | granularity-matched challenger partition (vertex level) | vertex (32k fs_LR) | vertex spin (N=1000, seed 0) | 0.097 |
| histg1 vs the gradient septile Spearman $\rho$ = -0.2346 | Tables S6 / S7 | granularity-matched challenger partition (vertex level) | vertex (32k fs_LR) | vertex spin (N=1000, seed 0) | 0.193 |
| histg1 vs the BigBrain Hist-G1 septile Spearman $\rho$ = +0.9897 | Tables S6 / S7 | granularity-matched challenger partition (vertex level) | vertex (32k fs_LR) | vertex spin (N=1000, seed 0) | 0.001 |
| receptor_pc1 vs the myelin septile Spearman $\rho$ = -0.5953 | Tables S6 / S7 | granularity-matched challenger partition (vertex level) | vertex (32k fs_LR) | vertex spin (N=1000, seed 0) | 0.001 |
| receptor_pc1 vs the gradient septile Spearman $\rho$ = +0.4667 | Tables S6 / S7 | granularity-matched challenger partition (vertex level) | vertex (32k fs_LR) | vertex spin (N=1000, seed 0) | 0.007 |
| receptor_pc1 vs the BigBrain Hist-G1 septile Spearman $\rho$ = -0.2496 | Tables S6 / S7 | granularity-matched challenger partition (vertex level) | vertex (32k fs_LR) | vertex spin (N=1000, seed 0) | 0.218 |

Constantin von Economo and Georg N Koskinas. *Die Cytoarchitektonik der Hirnrinde des erwachsenen Menschen*. Springer, Vienna, 1925.

Ting Xu, Karl-Heinz Nenning, Ernst Schwartz, Seok-Jun Hong, Joshua T. Vogelstein, Alexandros Goulas, Damien A. Fair, Charles E. Schroeder, Daniel S. Margulies, Jonathan Smallwood, Michael P. Milham, and Georg Langs. Cross-species functional alignment reveals evolutionary hierarchy within the connectome. *NeuroImage*, 223:117346, 2020.

Fang-Cheng Yeh, Sandip Panesar, David Fernandes, Antonio Meola, Masanori Yoshino, Juan C Fernandez-Miranda, Jean M Vettel, and Timothy Verstynen. Population-averaged atlas of the macroscale human structural connectome and its network topology. *NeuroImage*, 178:57–68, 2018.

Josefa Zaldívar-Díez and Miguel Ángel García-Cabezas. Cortical type: the basis for a modern neurodevelopmental synthesis (ontogeny, phylogeny, synaptic connectivity) of the human cerebral cortex. *Journal of Comparative Neurology*, 534(2):e70143, 2026. doi: 10.1002/cne.70143.
